# Intact nucleosomal context enables chromodomain reader MRG15 to distinguish H3K36me3 from -me2

**DOI:** 10.1101/2020.04.30.070136

**Authors:** Sarah Faulkner, Antony M. Couturier, Brian Josephson, Tom Watts, Benjamin G. Davis, Fumiko Esashi

## Abstract

A wealth of *in vivo* evidence demonstrates the physiological importance of histone H3 trimethylation at lysine 36 (H3K36me3), to which chromodomain-containing proteins, such as MRG15, bind preferentially compared to their dimethyl (H3K36me2) counterparts. However, *in vitro* studies using isolated H3 peptides have failed to recapitulate a causal interaction. Here, we show that MRG15 can clearly discriminate between synthetic, fully intact model nucleosomes containing H3K36me2 and H3K36me3. MRG15 docking studies, along with experimental observations and nucleosome structure analysis suggest a model where the H3K36 side chain is sequestered in intact nucleosomes via a hydrogen bonding interaction with the DNA backbone, which is abrogated when the third methyl group is added to form H3K36me3. Hence, this mechanism provides a ‘methyl-switch’ for contextdependent reader selectivity. These results highlight the importance of such intra-chromatin interactions in understanding epigenetic regulation, a feature which is absent in commonly-used peptide or histone-only models.

## Introduction

The N-terminal tail of histone H3 contains numerous lysine residues which are known to bear a range of post-translational modifications (PTMs) which regulate the structure and interaction partners of chromatin^1^. For some PTMs such as acetylation, the chromatin structure is known to be directly modulated by altering the strength of interaction between DNA and histone octamer^2^. However, other PTMs are thought to effect change in the nucleus by recruiting specific so-called ‘reader’ proteins that can then form multimeric complexes to carry out a range of functions. Perhaps the most well characterised PTM in this latter category is lysine methylation^3^.

Each lysine can bear up to three methyl groups, and in some cases each methylation state of a single residue has a unique function^4,5^. A striking example is lysine 36 of histone H3 (H3K36), which has a dedicated methyltransferase enzyme SETD2 (SET Domain containing 2), solely responsible for converting dimethylated H3K36 (H3K36me2) to trimethylated H3K36 (H3K36me3)^6^. This dedicated, conserved role suggests a critical, but as yet not fully understood, role for this subtle di- to tri- ‘methyl switch’. Furthermore, we can infer from *in vivo* observations that this ‘switch’ to H3K36me3 is functionally central to both genome stability and development; mutations to SETD2 have been implicated in causing a range of cancers^7,8^. In a laboratory setting, homozygous disruption of SETD2 is found to be embryonically lethal^9^.

One associated methyllysine reader known to maintain genome stability is the chromodomain protein MRG15 (MORF Related Gene on chromosome 15)^10,11^. MRG15 recognises methylated H3K36, and strong *in vivo* evidence suggests a clear preference for H3K36me3. Specifically, MRG15 has been shown to localise to the same sites as SETD2 and H3K36me3, and increasing or decreasing H3K36me3, by upregulating SETD2 or KDM4A (Lysine-specific Demethylase 4A) respectively, has a corresponding correlated impact on MRG15-mediated functions^10,12,13^. Notably, however, this preference is not recapitulated by *in vitro* experiments with peptide fragments from H3; to date these have shown no discrimination between di- and trimethyl states^14^. Moreover, to our knowledge, no structural basis for this MRG15-mediated interaction that would support such discrimination exists.

Here, through the synthesis and investigation of increasingly complex and context-dependent ligands for MRG15, we reveal that discrimination of H3K36me3 from -me2 requires, as a minimum, the presence of the surrounding intact nucleosomal environment. This suggests that some methyllysine readers such as the MRG15 chromodomain achieve selectivity via a ‘reading in context’ mechanism rather than on the basis of the methylation mark alone.

## Results

With the aim of assessing the impact of H3K36 methylation, we generated full-length histones containing 100% mark occupancy of H3K36me2 or H3K36me3. These were produced using chemical, carbon radical (C•) mediated C-C bond formation to precisely chemically ‘edit’ recombinantly expressed H3 with the required modified, methylated side chains (**Fig. 1A**)^15^. Specifically, we first generated a *X. laevis* H3K36C mutant using site-directed mutagenesis, expressed the histone recombinantly in *E. coli* and purified as previously reported^16^. The highly conserved nature of histone sequences has meant that results from *X. laevis* histones have proved broadly and successfully applicable to other species^17–19^. We began with the fully functional H3C110A mutant (denoted here as ‘H3 WT’) commonly used in biophysical studies^20^. As cysteine is not present anywhere else in this H3 variant, it provided us with a reactive handle to target for further transformation; first, through a *bis*-alkylation-elimination reaction to dehydroalanine (Dha), a powerful radical acceptor.^15^ This could then be subsequently reacted with appropriate sources of radicals Me_2_NCH_2_CH_2_CH_2_• or Me_3_NCH_2_CH_2_CH_2_• to append our side-chains of interest (**Fig. 1B**) as a post-translational mode of mutagenesis that allows access to relevant PTMs. However, despite demonstrated 100% occupancy of these PTM marks in H3 protein (see **Methods and Appendix**), GST-MRG15 fusion protein bound similar quantities of H3K36me2 and -me3, indicating no significant discrimination between them, even when found in full length histone at the appropriate site 36 (**Fig. 1C**).

**Figure 1:**
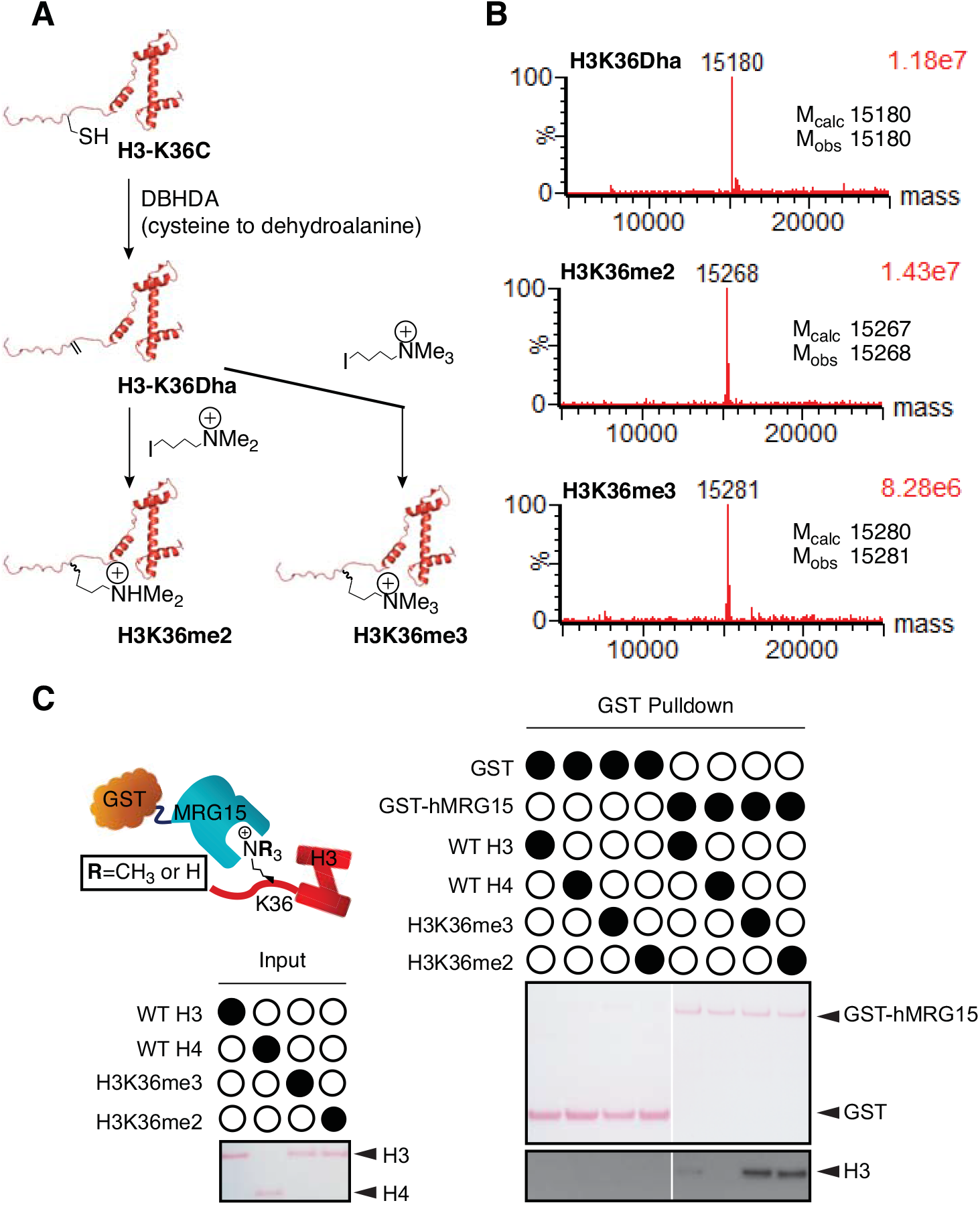
MRG15 interaction with synthetic H3 bearing K36me2 or K36me3. **A,** Scheme for post-translational mutagenesis to generate H3K36me2 and -me3 via 2,5 dibromohexanediamide (DBHDA)-mediated conversion of K36C cysteine to dehydroalanine (Dha), followed by radical-mediated post-translational mutagenesis to append the appropriate alkyl chain to Dha. **B,** LCMS-ESI+ characterization of H3K36Dha, -me2 and -me3. Mcalc; calculated mass, Mobs; observed mass. Peak intensity (%) is shown on the y axis, mass (Da) is shown on the x axis. Total ion count is shown in the top right corner. **C,** Pulldown using GST-MRG15 immobilised on GST Sepharose beads with full length histone H3 variants (WT, K36me2 and K36me3) and H4 (WT). Input panel and GST panel of GST pulldown are stained with Ponceau S, H3 panel is from Western blot using α-histone H3 as the primary antibody.

Upon solving the 3D X-ray crystal structure of the MRG15 chromodomain, Zhang et al suggested that Tyr26, Tyr46 and Trp49 might form an aromatic, hydrophobic pocket to bind methylated H3K36 in a manner that is analogous to others, such as the Tyr21, Trp42, Phe45 triad of mouse HP1α, which binds methyl H3K9^14,21^; this model is gaining increasing traction, as exemplified by the recent cryo-EM structure of H3K36_C_me3 nucleosome in complex with the closely related PWWP domain^22^. Our independent structural alignment and docking studies further supported this hypothesis for the monomeric MRG15 chromodomain (**Fig. 2A and B**). HP1 binding to methylated lysine has been extensively studied, and is thought to arise from two components: namely, a small stabilising contribution from hydrophobic interactions between methyl groups and aromatic residues, and a larger contribution from cation-pi interactions between the positive charge of N-methylated lysine and the quadrupole moment of tryptophan^23^. However, in this model, much of the stabilization afforded to -N(+)me3 also occurs with -N(+)Hme2, which has similar charge and hydrophobicity^24^. How then, do we reconcile the specific MRG15-mediated function of H3K36me3 over -me2 found in current *in vivo* observations, with the apparent lack of a molecular mechanism for clear discrimination between the two methylation states, especially when discrimination is also not present using H3K36me2/H3K36me3 histones alone?

**Figure 2:**
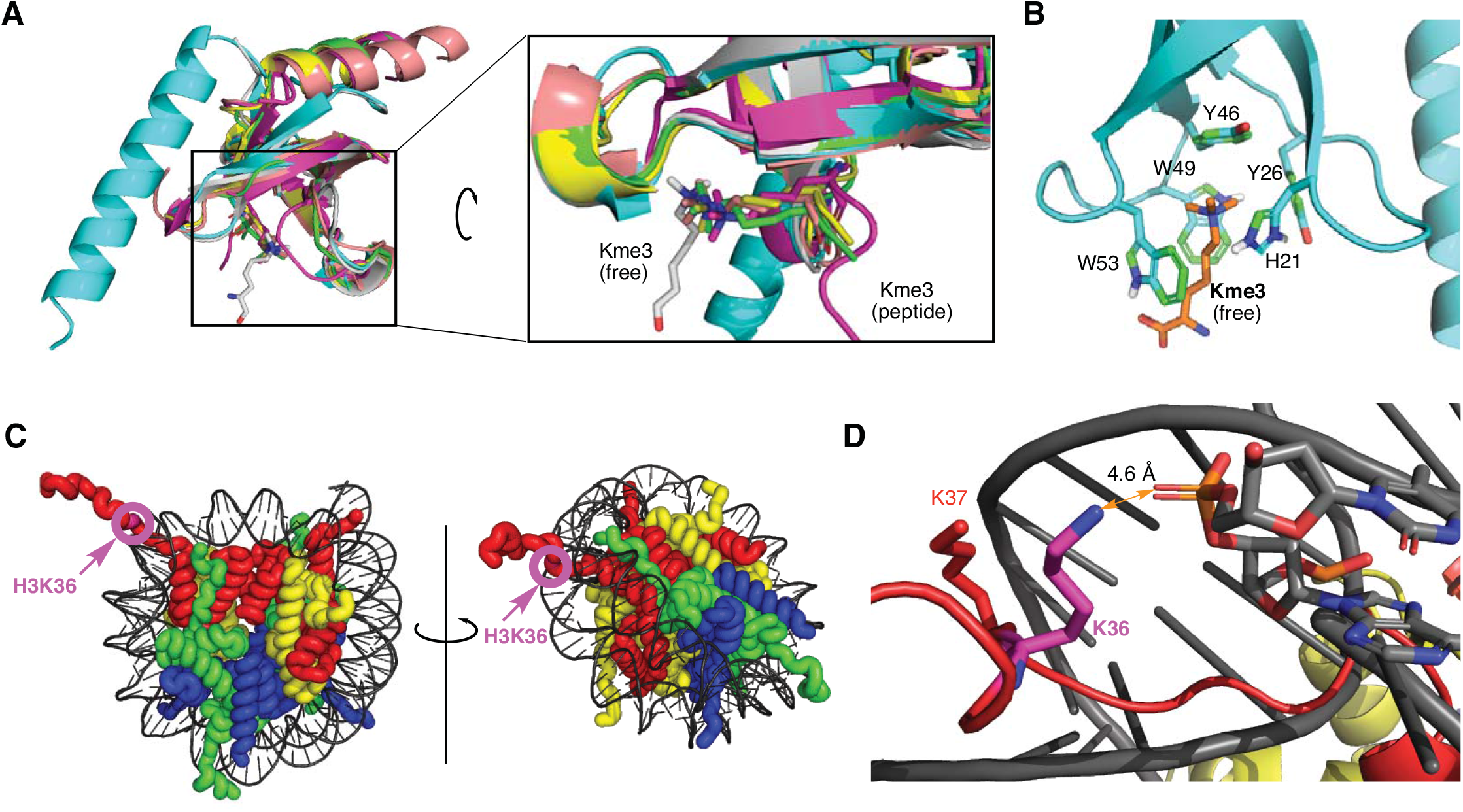
H3K36me3 in complex with chromodomains and in intact nucleosome. **A,** Overlay of chromodomains crystal structure (PDB) from MRG15 (cyan, 2F5K), HP-1 with H3K9me3 tail peptide (green, 1KNE), Chp1 with H3K9me3 tail peptide (pink, 2RSN), Chp1 with H3K9me3 tail peptide (yellow, 3G71), MPP8 with H3K9me3 tail peptide (salmon, 3QO2), MRG2 with free Kme3 amino acid (grey, 4PL6). Curved arrow represents 180° rotation about the central y axis. **B,** Docking structure of free Kme3 amino acid with MRG15 chromodomain (2F5K). Key aromatic residues are highlighted. **C,** Nucleosome crystal structure (1AOI) with H3K36 highlighted. Histone H3 is shown in red, H4 in yellow, H2A in green and H2B in blue. DNA is shown in grey. **D,** Zoomed view of H3 tail and DNA (1AOI). DNA backbone is shown in grey, H3 in red except for highlighted H3K36 which is shown in magenta. Heteroatoms are colour coded as follows; P; orange, O; red, N; blue.

K36 is unique among the commonly methylated H3 lysines in that it is located at the base of the H3 N-terminal tail, directly at the interface between histone octamer and the DNA phosphodiester backbone (**Fig. 2C and D**)^25^. We reasoned that likely K36-to-DNA backbone interactions at such proximity might therefore modulate MRG15 binding, effectively competing with MRG15 to envelop the residue. Such competition would potentially allow additional modes of discrimination. Previous *in vitro* studies, which used only peptides or isolated H3, would be unable to capture this ‘shielding’, competitive effect^14^.

We therefore hypothesised that discrimination between methylation states by MRG15 may be critically context-dependent, plausibly requiring an intact nucleosomal architecture. With the aim of reconciling our *in vivo* and *in vitro* observations, the exact same full length H3K36me2 and -me3 proteins generated by chemical editing / post-translational mutagenesis above were assembled into intact nucleosomes bearing suitable biotinylated DNA fragments for pulldown interaction studies (**Fig. 3A**). In brief, histone octamers containing H3K36me2 or -me3 were first assembled using standard refolding protocols^26^. Interestingly, in size exclusion chromatography (SEC) traces during intermediate stages, we noted that assembly equilibria were perturbed as the number of methyl groups on H3K36 increased. For example, the ratio between octamer and H3_2_-H4_2_ tetramer peaks decreased under identical conditions, suggesting that the addition of each methyl group already caused a small but significant decrease in core histone assembly stability (**Fig. 3B**).

**Figure 3:**
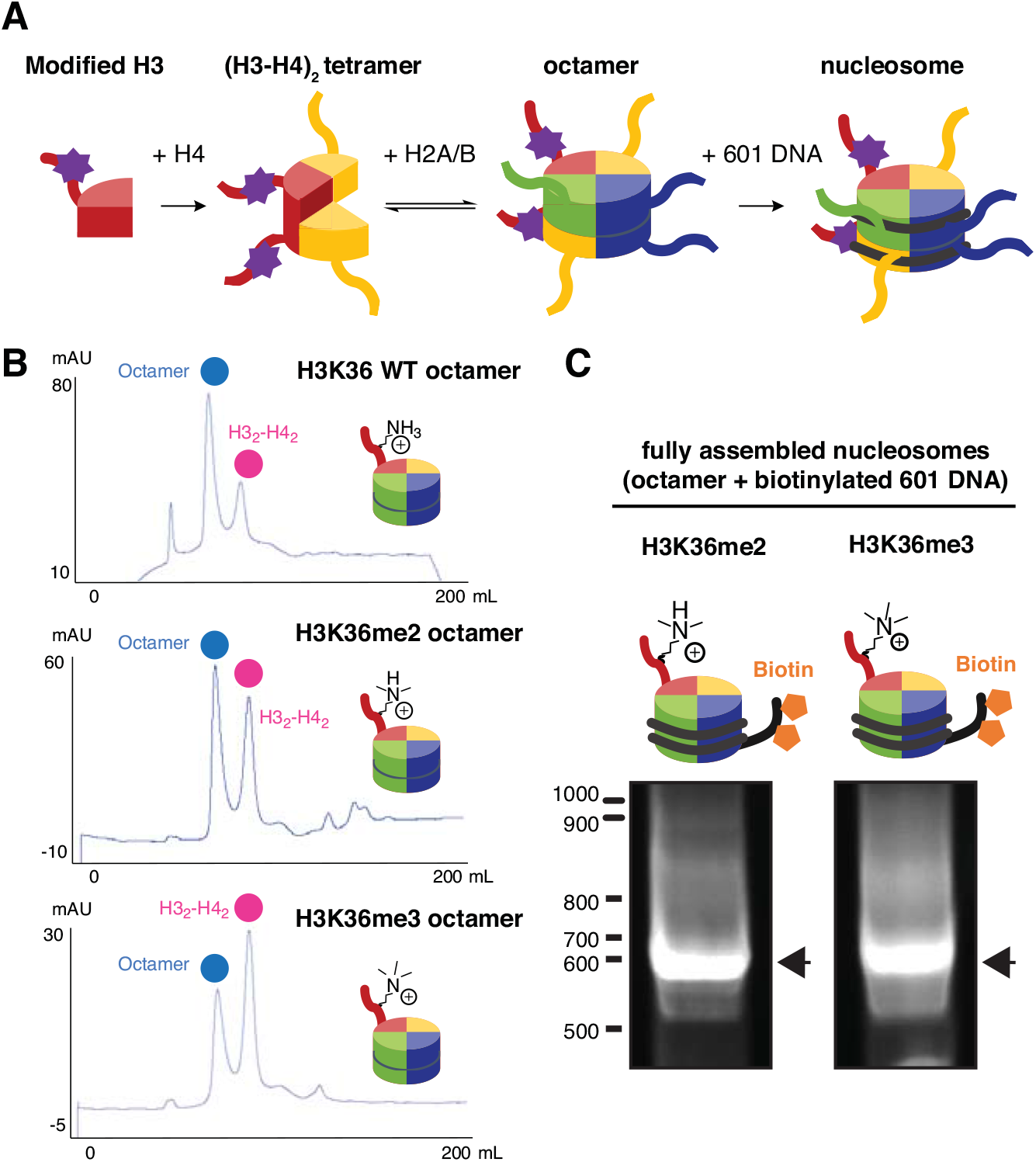
Assembly of intact nucleosome bearing synthetic H3K36me2 or -me3. **A,** Overall scheme for assembly of precisely methylated nucleosomes. Histone H3 is shown in red, H4 in yellow, H2A in green and H2B in blue. DNA is shown in grey. The synthetically methylated H3K36 is highlighted as a purple star. **B,** Size-Exclusion Chromatography (SEC) UV traces for octamer assembly with H3K36 WT, K36me2 and K36me3. UV absorbance (mAU) is shown on the y axis, elution volume (mL) on the x axis. **C,** Intact, fully assembled nucleosome bands (arrows), detected from native TBE gel. DNA molecular size (bp) is indicated on the left. Biotin is represented by orange pentagons.

With each of the H3 WT, H3K36me2 and H3K36me3-containing octamers in hand, we then reconstituted nucleosomes according to literature protocols using a biotinylated 186 base-pair (bp) DNA fragment bearing the 147 bp Widom 601 positioning sequence, which binds the histone octamer with high affinity^26^ (**Fig. 3C**), used here to generate a [39-N-0] positional configuration (where N is the nucleosome 147bp wrap site). Our in-house nucleosomes therefore harboured 39 bp of ‘spacer’ DNA region between assembled nucleosome and the affinity resin used during pulldown experiment. Denaturing SDS-PAGE confirmed the expected histone composition for all nucleosomes (**Supplemental Fig. S1**).

These precisely methylated nucleosomes were then tested in their binding to MRG15. Briefly, we immobilised them via their 186 bp biotinylated DNA onto magnetic streptavidin-coated beads and incubated them with MRG15 for several hours in the presence of salmon sperm DNA to outcompete non-specific binding, so that the distinct interaction between nucleosomes and MRG15 was allowed to reach equilibrium. The beads were thoroughly washed to remove all unbound protein, and the remaining bound MRG15 was visualised by Western blot. In contrast to previous results with peptides or full-length H3 (**Fig. 1C**) we observed a clear preference for -me3 nucleosomes over -me2 (**Supplemental Fig. S2A**). The MRG15 W53A mutant, in which a key residue of MRG15’s chromodomain aromatic triad is removed, exhibited loss of all binding in comparison to the WT, confirming the essential role of the chromodomain as opposed to the MRG domain (**Fig. 4A**) within MRG15. Notably, the observed binding strength and selectivity of MRG15 was dependent on ionic strength; use of 150 mM KCl increased both (**Fig. 4B**). Additionally, the type of countercation also modulated the interaction: a KCl-based buffer was used for pulldowns since prior measurements of salt concentrations in the nucleus, suggest that K^+^ is more abundant than other cations such as Na^+^^27,28^. Correspondingly, we found that the use of NaCl at 150 mM reduced both binding and selectivity (**Supplemental Fig. S2B**).

**Figure 4:**
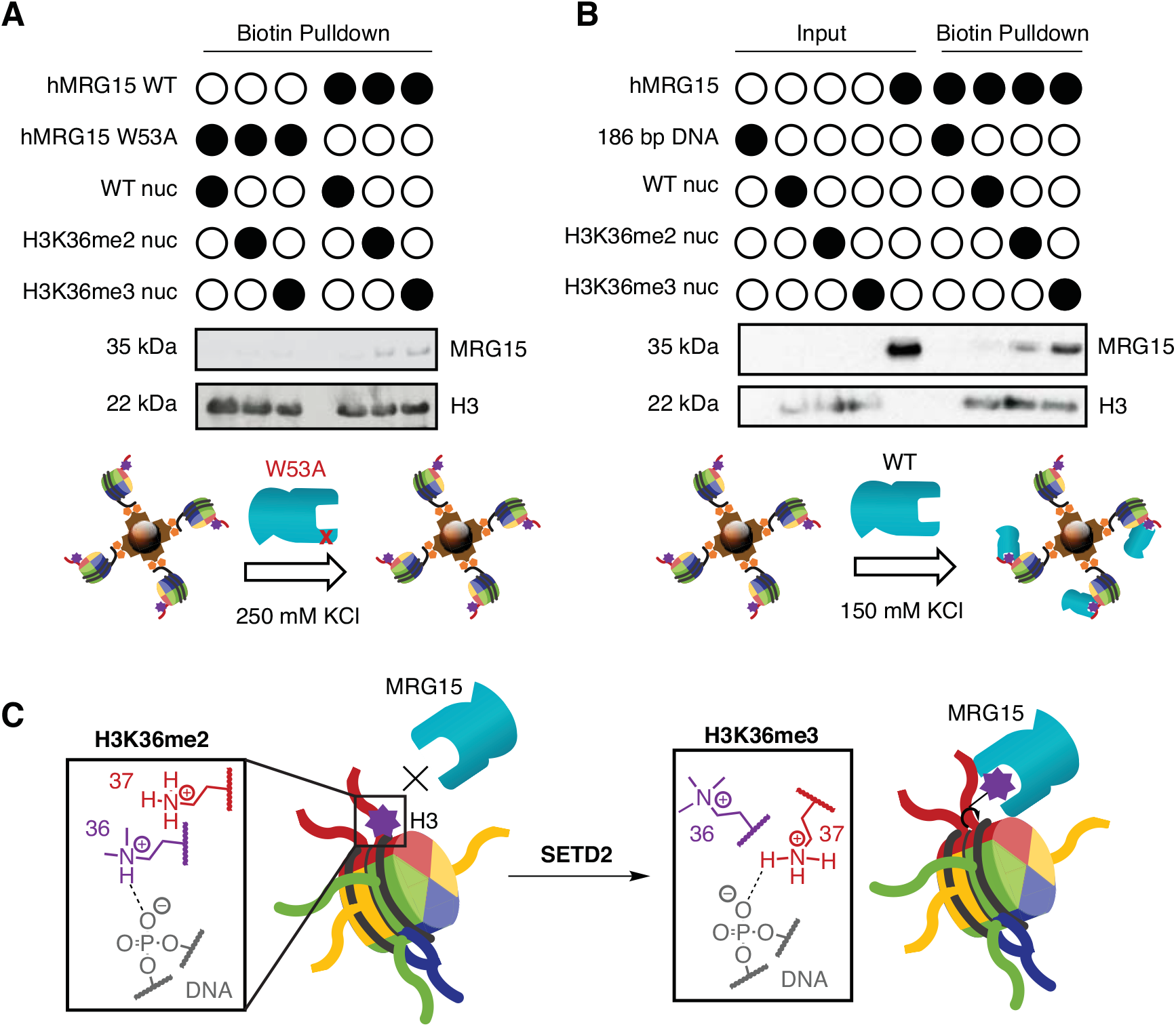
MRG15 interaction with intact nucleosomes bearing H3K36me2 or -me3. **A,** Western blot analysis of the biotin pulldown performed with the His-tagged WT MRG15 or W53A mutant using α-MRG15 (top panel) and α-histone H3 primary (bottom panel) antibodies. B, Western blot analysis of the pulldown performed with WT His-tagged MRG15 using α-MRG15 (top panel) and α-histone H3 (bottom panel) primary antibodies. C, Model for MRG15 discrimination between H3K36me2 and -me3 in intact nucleosomal context. Histone H3 is shown in red, H4 in yellow, H2A in green and H2B in blue. DNA is shown in grey. The methylated H3K36 is highlighted as a purple star.

## Discussion

Considering all these observations collectively, we propose the following model to explain MRG15 discrimination between di- and trimethylated nucleosomes. From 3D structural analysis, H3K36 is well positioned to form a context-driven hydrogen bond between its side chain -Nε–H and phosphodiester DNA backbone. In this state, H3K36 is shielded and inaccessible to MRG15. With a predicted pKa_H_ of 9.40 (ProPKA^29^), we expect this residue to exist primarily in a suitably protonated ammonium –NH_3_+ state (**Fig. 1A**) in K36, K36me and also K36me2, as up two of three hydrogens are replaced by methyl groups without disrupting bonding to phosphate oxygen. However, the addition of a third methyl to form K36me3 destroys the residue’s capacity for this hydrogen bond. We propose that this broken interaction, (perhaps also with the additional bulk and hydrophobicity of the trimethyl state), causes a conformational shift in the H3 tail where K36me3 becomes exposed, and therefore accessible to MRG15 (**Fig. 4C**).

Our data also suggest that a mechanism of selectivity is dependent not only on proper context but also on interactions modulated by charge density and ion type. In general, cations are known to abrogate interactions between DNA and octamer as they ion pair to the anionic DNA backbone; Na^+^ is known to do this more effectively than K^+30^. Moreover, ion type and ionic strength can modulate cooperativity and strength of hydrogen bonds^31^. Therefore, it stands to reason that discrimination by MRG15 may critically involve different modes of DNA-to-octamer interactions modulated by methylation states.

In this context, we also extended our analyses to assess interactions between MRG15 and other putative substrates. Commercially-available nucleosomes, prepared through multiple proprietary methods^32^ are available with H3K36me2 or -me3 (EpiCypher, Inc, Catalog No. 16-0319, Lot No. 19212004-20 for H3K36me2; Catalog No. 16-0320, Lot No. 18348003-20 for H3K36me3). These are assembled from recombinant human histones wrapped instead with shorter 147 bp 601 positioning sequence DNA attached directly to a 5’-biotin-triethyleneglycol (TEG) group^33,34^. In our hands, these nucleosomes lacking spacer DNA showed little preference for MRG15 interaction with H3K36me3 containing nucleosome compared to its H3K36me2 counterpart in biotin pulldown experiments (**Supplemental Fig. S3**).

Given that the histone modification methods of EpiCypher nucleosomes are proprietary, however, conclusions about the nature of this modest discrimination can only be inconclusive and we are unable to discount the possibility that differences arise due to differing methods in nucleosome synthesis and generation. Indeed, similarly prepared nucleosomes have previously allowed methylation states at H3K4, H3K9, H2K27, H3K36 and H4K20 to be distinguished by PTM specific antibodies and in other interaction systems^2,35-37^. Nonetheless, one of the known differences between these nucleosome types is the presence of a DNA spacer motif of 39 base pairs (i.e. 39-N-0 vs. 0-N-0), the estimated length of which is ~ 13 nm, which may explain these differences in MRG15 binding. We speculate that this spacer DNA would be particularly important given the tight positioning of H3K36 in proximity to the DNA at the entry and exit positions on the nucleosome; this is the termini of the minimum 147 bp 601 sequence, where directly appended biotin-TEG would need to be affixed to the protein/Dynabead format needed for assay. By contrast, we anticipate the additional spacer 39 bp DNA in discriminating nucleosomes would allow sufficient room for effective approach of 43.5 kDa full-length MRG15. Notably, unlike short ‘reader’ domains (for example the 8.5 kDa PWWP domains which also bind H3K36me3-containing nucleosomes^22^) or these domains fused with well-characterized epitope tags up to ~26 Da that have been used successfully with the directly-appended 147 bp TEG-biotin nucleosomes^36,37^, the full length MRG15 would need to bind as its 87 kDa homodimer, approaching the nucleosome in correct orientation. Indeed, while interaction studies on free 147 bp H3K36_C_me3-containing nucleosomes using methods such as NMR have previously proven successful^38–40^, additional linker DNA is typically necessary when the nucleosome is immobilised^22,41–44^. This further emphasizes that contextual caveats should be noted when assessing interactions between native ‘reader’ proteins (e.g. full length vs domain) and nucleosomes.

The mechanistic details of SETD2’s interplay with other key chromatin regulators is currently a topic of great interest^41^. Our finding presented in this study leads us to speculate that the buried nature of H3K36me2 may also have implications for the mode of SETD2 methylation, as it might be difficult for SETD2 to access its substrate when it is part of a whole nucleosome. SETD2 binds to the C-terminal domain tail of RNA Polymerase II (RNAPII), and has previously been assumed to methylate H3K36me2 in nucleosomes which have re-assembled after the polymerase has passed through them during transcription^45,46^. However, previous studies have shown that under conditions of moderate transcription, RNAPII is able to transcribe through nucleosomes by displacing only H2A-H2B dimers, leaving the H32-H42 tetramer intact^47^. This H32-H42 tetramer-only state is expected to leave H3K36 more exposed. Hence, it is conceivable that SETD2 may take advantage of this and methylate H3K36me2 concurrently with RNAPII transcription (**Supplemental Fig. S4**). Indeed, our observations from the octamer assembly suggest that H2A-H2B tetramers might dissociate more readily from H3K36me2-containing H32-H42 tetramers than the unmodified state, facilitating such a mechanism.

## Materials and Methods

### Histone and DNA preparation

Histone proteins were expressed and purified according to previously established protocols. Histone H3 bearing K36 substituted with cysteine (K36C) was generated by site-directed mutagenesis using an Agilent Quikchange XL Site-Directed Mutagenesis kit (Agilent #200517) with the canonical H3 gene in pET3 vector. Primers: H3K36C F: 5’-ctgctaccggcggagtctgcaaacctcaccgttaccg-3’ R: 5’-cggtaacggtgaggtttgcagactccgccggtagcag-3’. Generation of the mutant was verified by sequencing.

### LCMS-ESI+

Chemical protein modification reactions were monitored using LCMS-ESI+. Waters LCT Premier XE coupled to a Waters 1525 Micro HPLC using a Merck Chromolith C18 guard cartridge. Water (solvent A) and acetonitrile (solvent B), both containing 0.1% formic acid, were used as mobile phase at a variable flow rate. The gradient program was: 95% A (1 min isocratic, 0.4 ml/min) to 100% B after 5 min (0.4 ml/min) then isocratic for 3 min (flow rate increasing to 1.0 ml/min). The electrospray source parameters were: capillary voltage: 3000 V, cone voltage: 160 V. Nitrogen was used as the desolvation gas and nebuliser at a total flow of 600 L/hr. Data were processed using MassLynx.

### H3K36Dha synthesis

H3K36C (4.7 mg, 0.347 μMol, 1 eq) was weighed into a 1.5 mL Eppendorf tube and dissolved in 500 μL reaction buffer (5M GdnHCl, 50 mM Tris-HCl pH 8.0). Approx. 6 mg DTT was then added and the reaction was shaken for 45 minutes at 37°C. During this time a GE PD G-25 minitrap column was equilibrated with reaction buffer (3 x 2 mL). The H3K36C solution was then passed through the minitrap and eluted in 1 mL reaction buffer. Immediately afterwards, MDBP (5.13 μL in 50 μL DMSO, 100 eq.) was added, upon which the reaction became a cloudy suspension. The reaction was incubated at 37°C and aliquots taken for LCMS-ESI+ analysis at half-hourly or hourly timepoints. After 5 hours when the reaction had gone to completion as suggested by LCMS-ESI+ data, it was quenched by passing through a GE PD G-25 miditrap pre-equilibrated with Luche buffer (3 x 5 mL; 5M GdnHCl, 500 mM NH_4_OAc pH 6.0) and eluted in 1.5 mL Luche buffer into a 2 mL Eppendorf tube. Protein concentration was measured by nanodrop (ε_0_= 4470 M^-1^cm^-1^, MW= 15.18 kDa). Solutions were diluted to approx. 1 mg/ mL in Luche buffer, re-measured by nanodrop, divided into 0.75 mL aliquots and placed in a nitrogen glovebox to degas overnight at room temperature.

### H3K36me2 and -me3 synthesis

Following equilibration with N_2_ (g) overnight in anaerobic glovebox, 100 μL H3K36Dha in Luche buffer (1 mg/ mL in 5M GdnHCl, 500 mM NH_4_OAc pH 6.0) were mixed first into (3-iodopropyl)-trimethylamine (20 mg) for H3K36me3 or (3-iodopropyl)dimethylamine hydroiodide (20 mg) for H3-K36me2 then into NaBH4 (2 mg) pre-portioned into a 2 mL eppendorf tube. This was shaken at 4°C for 30 mins before being passed through GE PD G-25 spintraps pre-equilibrated with unfolding buffer (7M GdnHCl, 10 mM Tris-HCl pH 7.5, 1 mM DTT). The reactions were then analysed by LCMS-ESI+ to determine conversion.

### 186 bp biotinylated DNA preparation

Plasmid bearing 601 DNA was expressed and purified according to previously reported protocols^16^. The 186 bp fragment was excised from the plasmid by restriction digest with EcoRV (NEB: R0195) and EcoRI (NEB: R0101) according to manufacturer protocols. The reaction was incubated at 37°C for 16 h and analysed by agarose gel electrophoresis. Biotinylation was then carried out according to previously described protocols^48^. Briefly, 186 bp DNA (0.4 mg/ mL) in 1x NEBuffer 2 (NEB: B7002) was incubated with dATP (100 μM), Klenow 3’ to 5’ exo (NEB: M0212) and Biotin-11-dUTP (40 μM, Jena Bioscience, NU-803-BIOX) at 25°C for 2 h, before quenching with addition of EDTA to 20 mM and phenol/ chloroform extraction. Final DNA concentration was 2345.4 ng/ μL (260.6 μg/ L culture).

### Complex reconstitution

Histone octamer reconstitution was performed according to previously established protocols. Briefly, approximately 2 mg of each lyophilised histone were weighed into a 2 mL Eppendorf tube and dissolved in 1 mL freshly prepared unfolding buffer (6 M GdnHCl, 20 mM Tris-HCl pH 7.5, 5 mM DTT). The solutions were then incubated at room temperature for 1 hour before measuring concentrations via nanodrop. H2A, H2B, H4 and H3 wild-type (WT), K36me2 or K36me3 were then mixed in an equimolar ratio and adjusted with unfolding buffer to a final concentration of approx. 1 mg/ mL. The combined histones were then transferred to a 6 kDa Pur-a-Lyzer Mega dialysis cup, and dialysed against 3 changes of 600 mL of pre-chilled refolding buffer (2M NaCl, 10 mM Tris-HCl pH 7.5, 1 mM EDTA and 5 mM β-mercaptoethanol) at 4°C for 6, 14 and 6 h.

The octamer was concentrated to ≤1 mL using a 5 kDa MWCO Vivaspin concentrator at 4°C, and centrifuged (7000 rpm, 10 min, 4°C) to pellet any precipitate. The supernatant was then loaded onto the SEC 16/60 and eluted in refolding buffer over 1.2 CV at 0.2 mL/ min. Octamer-containing fractions were identified by retention time and/or denaturing SDS-PAGE (4-12% Bis-Tris); fractions containing all 4 histones gave 4 corresponding bands on the gel. These fractions were combined and concentrated to a volume of 1 mL using a 6 mL 6 kDa MWCO Vivaspin concentrator at 4°C, avoiding excess mechanical stress to prevent dissociation. The octamer was diluted to a final concentration of 50% glycerol for storage at −20°C.

### Nucleosome reconstitution by salt gradient dialysis

Each of the three DNA/ octamer mixtures were then pipetted into a 6 kDa Pur-a-Lyzer mini dialysis membrane and dialysed sequentially into RRTCS (10 mM Tris-HCl pH 7.5, 1 mM EDTA, 1 mM DTT with decreasing KCl concentrations of 2M, 0.85 M and 0.65 M) for 2 h each, then RRTCS 0.25 M (0.25 M KCl, 10 mM Tris-HCl pH 7.5, 1 mM EDTA, 1 mM DTT) overnight. Heat shifting was then performed as follows. Each DNA/ octamer mixture was divided into 3 aliquots and incubated for 2 h at 4°C, 37°C or 55°C. Samples were run on 6% TBE gel in 1 x TBE (150 V, 90 min, 4°C) stained with SYBR Gold (Thermo Fisher Scientific. S11494) and InstantBlue Protein Stain (Expedeon).

### PreScission protease preparation

pGEX vector bearing the gene for PreScission protease was transformed into BL21 cells and grown in LB supplemented with ampicillin (100 μg/ mL) to OD_600_ of 0.5 at 37°C, at which point the temperature was lowered to 18°C before induction with 0.2 mM IPTG. Protein expression was continued for 18 h before harvesting cells by centrifugation (4°C, 8000 rpm, 10 min in Beckman Coulter Avanti JXN-26 centrifuge, JLA 8.1 rotor).

Pelleted cells were resuspended in 100 mL PreScission Lysis buffer (PBS pH 7.4 supplemented with 150 mM NaCl, 0.1% Triton X100, 1 mM DTT) supplemented with Protease Inhibitor Cocktail (Calbiochem). Cells were lysed on ice with sonication after which DNAse I (1 μL, Roche) and MgCl_2_ (to 1 mM) were added. The lysate was incubated at 4°C with gentle rotation before ultracentrifugation (1 h, 42,000 rpm, 4°C) to pellet insoluble debris. The supernatant was decanted, and added to GST-Sepharose beads (4 mL of slurry, pre-washed three times with PreScission Lysis buffer). The GST beads were incubated (1 h, 4°C, gentle rotation) to allow the GST tag to bind, before washing with PreScission wash buffer (PBS pH 7.4 supplemented with 300 mM NaCl, 0.1% Triton X100, 1 mM DTT). The beads were then resuspended in PreScission was buffer (30 mL) and supplemented with 5 mM ATP and 15 mM MgCl_2_ and incubated (1 h, 4°C, gentle rotation) before washing twice with PreScission Lysis buffer (30 mL). Beads were washed with PreScission elution buffer (PBS pH 7.4 supplemented with 150 mM NaCl, 0.1% Triton X100, 1 mM DTT, 25 mM glutathione), and eluted PreScission dialysed in Prescission dialysis buffer (50 mM Tris-HCl pH 8.0, 150 mM KCl, 10% glycerol, 1 mM ETDA, 2 mM DTT), flash frozen in aliquots, and stored at −80°C.

### MRG15 preparation

MRG15 bearing an N-terminal His6 tag was cloned into the pGEX-6P-3 vector at BamHI/ NotI sites using standard techniques. The pGEX-6P-3 vector was a kind gift from the lab of Ivan Ahel. The plasmid was transformed into Artic Express cells and grown in LB supplemented with ampicillin (100 μg/ mL) and gentamycin (20 μg/ mL) to OD_600_ of 0.5 at 37°C, at which point the temperature was lowered to 13°C before induction with 0.2 mM IPTG. Protein expression was continued for 24 h before harvesting cells by centrifugation (4°C, 8000 rpm, 10 min in Beckman Coulter Avanti JXN-26 centrifuge, JLA 8.1 rotor).

Pelleted cells were resuspended in 30 mL Trit buffer (PBS pH 7.4 supplemented with 150 mM NaCl, 0.05% Triton X100) and Protease Inhibitor Cocktail (Calbiochem). Cells were lysed on ice with sonication after which DNAse I (1 μL, Roche) and MgCl_2_ (to 1 mM) were added. The lysate was incubated at 4°C with gentle rotation before centrifugation (45 min, 23,000 rpm, 4°C) to pellet insoluble debris. The supernatant was decanted, filtered (45 μm, Sartorius), and added to GST-Sepharose beads (2 mL of slurry, pre-washed three times with Trit buffer). The GST beads were incubated (1 h 30 min, 4°C, gentle rotation) to allow the GST tag to bind, before and washing once with Trit buffer (50 mL) and twice with wash buffer (50 mL of 50 mM Na2HPO4, 50 mM NaH2PO4, pH 7.4, 0.05% Triton X100, 200 mM NaCl, 5 mM Imidazole). The beads were then resuspended in Trit buffer (30 mL) and supplemented with 5 mM ATP and 15 mM MgCl_2_ and incubated (1 h, 4°C, gentle rotation) before washing twice with Trit buffer (30 mL). An aliquot of GST-MRG15 sepharose beads was flash frozen and stored at −80°C for GST pulldowns. The remainder was washed twice with p5 buffer (30 mL of 50 mM Na2HPO4, 50 mM NaH2PO4, pH 7.4, 0.05% Triton X100, 200 mM NaCl, 5 mM Imidazole). PreScission protease was then added to cleave hMRG15 from the immobilised GST tag, and incubated for 16 h (4°C, gentle rotation).

The GST bead slurry was then loaded onto a gravity column and the supernatant was eluted directly onto Talon beads (600 μL of slurry, pre-washed three times with 50 mL p5). The GST bead slurry was washed twice with Trit buffer, flash frozen and stored at −80°C for GST pulldown. The talon beads were incubated with the supernatant (1 h 30 min, 4°C, gentle rotation) to allow the His6 tag to bind. The beads were then loaded onto a gravity column and washed with p5 (50 mL) and p30 (50 mL of 50 mM Na_2_HPO_4_, 50 mM NaH_2_PO_4_, pH 7.4, 0.05% Triton X100, 200 mM NaCl, 30 mM Imidazole) before elution in p500 (1 mL aliquots of 50 mM Na2HPO4, 50 mM NaH2PO4, pH 7.4, 0.05% Triton X100, 200 mM NaCl, 500 mM Imidazole). Eluted fractions were then dialysed in dialysis buffer (50mM Tris-HCl pH 7.5, 150 mM NaCl, 2 mM DTT, 1 mM EDTA, 10% glycerol), flash frozen and stored at −80°C.

### MRG15 W53A preparation

MRG15 W53A was generated by site-directed mutagenesis using an Agilent Quikchange XL Site-Directed Mutagenesis kit (Agilent #200517) with the MRG15 WT in pGEX-6P-3 vector. Primers were taken from Bleuyard et. al^10^:

F: 5‘-CTTCATACATTACAGTGGTTGGAATAAAAATGCGGATGAATGGGTTCCG-3‘

R: 5‘-CGGAACCCATTCATCCGCATTTTTATTCCAACCACTGTAATGTATGAAG-3‘

MRG15 W53A was then expressed and purified following the protocol for MRG15 WT described above.

### Western blot

Proteins were transferred from the SDS-PAGE gel to a 0.2 μm Amersham Protran Nitrocellulose blotting membrane (GE life sciences) using a BioRad Trans-blot cell gel tank filled with transfer buffer (192 mM glycine, 25 mM Tris base pH 8.3, 20% methanol). Transfer was performed for 1 h at 90 V, 4°C. Bands on the membrane were then visualised using Ponceau stain (0.1% Ponceau S, 1% acetic acid) and membrane cut into sections as appropriate for primary antibody application. Secondary antibody was polyclonal goat α rabbit IgG HRP conjugate (Dako, lot 20066477, 1:1000 dilution). ECL (Amersham Biosciences) was used to visualise the bound secondary antibody. The membrane was covered in ECL and incubated in the dark (2 min, room temp) before exposing in a cassette (Hypercasette, Amersham Biosciences) and developing onto chemiluminescence film (Amersham Hyperfilm ECL) using a Xenograph SRX-101A developer.

### GST-MRG15 pulldown with histones

GST or GST-MRG15 on GSH Sepharose beads were blocked in NET250 BSA buffer (50 mM Tris-HCl pH 7.5, 250 mM NaCl, 0.5% NP40, 2 mM DTT, 5 mg/ mL BSA) for 2 h at 4°C. 1 μg of histone (H3 WT, H3-K36me2, H3-K36me3 or H4) was then added, and the samples incubated for a further 1 h at 4°C. The beads were then washed three times with NET250 BSA to remove unbound histones, made up in 2 x SDS (NuPAGE LDS sample buffer + 80 mM DTT) and run on SDS-PAGE in MES-SDS buffer (200 V, 40 min, 4°C) and imaged via Western blot. PBS with 0.05% Tween20 was used for washing steps, and PBS-tween with 5% milk powder was used for blocking steps. Primary antibodies used were: rabbit α-GST (Santa Cruz sc138 mouse monoclonal IgG_1_ α-GST HRP conjugate, lot D2208) for GST and GST-MRG15 portions of membrane, and rabbit α-histone H3 for histone portion of membrane (Bethyl, A300-B22 A-1, 1:1000 dilution).

### Streptavidin pulldown with biotinylated nucleosomes and MRG15

Biotinylated 186 bp nucleosomes were immobilised onto Dynabeads Streptavidin MyOne T1 beads according to the manufacturer’s protocol with modifications. Briefly, eight 5 μL aliquots of beads were washed in three times 1 mL HBK (10 mM HEPES pH 7.5, 150 mM KCl, 1 mM EDTA, 1 mM DTT, 0.1% NP40) at 4°C. Each aliquot of beads was then resuspended in 1 mL of the appropriate buffer and 0.2 μg of each 186 bp nucleosome variant or free 186 bp DNA was added. The beads were then incubated with gentle rotation (15 rpm, 30 min room temp) to allow nucleosome or DNA to bind. The beads were then washed carefully three times with 1 mL HBK with 0.1% BSA, and resuspended in 1 mL HBK with 5% BSA. The beads were incubated for a further 1h with gentle rotation (15 rpm, 4°C) to ensure complete blocking; salmon sperm DNA (20 μg) and MRG15 (0.1 μg) were added as appropriate before incubating for a further 4 h (15 rpm, 4°C) to allow binding interactions between protein and/or DNA to equilibrate.

The beads were then washed four times with binding/ washing buffer to remove any unbound hMRG15. They were then made up in 2 x SDS (NuPAGE LDS sample buffer + 80 mM DTT) and run on SDS-PAGE in MES-SDS buffer (200 V, 40 min, 4°C) and imaged via Western blot. TBS with 0.05% Tween20 was used for washing steps, and TBS-tween with 5% BSA was used for blocking steps. Primary antibodies used were: rabbit α-MRG15 (Cell Signalling Technologies, lot 1, 1:1000 dilution) for MRG15 portion of membrane, and rabbit α-H3 for histone portion of membrane (Bethyl, A300-B22 A-1, 1:1000 dilution).

### Overlay and docking

UCSF Chimera was used to overlay MRG15 chromodomain (PDB ID 2F5K) with structures of chromodomains of HP1, MRG2, CHP1 and MMP8 0bound to trimethyllysine; 1KNE, 4PL6, 2RSN/3G7L and 3QO2, respectively. Initial docking was carried out using GOLD. The M3L ligand was built in Avogadro and saved as an .sdf file. 4PL6 was used as the template for docking with residues 4 Å away from the M3L ligand used to define the binding site and the detect cavity option was not selected. Hydrogens were added to all structures. 2F5K was used as the target for the rigid docking. No protonation states were changed and the following residues were given constrained rotamer values from the GOLD library. Residues constrained were 21, 26, 46, 49, 53, 50, 55. GOLD was allowed to generate 100 possible conformers and the highest scoring docking results were calculated using the PLP score.

## Acknowledgements

FE is supported by Wellcome Trust Senior Research Fellowships in Basic Biomedical Science (101009/Z/13/Z) and is thankful for support from the Edward P. Abraham Research Fund. The Rosalind Franklin Institute Funded by UK Research and Innovation through the Engineering and Physical Sciences Research Council. SF is a recipient of the Cancer Research UK Non-clinical Training Award (C5255/A20936). BJ is a recipient of the Clarendon Scholarship. TW is grateful to the EPSRC Centre for Doctoral Training in Synthesis for Biology and Medicine (EP/L015838/1) for a studentship, generously supported by AstraZeneca, Diamond Light Source, Defence Science and Technology Laboratory, Evotec, GlaxoSmithKline, Janssen, Novartis, Pfizer, Syngenta, Takeda, UCB and Vertex. We thank Profs V. Gouverneur and F. Duarte Gonzalez for their advice and guidance, and Dr MC. Keogh for sharing unpublished results and discussion.

## Author contributions

FE, BGD and SF conceived and planned the project. SF generated and assembled synthetic nucleosomes, purified MRG15 and conducted binding experiments, assisted by AC and BJ. TW conducted structure alignments and associated analyses. FE, BGD and SF wrote the manuscript with input from all contributing authors.

## Conflict of interest

We declare no financial or non-financial conflict of interests on this study.

## Supporting Information

- Appendix (Source Data)

## Supplemental Figure Legends

**Figure S1:**
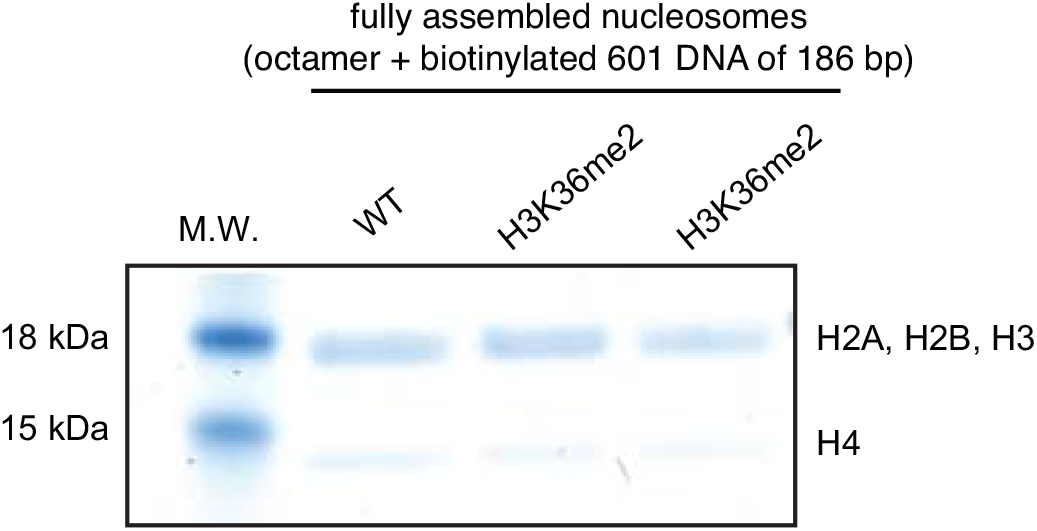
Denaturing SDS-PAGE Analysis of Nucleosomes. Staining of SDS-PAGE of assembled nucleosomes with InstantBlue Protein Stain (Expedeon) shows expected histone composition for all species. M.W: GeneFlow BluEye pre-stained protein molecular weight marker.

**Figure S2:**
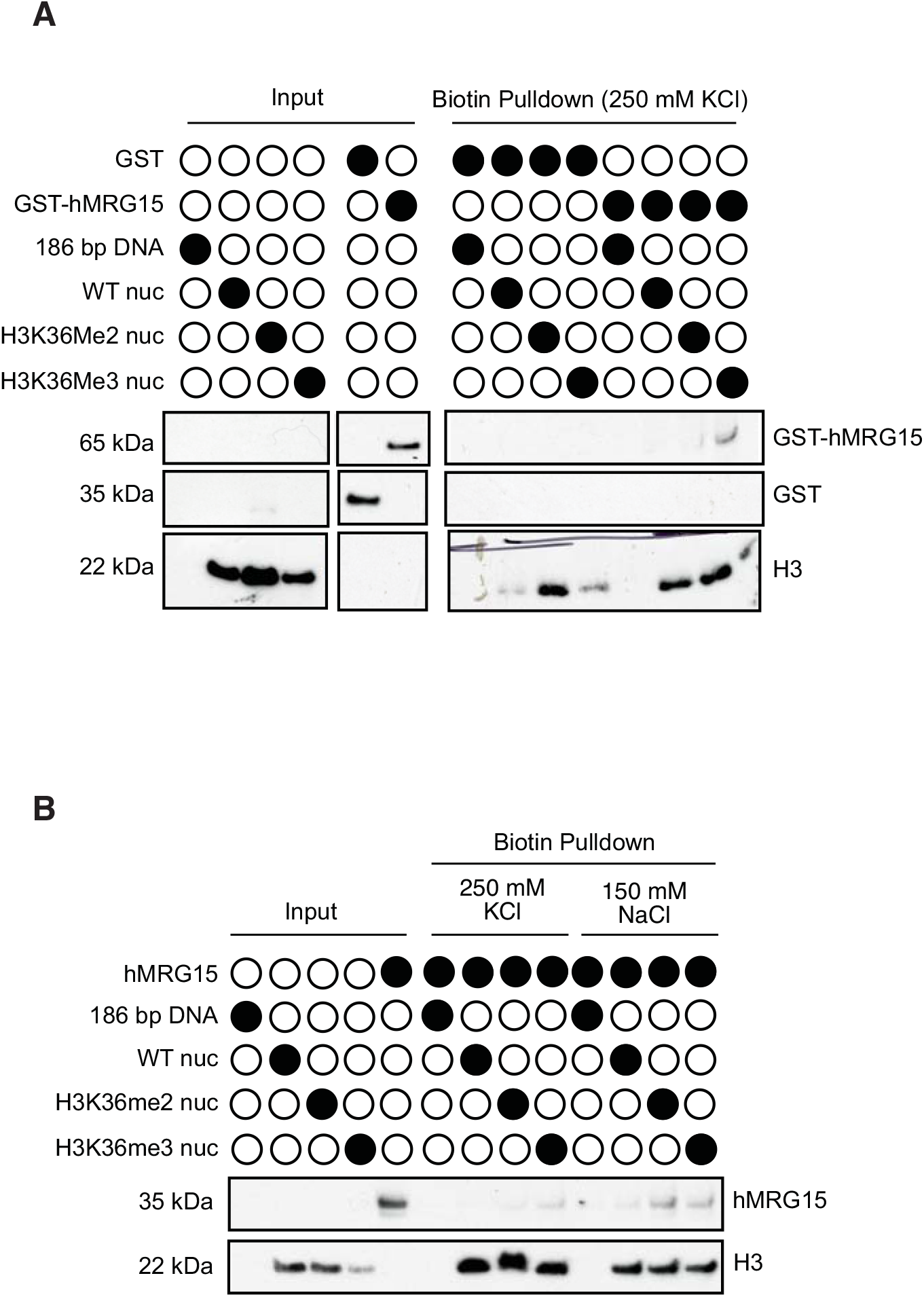
Pulldown Experiment with GST- or his6-tagged MRG15. **A,** Nucleosomes, comprising 186 bp biotinylated DNA, were immobilised on streptavidin dynabeads and beads were preincubated with BSA at a blocking step. Unbound nucleosomes and Bovine Serum Albumin (BSA) were then washed from the beads with 250 mM KCl buffer. Subsequently, the beads were incubated with GST-MRG15 or GST alone (negative control) with BSA and salmon sperm DNA to out-compete non-specific interactions. Unbound species were then removed in wash steps, and the beads analysed by SDS-PAGE and Western blot. **B,** As in A, except using his6-MRG15 in binding buffer containing 250 mM KCl or 150 mM NaCl. Unbound species were removed in wash steps, and the beads analysed by SDS-PAGE and Western blot.

**Figure S3:**
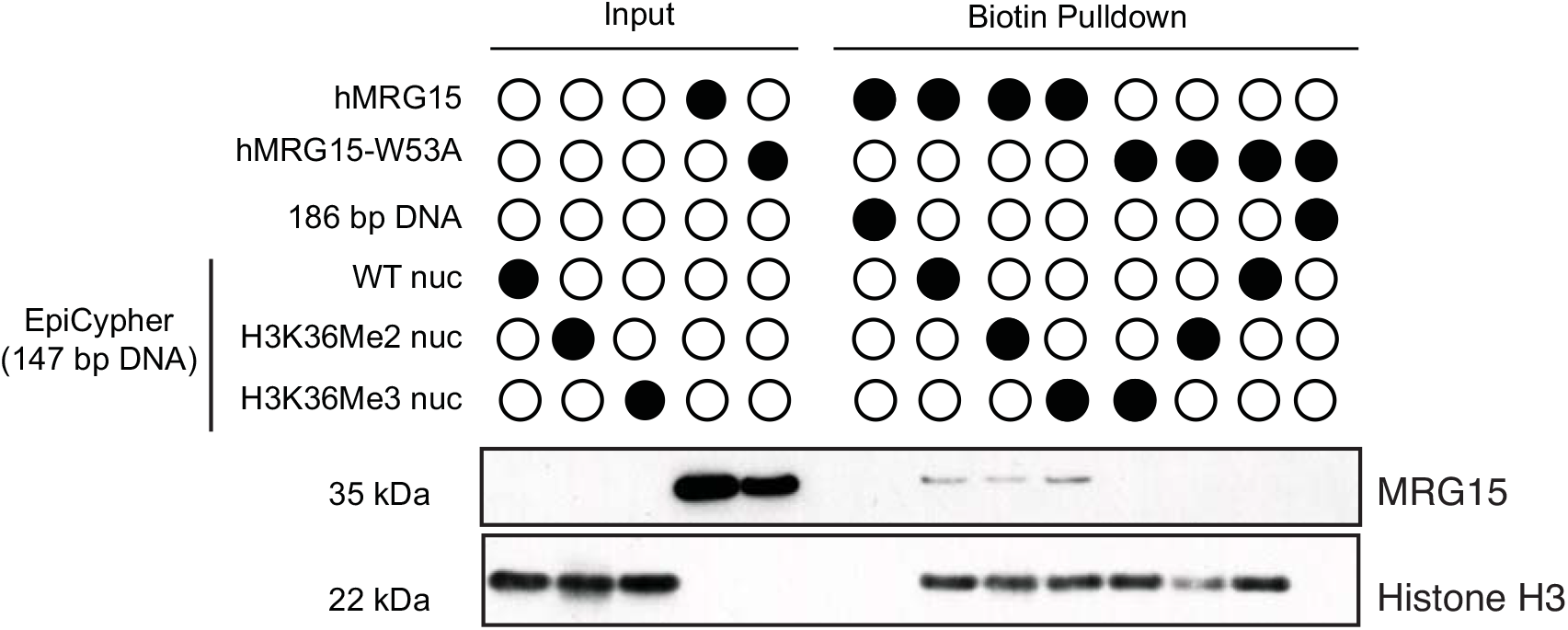
Pulldown Experiment using MRG15 and EpiCypher nucleosomes. EpiCypher nucleosomes, comprising 147 bp biotinylated DNA, were immobilised on streptavidin dynabeads and beads were preincubated with BSA at a blocking step. Unbound nucleosomes and Bovine Serum Albumin (BSA) were then washed from the beads with 150 mM KCl buffer. Subsequently, the beads were incubated with his-tagged wild type (WT) MRG15 or the W53A mutant in 150 mM KCl buffer in the presence of BSA and salmon sperm DNA to out-compete non-specific interactions. Unbound species were then removed in wash steps, and the beads analysed by SDS-PAGE and Western blot.

**Figure S4:**
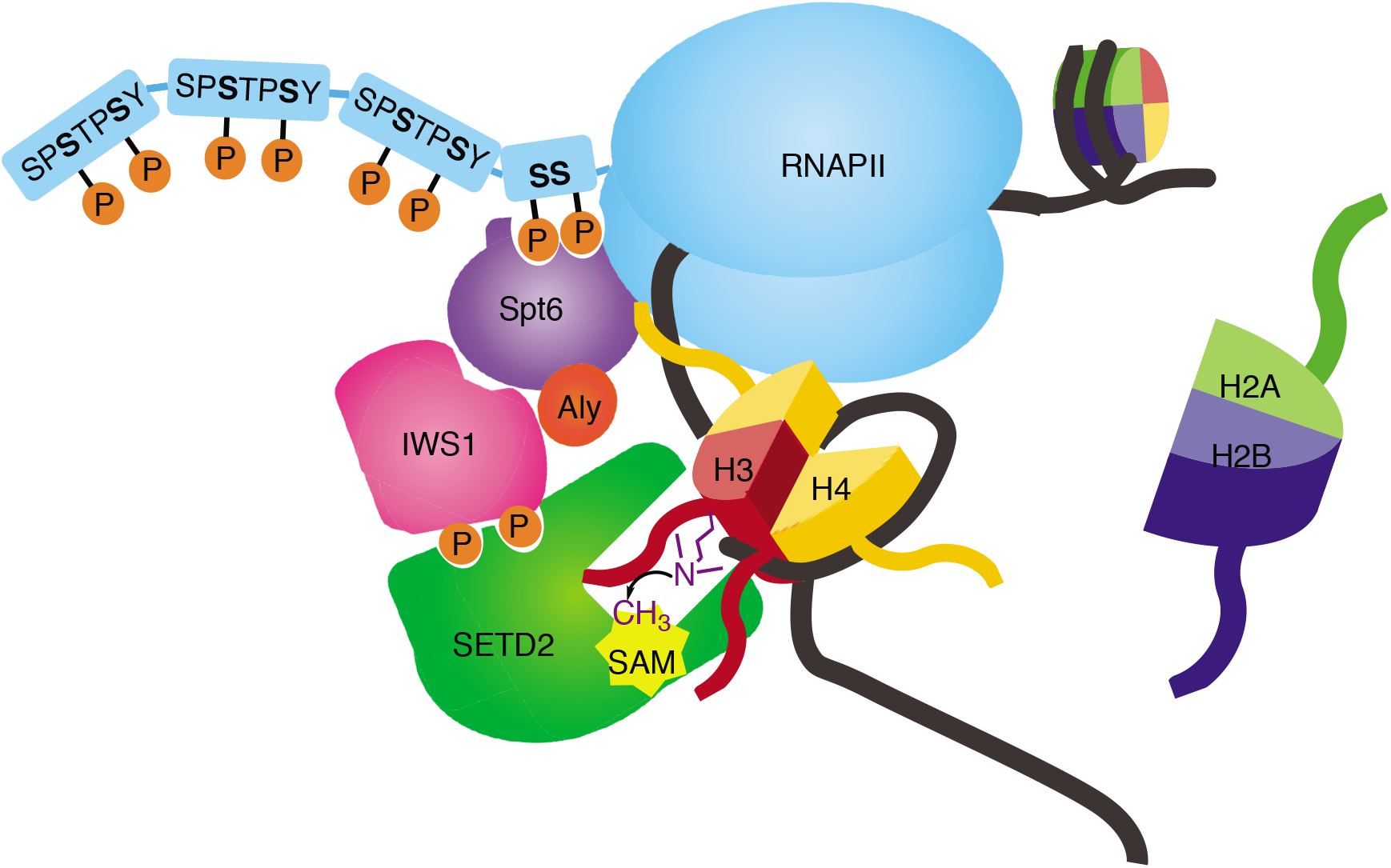
A model for SETD2-catalysed methylation of H3K36me2. A depiction of a model where RNA polymerase II associated SETD2 may catalyse trimethylation of H3K36 during transcription where DNA-H3K36 contacts are loosened and/or nucleosomes are partially disassembled.

## Notes

### Competing Interest Statement

The authors have declared no competing interest.

## References

1. Strahl, B. D. & Allis, C. D. The language of covalent histone modifications. Nature 403, 41–45 (2000).

2. Bannister, A. J. & Kouzarides, T. Regulation of chromatin by histone modifications. Cell Res.. 21, 381–395 (2011).

3. Rea, S. et al. Regulation of chromatin structure by site-specific histone H3 methyltransferases. Nature 406, 593–599 (2000).

4. Murray, K. The Occurrence of iε-N-Methyl Lysine in Histones. Biochemistry 3, 10–15 (1964).

5. Heintzman, N. D. et al. Distinct and predictive chromatin signatures of transcriptional promoters and enhancers in the human genome. Nat. Genet. 39, 311 (2007).

6. Edmunds, J. W., Mahadevan, L. C. & Clayton, A. L. Dynamic histone H3 methylation during gene induction: HYPB/Setd2 mediates all H3K36 trimethylation. EMBO J. 27, 406–420 (2008).

7. Zhu, X. et al. Identification of functional cooperative mutations of SETD2 in human acute leukemia. Nat. Genet. 46, 287 (2014).

8. Newbold, R. F. & Mokbel, K. Evidence for a tumour suppressor function of SETD2 in human breast cancer: a new hypothesis. Anticancer Res. 30, 3309–11 (2010).

9. Hu, M. et al. Histone H3 lysine 36 methyltransferase Hypb/Setd2 is required for embryonic vascular remodeling. Proc. Natl. Acad. Sci. 107, 2956–2961 (2010).

10. Bleuyard, J.-Y. et al. MRG15-mediated tethering of PALB2 to unperturbed chromatin protects active genes from genotoxic stress. Proc. Natl. Acad. Sci. 114, 7671–7676 (2017).

11. Hayakawa, T. et al. MRG15 binds directly to PALB2 and stimulates homology-directed repair of chromosomal breaks. J. Cell Sci. 123, 1124–1130 (2010).

12. Gautam, D., Johnson, B. A., Mac, M. & Moody, C. A. SETD2-dependent H3K36me3 plays a critical role in epigenetic regulation of the HPV31 life cycle. PLOS Pathog. 14, e1007367 (2018).

13. Hayakawa, T. et al. RBP2 is an MRG15 complex component and down-regulates intragenic histone H3 lysine 4 methylation. Genes to Cells 12, 811–26 (2007).

14. Zhang, P. et al. Structure of human MRG15 chromo domain and its binding to Lys36-methylated histone H3. Nucleic Acids Res. 34, 6621–6628 (2006).

15. Wright, T. H. et al. Posttranslational mutagenesis: A chemical strategy for exploring protein side-chain diversity. Science 354, 597–623 (2016).

16. Lercher, L. et al. Generation of a synthetic GlcNAcylated nucleosome reveals regulation of stability by H2A-Thr101 GlcNAcylation. Nat. Commun. 6, 7978 (2015).

17. Wells, D. E. Compilation analysis of histones and histone genes. Nucleic Acids Res. 14, 119–149 (1986).

18. Richmond, T. J., Finch, J. T., Rushton, B., Rhodes, D. & Klug, A. Structure of the nucleosome core particle at 7 Å resolution. Nature 311, 532–537 (1984).

19. Luger, K., Suto, R. K., Clarkson, M. J. & Tremethick, D. J. Crystal structure of a nucleosome core particle containing the variant histone H2A.Z. Nat. Struct. Biol. 7, 1121–1124 (2000).

20. Luger, K., Rechsteiner, T. J., Flaus, A. J., Waye, M. M. . & Richmond, T. J. Characterization of nucleosome core particles containing histone proteins made in bacteria. J. Mol. Biol. 272, 301–311 (1997).

21. Nielsen, P. R. et al. Structure of the HP1 chromodomain bound to histone H3 methylated at lysine 9. Nature 416, 103–107 (2002).

22. Wang, H., Farnung, L., Dienemann, C. & Cramer, P. Structure of H3K36-methylated nucleosome-PWWP complex reveals multivalent cross-gyre binding. Nat. Struct. Mol. Biol. 27, 8–13 (2019).

23. Hughes, R. M., Wiggins, K. R., Khorasanizadeh, S. & Waters, M. L. Recognition of trimethyllysine by a chromodomain is not driven by the hydrophobic effect. Proc. Natl. Acad. Sci. 104, 11184–11188 (2007).

24. Jacobs, S. A. & Khorasanizadeh, S. Structure of HP1 chromodomain bound to a lysine 9-methylated histone H3 tail. Science 295, 2080–3 (2002).

25. Luger, K., Mä Der, A. W., Richmond, R. K., Sargent, D. F. & Richmond, T. J. Crystal structure of the nucleosome core particle at 2.8 A° resolution. Nature 389, 251–260 (1997).

26. Dyer, P.N., Edayathumangalam, R.S., White, C.L., Bao, Y., Chakravarthy, S., Muthurajan, U.M., Luger, K. Reconstitution of Nucleosome Core Particles from Recombinant Histones and DNA. Methods Enzymol. 375, 23–44 (2004).

27. Dick, D. A. The distribution of sodium, potassium and chloride in the nucleus and cytoplasm of Bufo bufo oocytes measured by electron microprobe analysis. J. Physiol. 284, 37 (1978).

28. Century, T. J., Fenichel, I. R. & Horowitz, S. B. The concentrations of water, sodium and potassium in the nucleus and cytoplasm of amphibian oocyte. J. Cell Sci 7, 5–13 (1970).

29. Olsson, M. H. M., Søndergaard, C. R., Rostkowski, M. & Jensen, J. H. PROPKA3: Consistent Treatment of Internal and Surface Residues in Empirical p *K* _a_ Predictions. J. Chem. Theory Comput. 7, 525–537 (2011).

30. Allahverdi, A., Chen, Q., Korolev, N. & Nordenskiöld, L. Chromatin compaction under mixed salt conditions: Opposite effects of sodium and potassium ions on nucleosome array folding. Sci. Rep. 5, 8512 (2015).

31. Urbic, T. Ions increase strength of hydrogen bond in water. Chem. Phys. Lett. 610–611, 159–162 (2014).

32. M-C Keogh (Chief Scientific officer, EpiCypher). Personal communication. (2020).

33. H3K36Me3 Recombinant Nucleosome (dNuc) for Enzyme Screening Assays. Available at: https://www.epicypher.com/products/nucleosomes/nucleosome-recombinant-human-h3k36me3-dnuc. (Accessed: 10th April 2020)

34. H3K36Me2 Recombinant Nucleosome (dNuc) for Enzyme Screening Assays. Available at: https://www.epicypher.com/products/nucleosomes/nucleosome-recombinant-human-h3k36me2-dnuc-biotinylated. (Accessed: 10th April 2020)

35. Shah, R.N., et al. Examining the Roles of H3K4 Methylation States with Systematically Characterized Antibodies. Mol Cell. 72, 162–177 (2018)

36. Grzybowski, A.T., Chen, Z., Ruthenburg, A.J. Calibrating ChIP-Seq with Nucleosomal Internal Standards to Measure Histone Modification Density Genome Wide. Mol Cell. 58, 886–99 (2015).

37. Weinberg, D. N. et al. The histone mark H3K36me2 recruits DNMT3A and shapes the intergenic DNA methylation landscape. Nature 573, 281–286 (2019).

38. Van Nuland, R. et al. Nucleosomal DNA binding drives the recognition of H3K36-methylated nucleosomes by the PSIP1-PWWP domain. Epigenetics & Chromatin 6, 12 (2013).

39. Musselman, C. A. et al. Binding of PHF1 Tudor to H3K36me3 enhances nucleosome accessibility. Nature Comm. 4, 2969 (2013)

40. Tian, W. et al. The HRP3 PWWP domain recognizes the minor groove of doublestranded DNA and recruits HRP3 to chromatin. Nucleic Acids Res. 47, 5436–5448 (2019).

41. Jani, K. S. et al. Histone H3 tail binds a unique sensing pocket in EZH2 to activate the PRC2 methyltransferase. Proc. Natl. Acad. Sci. U. S. A. 116, 8295–8300 (2019).

42. Sankaran, S. M., Wilkinson, A. W. & Gozani, O. A PWWP domain of histone-lysine N-methyltransferase NSD2 binds to dimethylated Lys36 of histone H3 and regulates NSD2 function at chromatin. J. Biol Chem. 291, 8465–74 (2016).

43. Larschan, E. et al. MSL Complex Is Attracted to Genes Marked by H3K36 Trimethylation Using a Sequence-Independent Mechanism. Mol. Cell 28, 121–133 (2007).

44. Gibson, M. D., Gatchalian, J., Slater, A., Kutateladze, T. G. & Poirier, M. G. PHF1 Tudor and N-terminal domains synergistically target partially unwrapped nucleosomes to increase DNA accessibility. Nucleic Acids Res. 45, 3767–3776 (2017).

45. Li, J. et al. SETD2: An epigenetic modifier with tumor suppressor functionality. Oncotarget 7, 50719–50734 (2016).

46. Yoh, S. M., Lucas, J. S. & Jones, K. A. The Iws1:Spt6:CTD complex controls cotranscriptional mRNA biosynthesis and HYPB/Setd2-mediated histone H3K36 methylation. Genes Dev. 22, 3422–3434 (2008).

47. Kulaeva, O. I., Hsieh, F.-K., Chang, H.-W., Luse, D. S. & Studitsky, V. M. Mechanism of transcription through a nucleosome by RNA polymerase II. Biochim. Biophys. Acta 1829, 76–83 (2013).

48. Bartke, T. et al. Nucleosome-interacting proteins regulated by DNA and histone methylation. Cell 143, 470–484 (2010).

