## Appendix for "Intact nucleosomal context enables chromodomain reader MRG15 to distinguish H3K36me3 from -me2"

### Appendix Part I: Unprocessed and Additional Data for Figures

#### Additional data for Fig. 1B (Appendix Figs. 1-4)

Schematics of H3K36C treatment with DTT

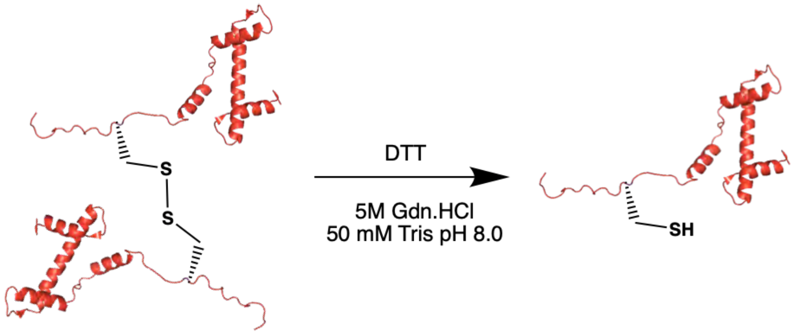


LCMS-ESI+ spectra for DTT treatment of H3K36C: Chromatogram

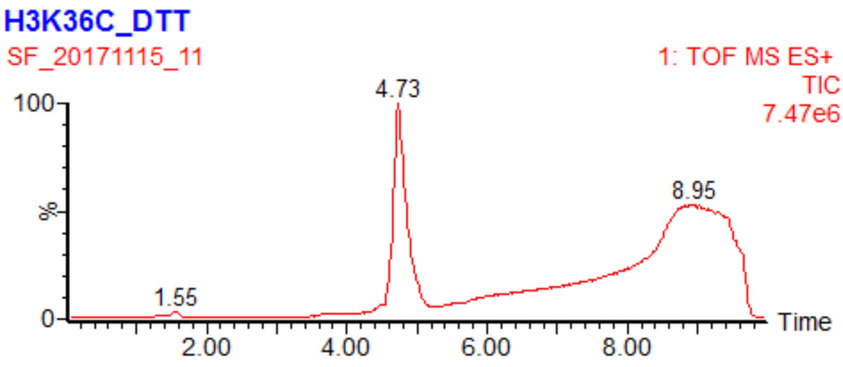


Ion series

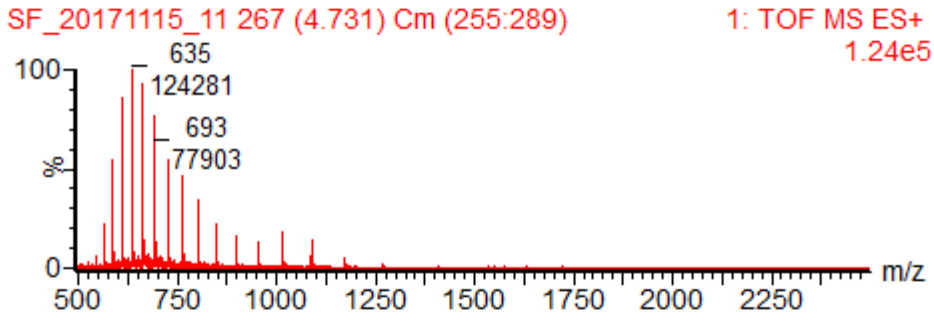


Deconvoluted mass spectrum

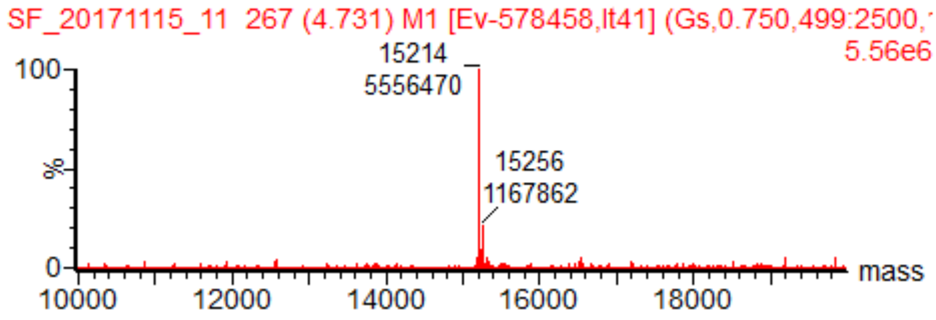


H3K36C: Expect 15214Da [M], 15256 Da [M+ MeCN]; found 15214 Da [M], 15256 Da [M+ MeCN]

**Appendix Fig. 1:** LCMS- ESI+ characterization of H3K36C treated with DTT.

Schematics of H3K36C conversion to H3K36Dha

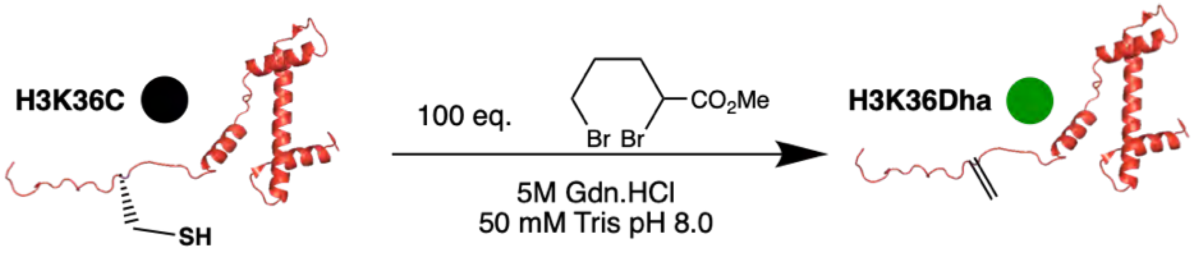


LCMS-ESI+ spectra for H3K36C conversion to H3K36Dha: Chromatogram

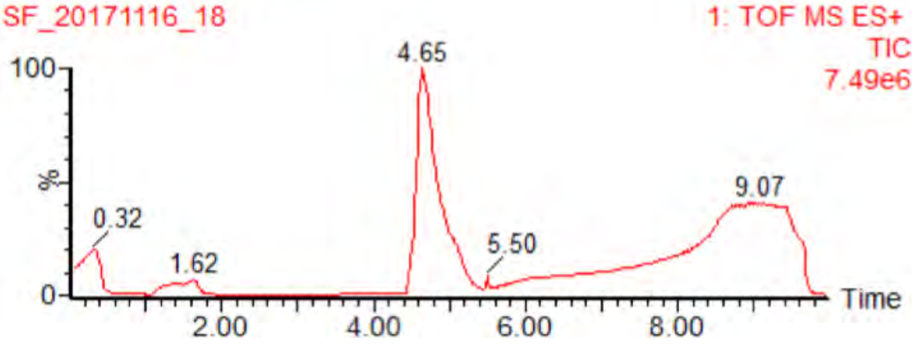


Ion series

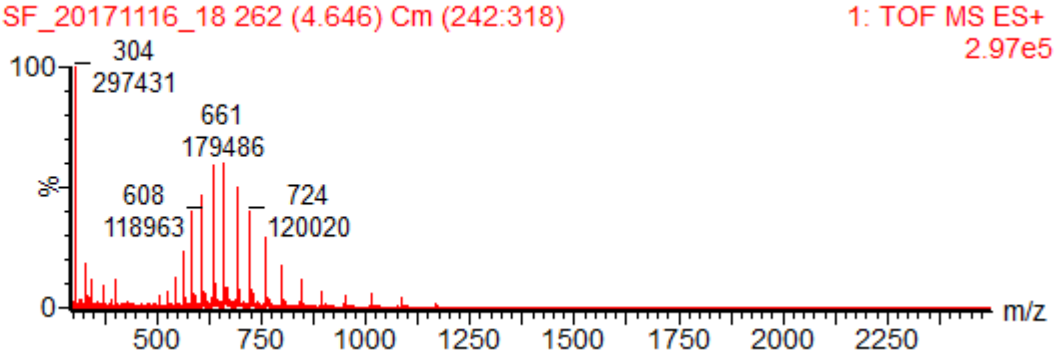


Deconvoluted mass spectrum

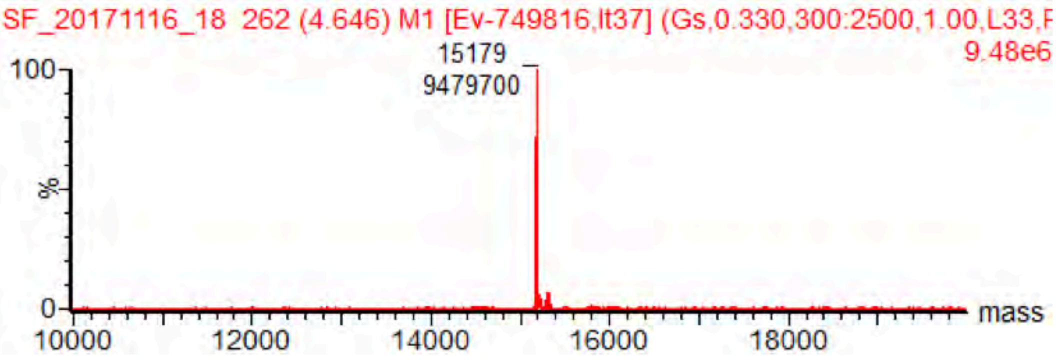


H3K36Dha: Expect 15181 Da, found 15180 Da

**Appendix Fig. 2:** LCMS-ESI+ characterization data for H3K36Dha

Schematics of H3K36Dha conversion to H3K36me2

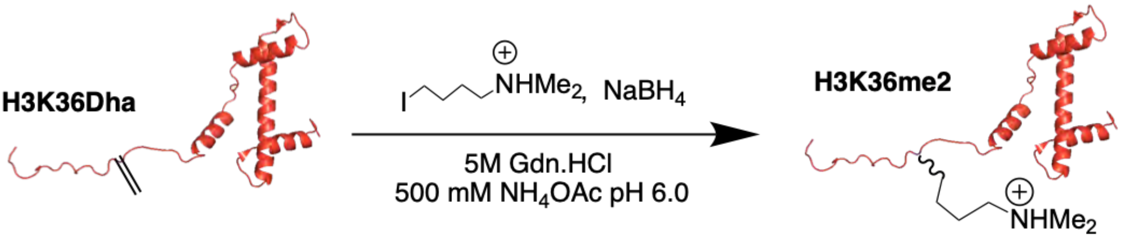


LCMS-ESI+ spectra for H3K36Dha conversion to H3K36me2: Chromatogram

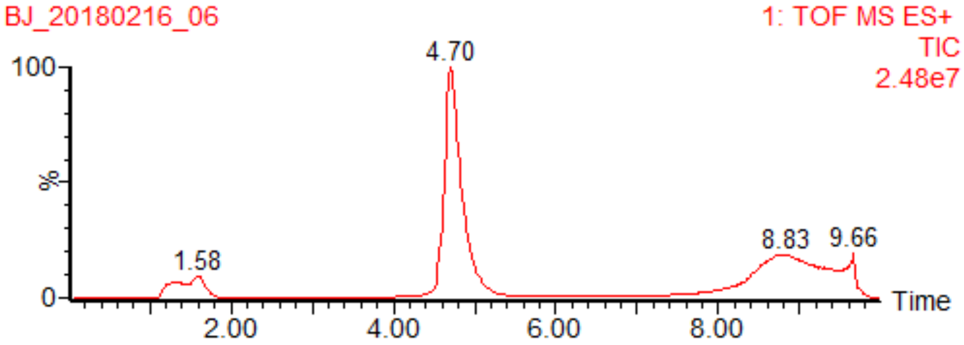


Ion series

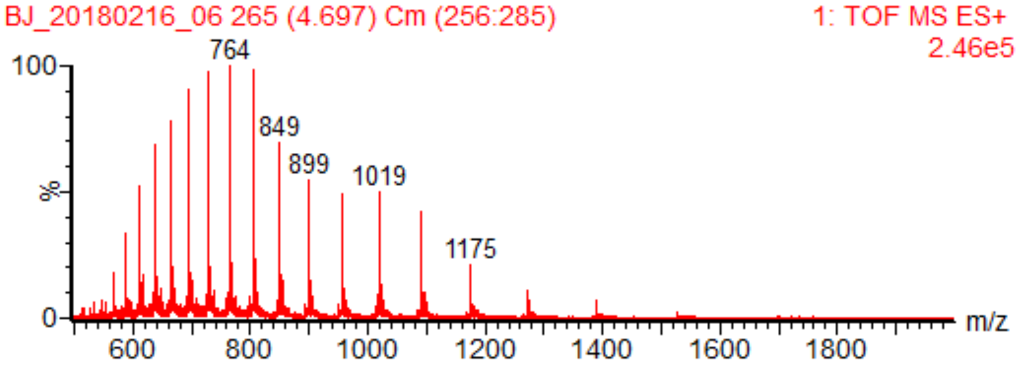


Deconvoluted mass spectrum

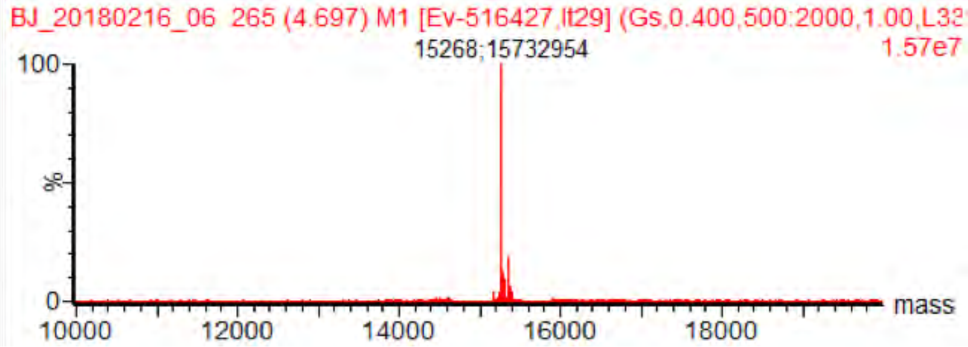


H3K36me2: Expected 15267 Da, observed 15268 Da.

**Appendix Fig. 3:** LCMS-ESI+ characterization data for H3K36me2.

Schematics of H3K36Dha conversion to H3K36me3

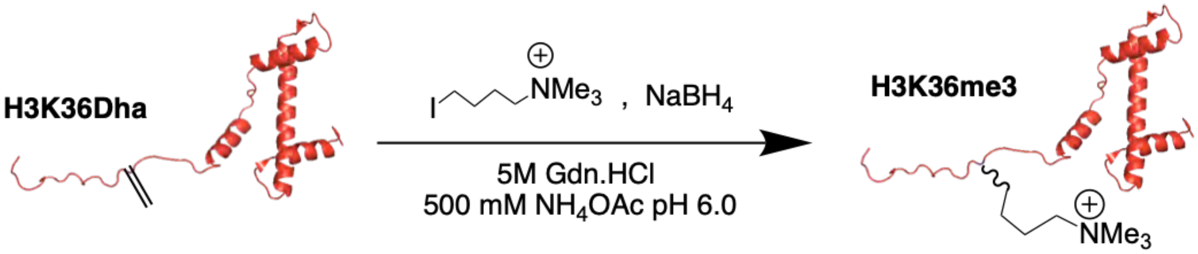


LCMS-ESI+ spectra for conversion to H3K36me3: Chromatogram

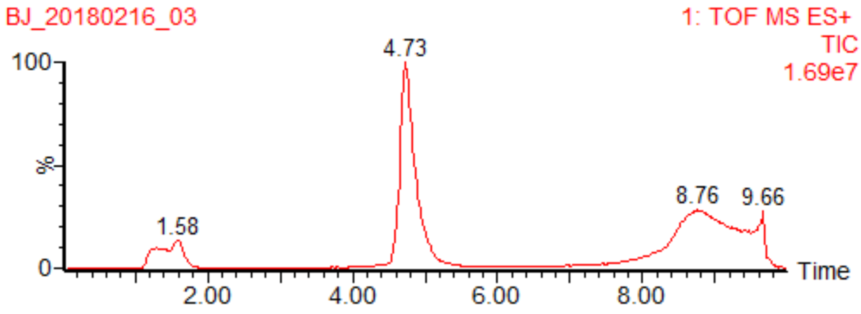


Ion series

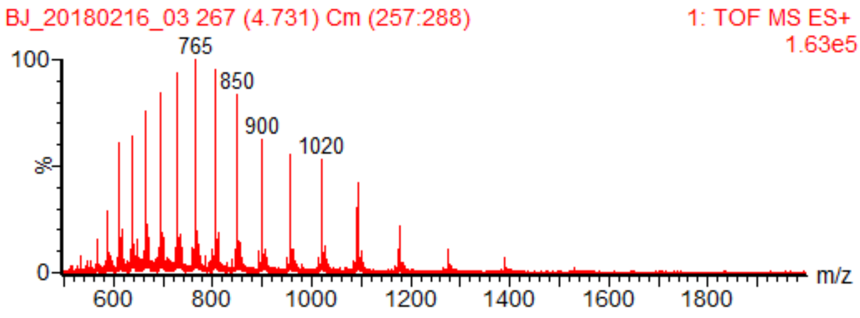


Deconvoluted mass spectrum

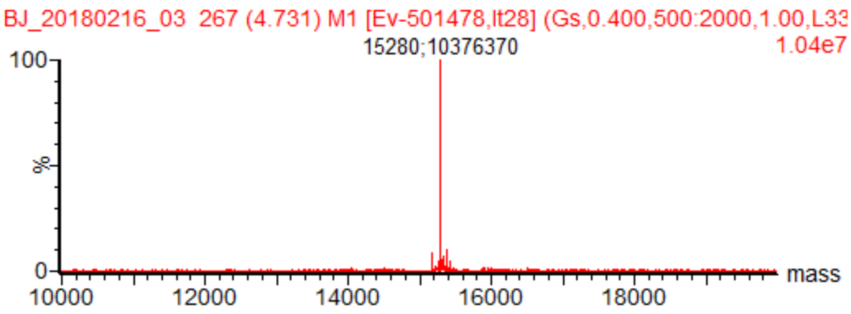


H3K36me3: Expected 15280 Da, observed 15280 Da.

**Appendex Fig. 4:** LCMS-ESI+ characterization data for H3K36me3

#### Additional data for Fig. 1C (Appendix Figs. 5-8 and Appendix Table 1)

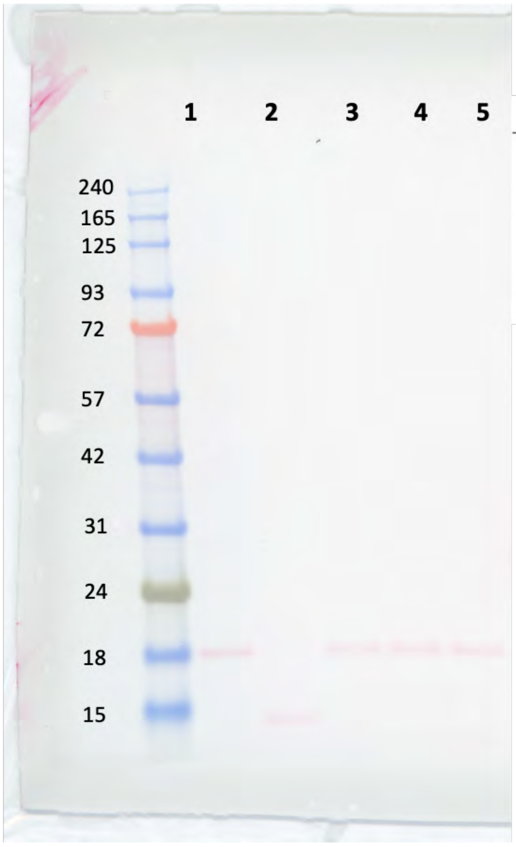


| Number | Description |
|---|---|
| 1 | H3 WT |
| 2 | H4 |
| 3 | H3-K36me3 |
| 4 | H3-K36me2 |
| 5 | H3 WT non refolded |

**Appendix Fig. 5:** Uncropped Ponceau-stained membrane for histone pulldown input (Fig. 1c) The table to the right shows sample description of lanes.

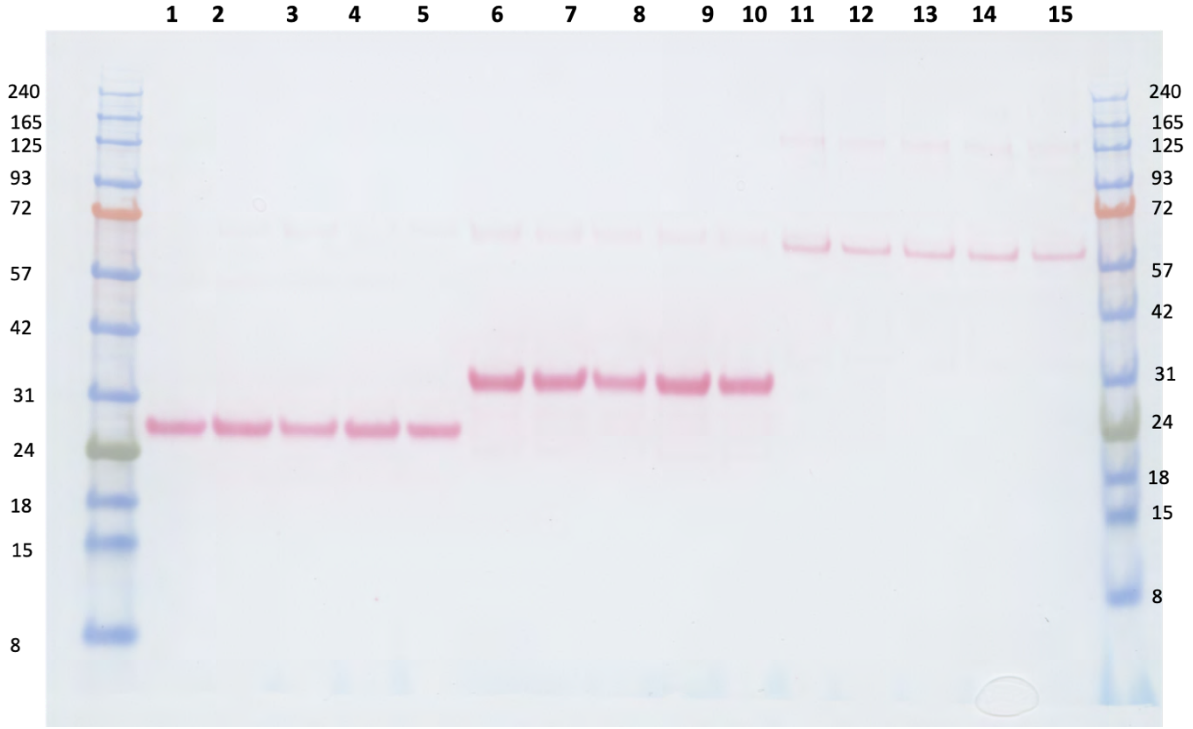


**Appendix Fig. 6:** Uncropped Ponceau-stained membrane for histone pulldown (Fig. 1c). See Appendix Table 1 for description of lanes.

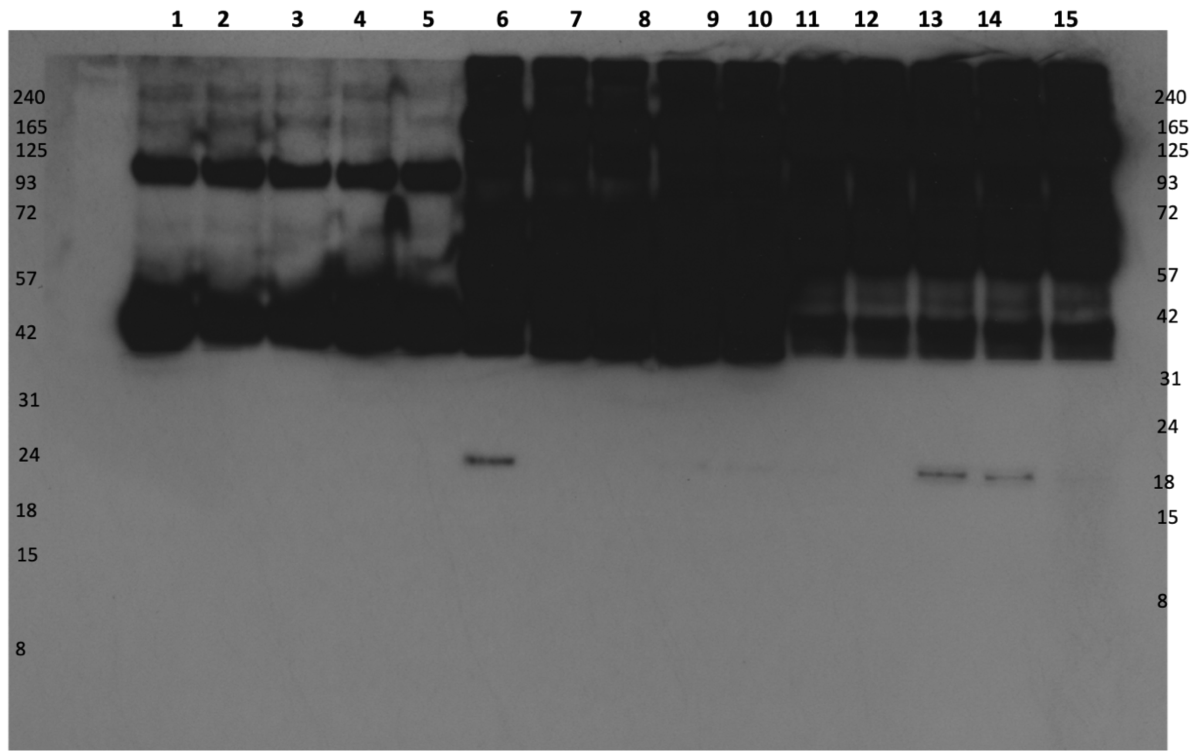


**Appendix Fig. 7:** Uncropped Western blot for histone pulldown (Fig. 1c) (short exposure). See Appendix Table 1 for description of lanes.

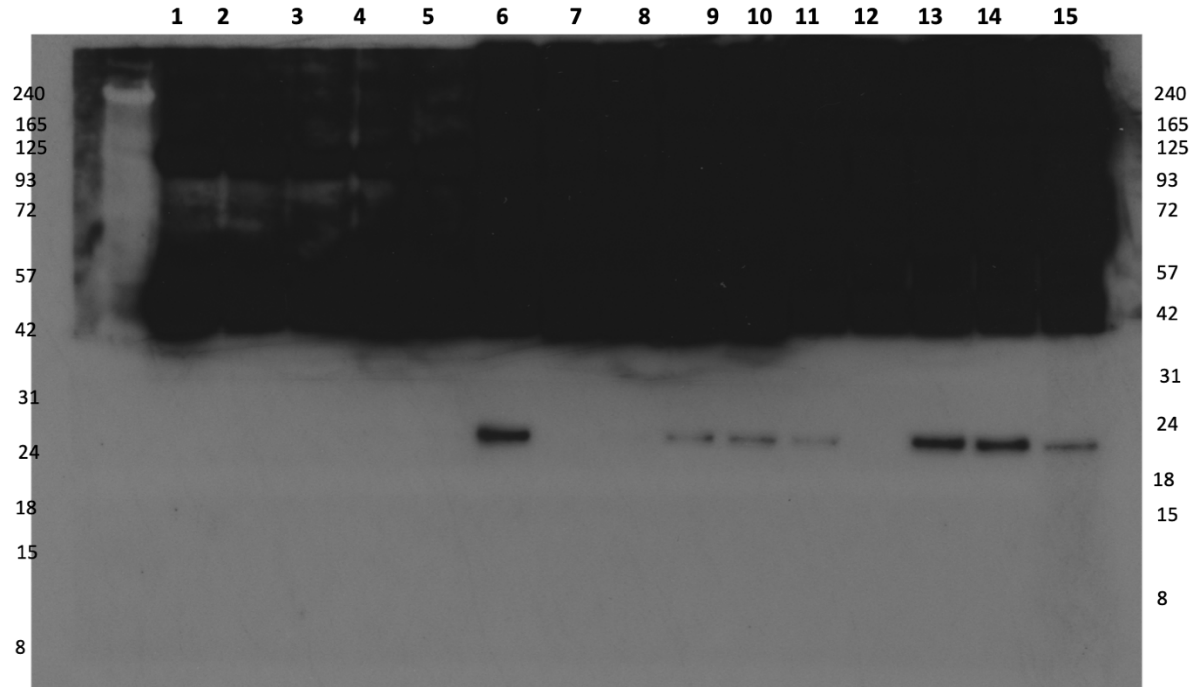


**Appendix Fig. 8:** Uncropped Western blot for histone pulldown (Fig. 1c) (long exposure). See Appendix Table 1 for description of lanes.

| *Number* | *Description* |
| --- | --- |
| *1* | H3 WT/ GST |
| *2* | H4/ GST |
| *3* | H3-K36me3/ GST |
| *4* | H3-K36me2/ GST |
| *5* | H3 WT non-refolded/ GST |
| *6* | H3 WT/ GST-PALB2 CHAM domain |
| *7* | H4/ GST-PALB2 CHAM domain |
| *8* | H3-K36me3/ GST-PALB2 CHAM domain |
| *9* | H3-K36me2/ GST-PALB2 CHAM domain |
| *10* | H3 WT non-refolded/ GST-PALB2 CHAM domain |
| *11* | H3 WT/ GST-MRG15 |
| *12* | H4/ GST-MRG15 |
| *13* | H3-K36me3/ GST-MRG15 |
| *14* | H3-K36me2/ GST-MRG15 |
| *15* | H3 WT non-refolded/ GST-MRG15 |

**Appendix Table 1:** Description of lanes in Appendix Figs 6, 7 and 8.

**Additional data for Fig. 3B (Appendix Figs. 9-11)**

<u>WT octamer purification</u>

UV trace from SEC

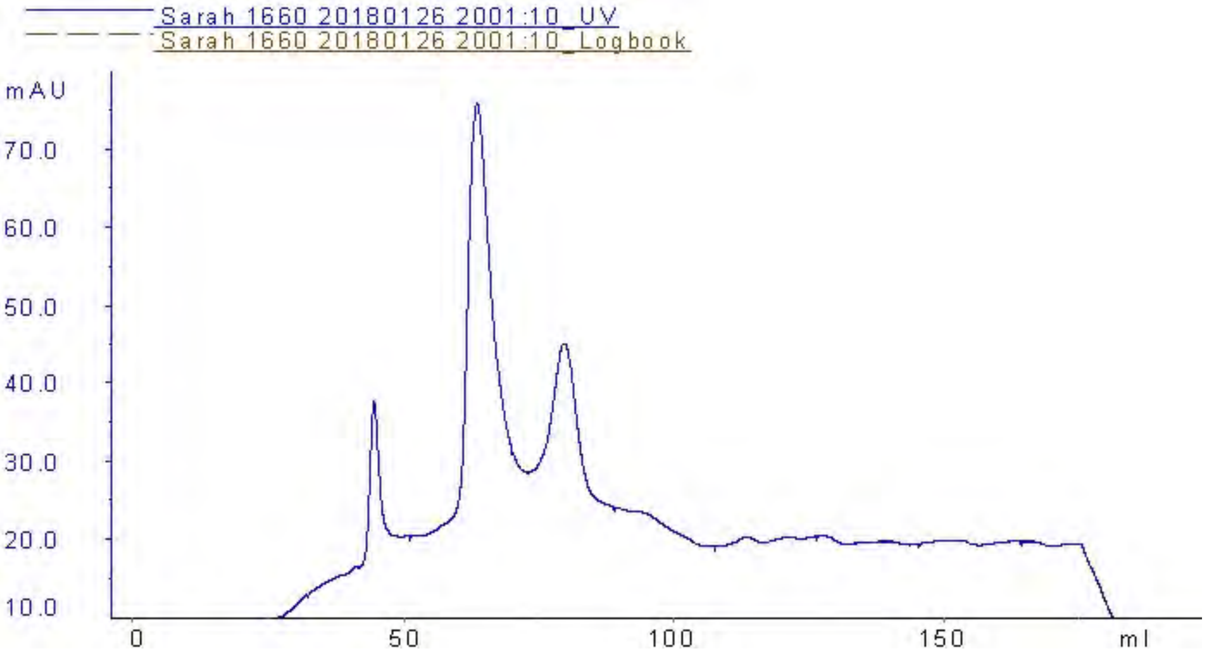


SDS-PAGE

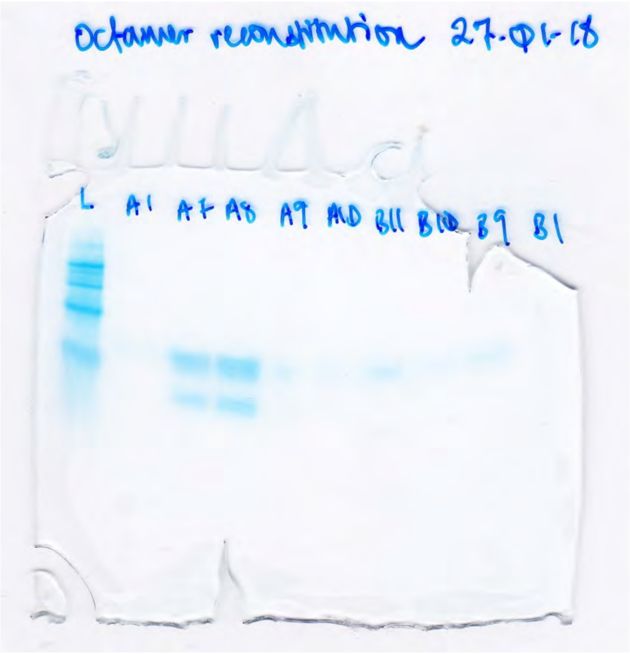


**Appendix Fig. 9:** Characterization data for H3 WT containing nucleosome

H3K36me2 octamer purification

UV trace from SEC

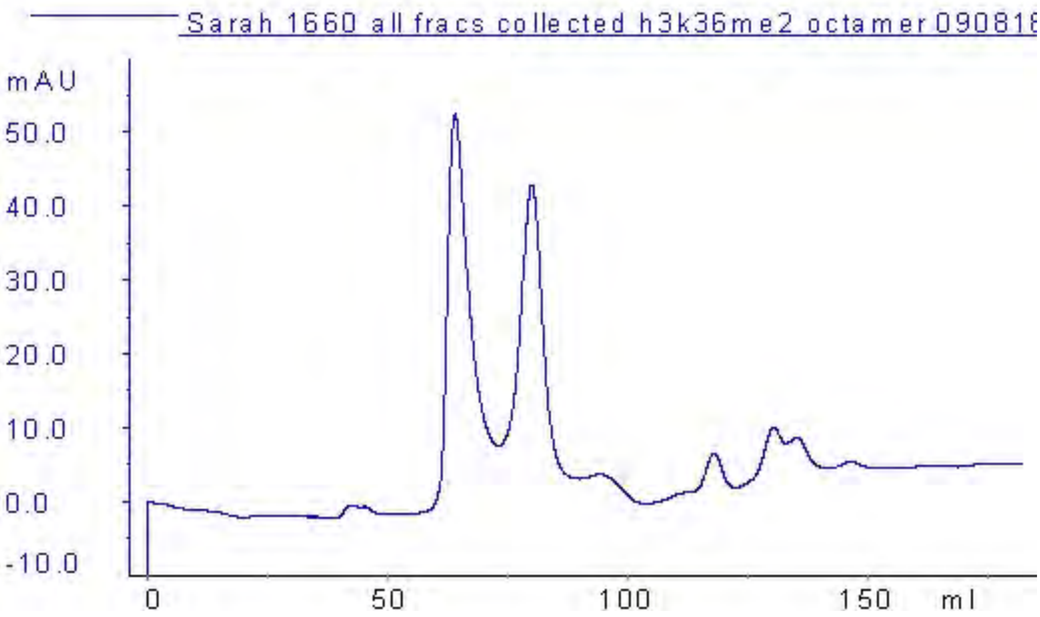


SDS-PAGE

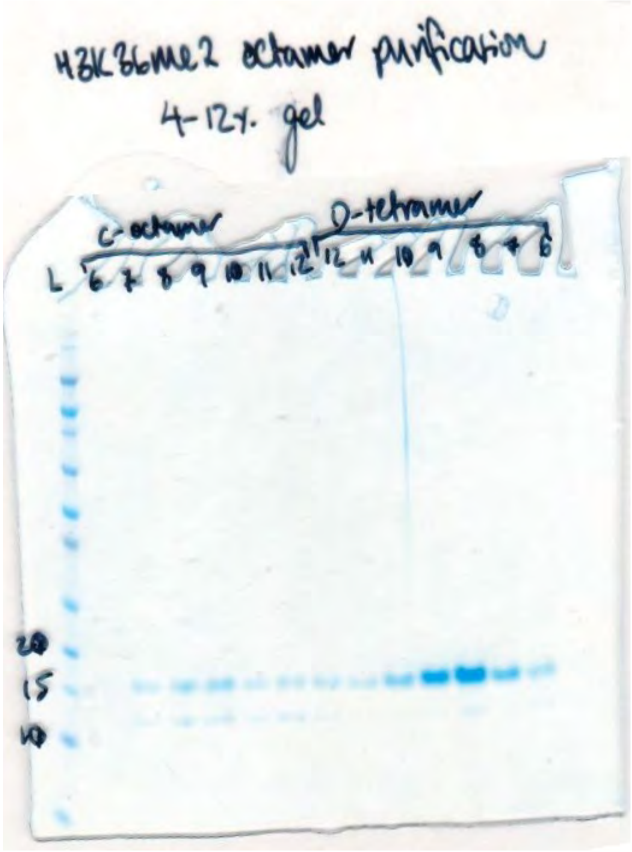


**Appendix Fig. 10:** Characterization data for H3K36me2 containing nucleosome

H3K36me3 octamer purification

UV trace from SEC

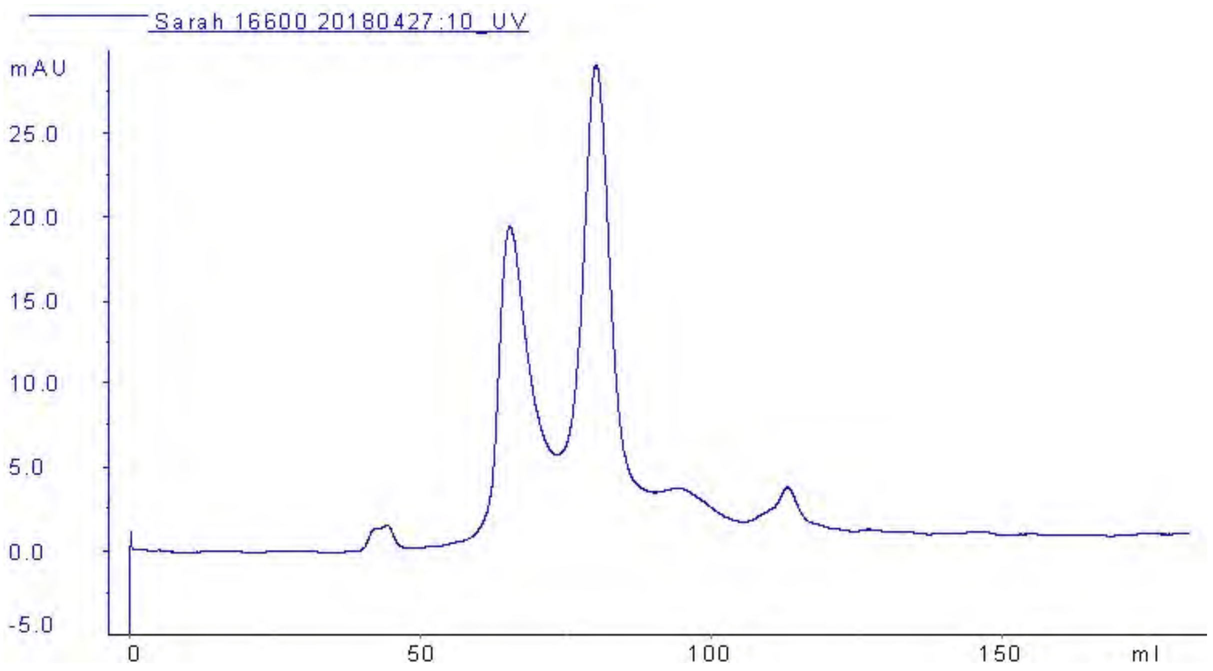


**Appendix Fig. 11:** Characterization data for H3K36me2 containing nucleosome.

**Additional data for Fig. 3C (Appendix Figs. 11-12)**

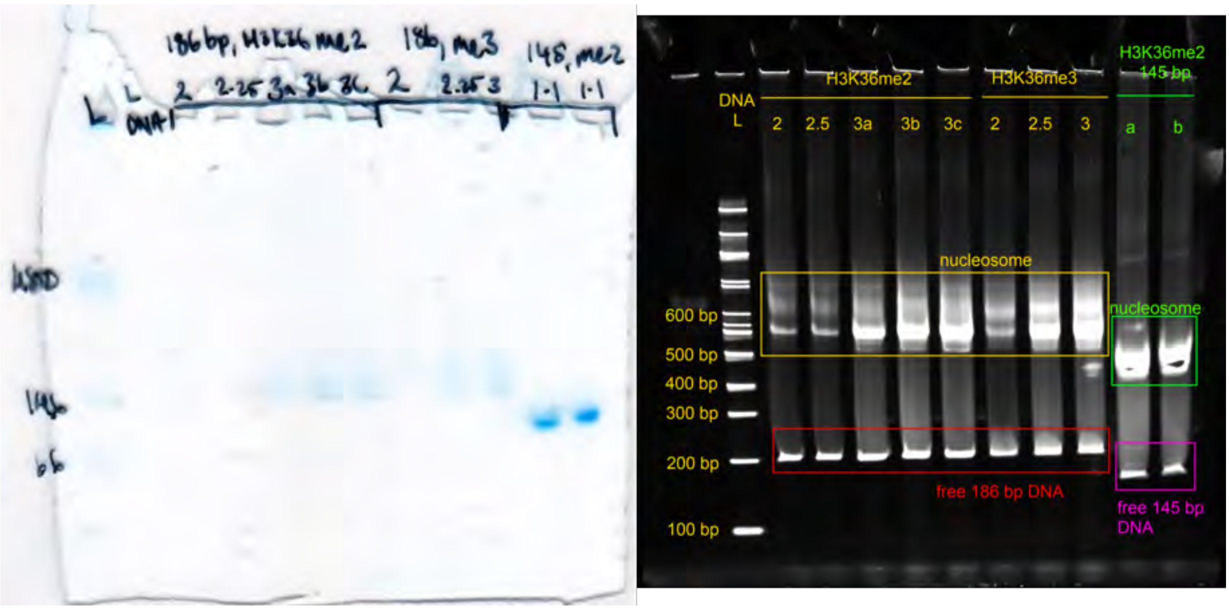


**Appendix Fig. 11:** 6% TBE gel of nucleosome reconstitution samples with me2/ me3 octamer and 186 bp DNA (2:1 to 3:1 DNA- octamer ratios)- left= Coomassie stain, right= SYBR gold.

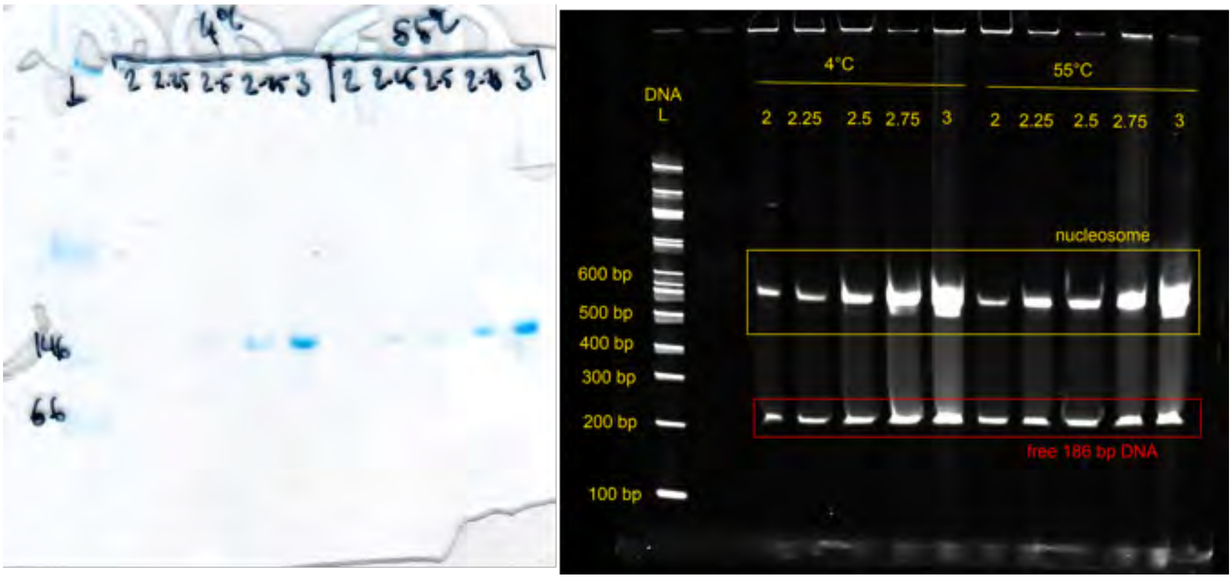


**Appendix Fig. 12:** 6% TBE gel of nucleosome reconstitution samples with WT octamer and 186 bp DNA (2:1 to 3:1 DNA- octamer ratios)- left= Coomassie stain, right= SYBR gold.

**Additional data for Fig. 4A (Appendix Figs. 13-15 and Appendix Table 2)**

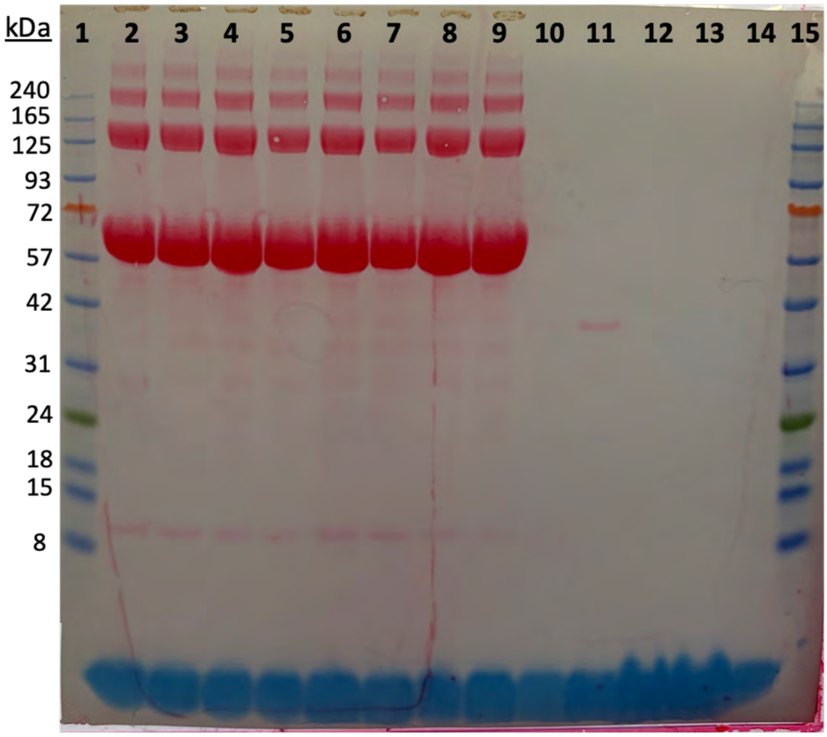


**Appendix Fig. 13:** Uncropped Ponceau-stained membrane for fig. 4A.

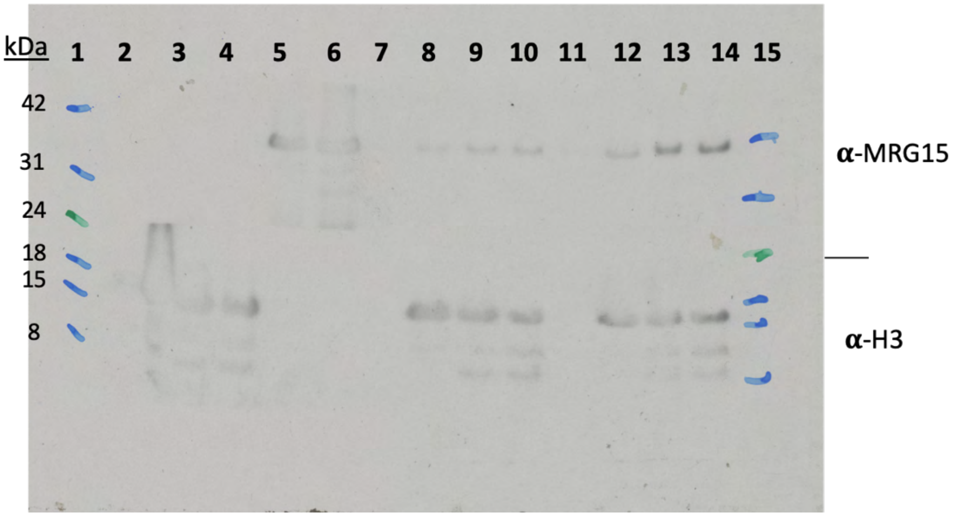


**Appendix Fig. 14:** Uncropped Western blot for fig. 4A- short exposure, used for MRG15 lane.

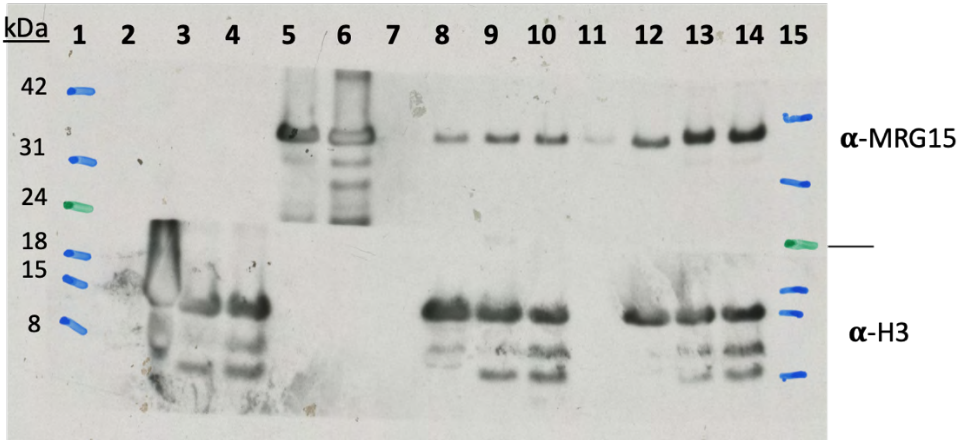


**Appendix Fig. 15:** Uncropped Western blot for Fig. 4a - long exposure, used for H3 lane.

| *Number* | *Description* |
|---|---|
| *1* | Ladder |
| *2* | WT nucleosome |
| *3* | Me2 nucleosome |
| *4* | Me3 nucleosome |
| *5* | hMRG15 W53A |
| *6* | hMRG15 WT |
| *7* | 186 bp DNA/ hMRG15 W53A |
| *8* | WT nuc/ hMRG15 W53A |
| *9* | Me2 nuc/ hMRG15 W53A |
| *10* | Me3 nuc/ hMRG15 W53A |
| *11* | 186 bp DNA/ hMRG15 WT |
| *12* | WT nuc/ hMRG15 WT |
| *13* | Me2 nuc/ hMRG15 WT |
| *14* | Me3 nuc/ hMRG15 WT |
| *15* | Ladder |

**Appendix Table 2.** Description of lanes in Fig. 4a blot (Appendix Figs. 13-15).

**Additional data for Fig. 4B (Appendix Fig. 16 and Appendix Table 3)**

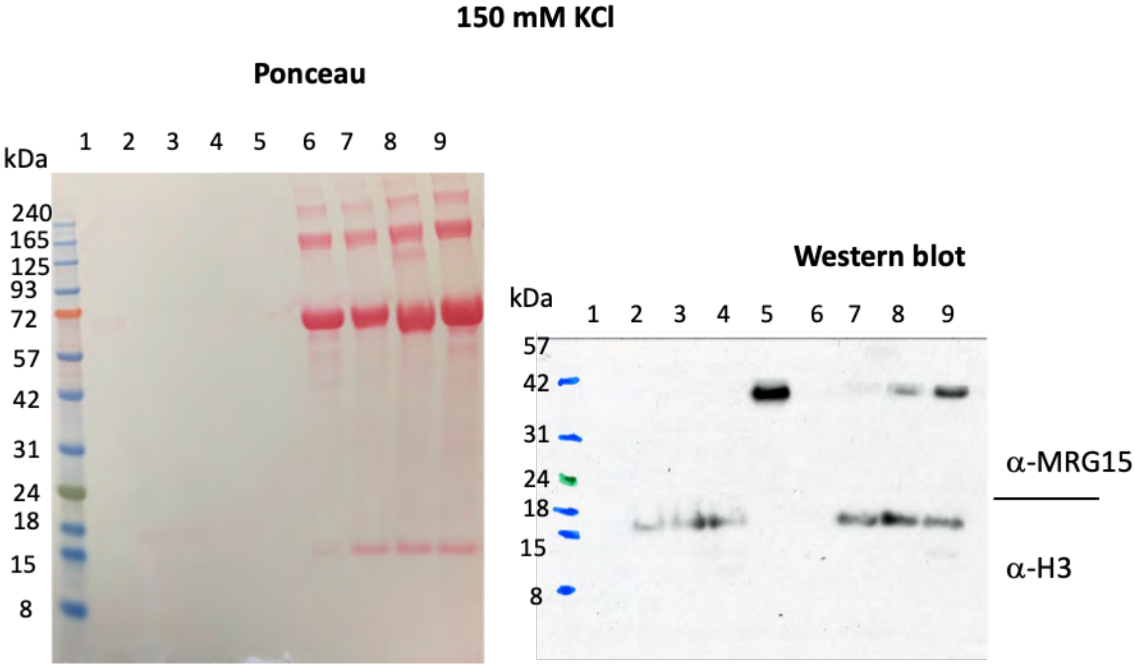


**Appendix Fig. 16:** Uncropped Ponceau-stained membrane (left) and developed Western blot (right) for pulldown in 150 mM KCl buffer with WT hMRG15 (Fig. 4b). See Appendix Table 3 for description of lanes.

| *Number* | *Description* |
|---|---|
| *1* | Ladder |
| *2* | 186 bp DNA |
| *3* | 186 bp WT nucleosome |
| *4* | 186 bp me2 nucleosome |
| *5* | 186 bp me3 nucleosome |
| *6* | hMRG15 |
| *7* | 186 bp DNA/hMRG15 (KCl) |
| *8* | WT nucleosome/hMRG15 (KCl) |
| *9* | me2 nucleosome/ hMRG15 (KCl) |
| *10* | Me3 nucleosome/ hMRG15 (KCl) |

**Appendix Table 3** Lanes in Appendix Fig. 16

**Additional data for Supplemental Fig. S1 (Appendix Fig. 17)**

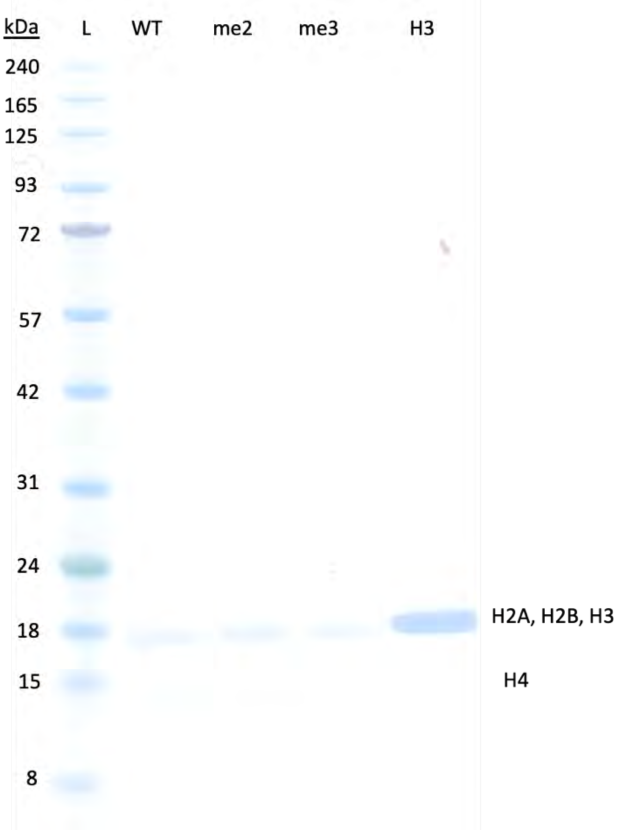


**Appendix Fig. 17:** Uncropped SDS-PAGE analysis of nucleosomes. H3 is loaded as control.

#### Additional data for Supplemental Fig. S2A (Appendix Fig. 18 and Appendix Table 4)

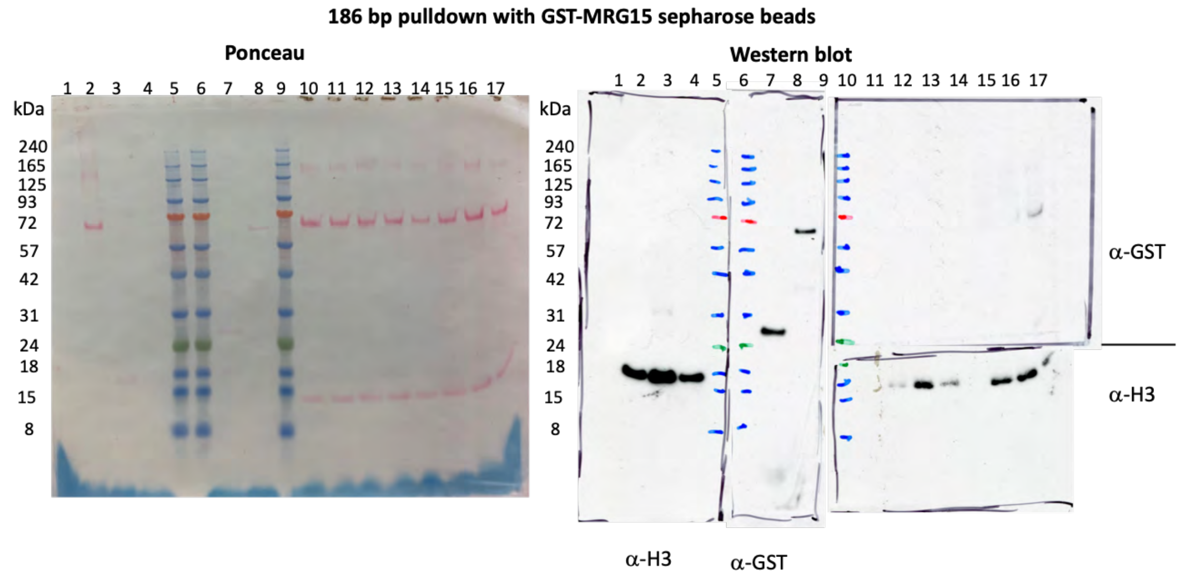


**Appendix Fig. 18:** Uncropped Ponceau-stained membrane (left) and developed Western blot (right) for pulldown of GST-MRG15 using 186 bp nucleosomes immobilised on Streptavidin beads (Fig. S3A). See Appendix Table 5 for description of lanes.

| *Number* | *Description* |
|---|---|
| *1* | 186 bp DNA |
| *2* | 186 bp WT nucleosome |
| *3* | 186 bp me2 nucleosome |
| *4* | 186 bp me3nucleosome |
| *5* | Ladder |
| *6* | Ladder |
| *7* | GST |
| *8* | GST-MRG15 |
| *9* | Ladder |
| *10* | 186 bp DNA/GST |
| *11* | WT nucleosome/GST |
| *12* | me2 nucleosome/GST |
| *13* | Me3 nucleosome/GST |
| *14* | 186 bp DNA/GST-MRG15 |
| *15* | WT nucleosome/GST-MRG15 |
| *16* | me2 nucleosome/GST-MRG15 |
| *17* | Me3 nucleosome/GST-MRG15 |

**Appendix Table 4:** Lanes in Appendix Fig. 18

#### Additional data for Supplemental Fig. S2B (Appendix Fig. 19 and Appendix Table 5)

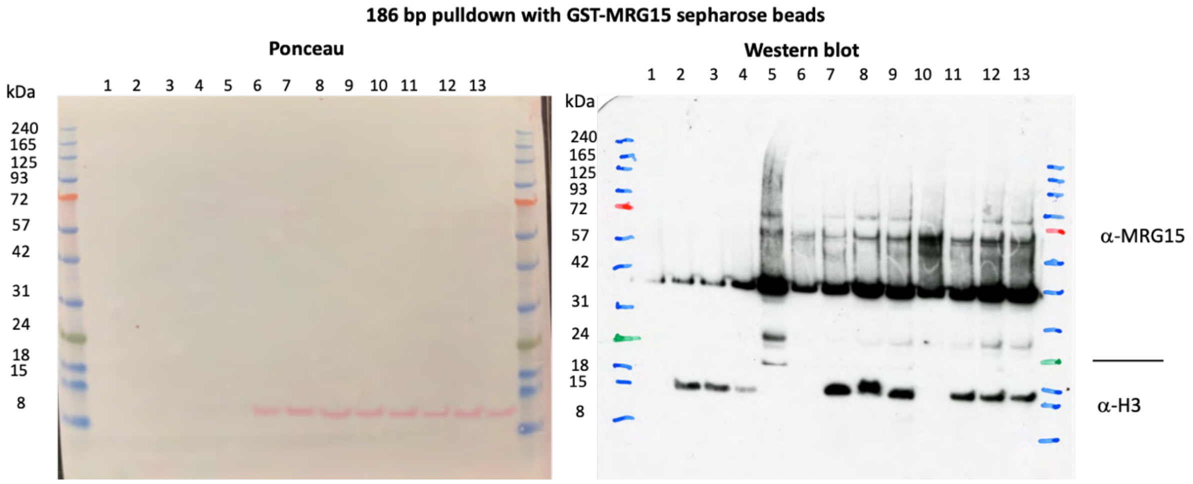


**Appendix Fig. 19:** Uncropped Ponceau-stained membrane (left) and developed Western blot (right) for pulldown to screen 250 mM KCl buffer vs 150 mM NaCl buffer (Fig. S3B). See table S6 for description of lanes.

| *Number* | *Description* |
|---|---|
| *1* | Ladder |
| *2* | 186 bp DNA |
| *3* | 186 bp WT nucleosome |
| *4* | 186 bp me2 nucleosome |
| *5* | 186 bp me3 nucleosome |
| *6* | hMRG15 |
| *7* | 186 bp DNA/hMRG15 (KCl) |
| *8* | WT nucleosome/hMRG15 (KCl) |
| *9* | me2 nucleosome/ hMRG15 (KCl) |
| *10* | Me3 nucleosome/ hMRG15 (KCl) |
| *11* | 186 bp DNA/hMRG15 (NaCl) |
| *12* | WT nucleosome/hMRG15 (NaCl) |
| *13* | me2 nucleosome/ hMRG15 (NaCl) |
| *14* | Me3 nucleosome/ hMRG15 (NaCl) |
| *15* | Ladder |

**Appendix Table 5:** Lanes in Appendix Fig. 19.

**Additional data for Supplemental Fig. S3 (Appendix Fig. 20 and Appendix Table 5)**

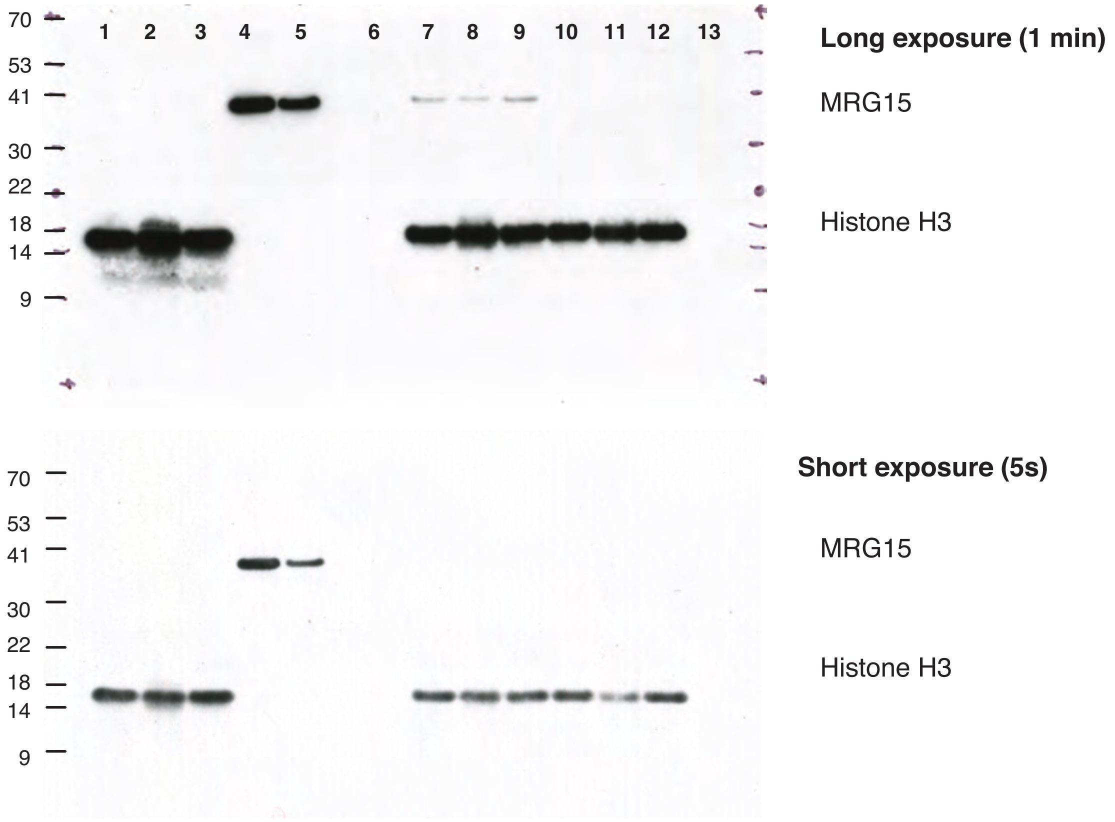


**Appendix Fig. 20:** Uncropped images of Western blot (right) for pulldown of WT or W53A MRG15 using EpiCypher nucleosomes, assembled with the 147 bp 601 DNA with 5' biotinylation, immobilised on Streptavidin beads (Fig. S1). ). See Appendix Table 4 for description of lanes.

| *Number* | *Description* |
|---|---|
| *1* | WT EpiCypher nucleosome |
| *2* | me2 EpiCypher nucleosome |
| *3* | me3 EpiCypher nucleosome |
| *4* | MRG15 WT |
| *5* | MRG15 W53A |
| *6* | 186 bp DNA/MRG15 WT |
| *7* | WT EpiCypher nucleosome/ MRG15 WT |
| *8* | me3 EpiCypher nucleosome/ MRG15 WT |
| *9* | me2 EpiCypher nucleosome/ MRG15 WT |
| *10* | WT EpiCypher nucleosome/ MRG15 W53A |
| *11* | me2 EpiCypher nucleosome/ MRG15 W53A |
| *12* | me3 EpiCypher nucleosome/ MRG15 W53A |
| *13* | 186 bp DNA/ MRG15 W53A |

**Appendix Table 5:** Lanes in Appendix Fig. 20

### Appendix Part II- Additional Data for Protein and DNA Preparation

##### Additional data for histone purification (Appendix Figs. 21-36)

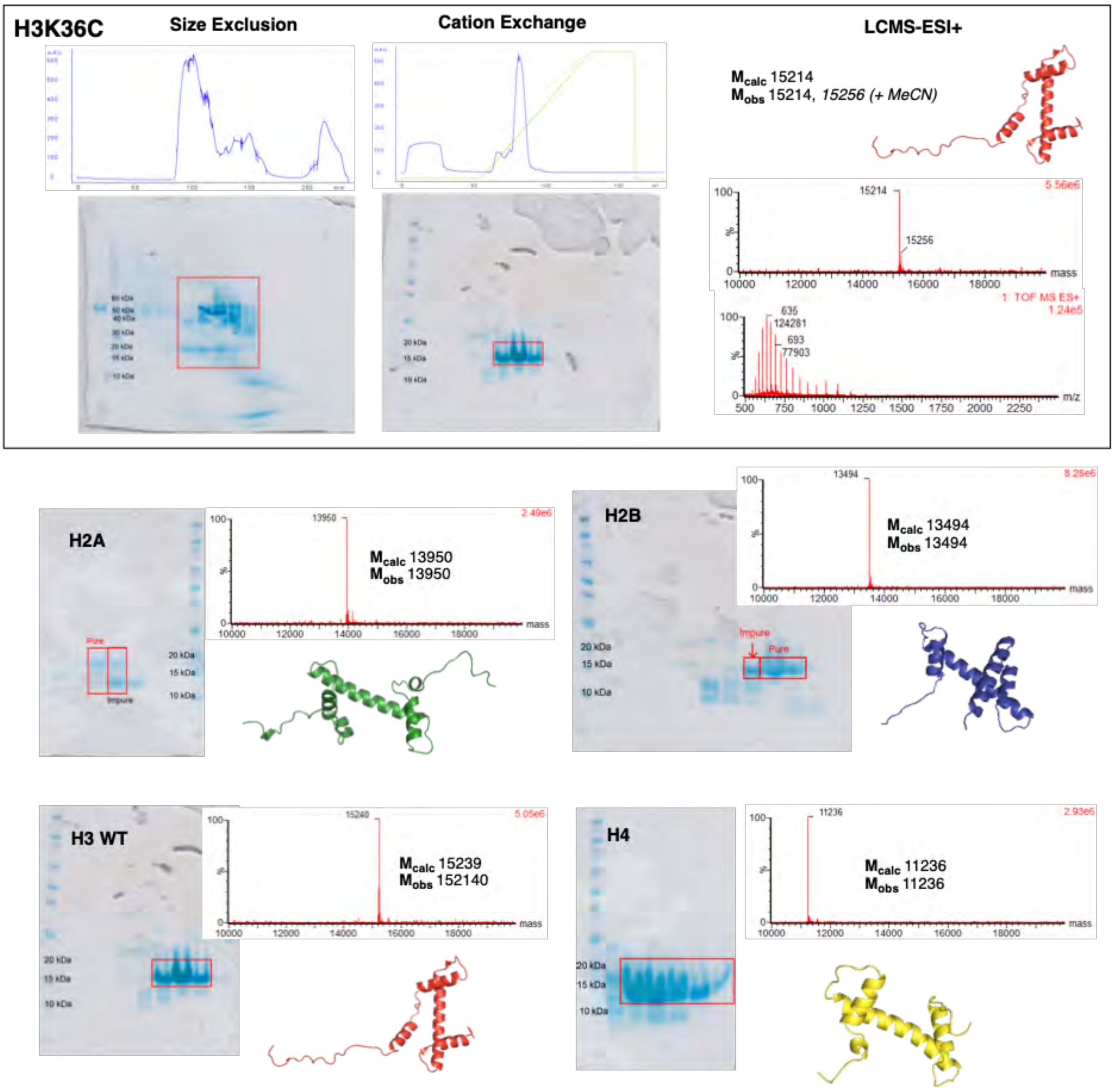


**Appendix Fig. 21:** Expression and purification of the five histone variants**.** Full characterization data i.e. size exclusion and cation exchange FPLC traces, SDS-PAGE analysis, and LCMS-ESI+ ion series and deconvoluted spectrum are shown as an example for H3K36C. Final SDS-PAGE from cation exchange step and deconvoluted LCMS-ESI+ spectra are shown for all other variants.

**H3 WT**

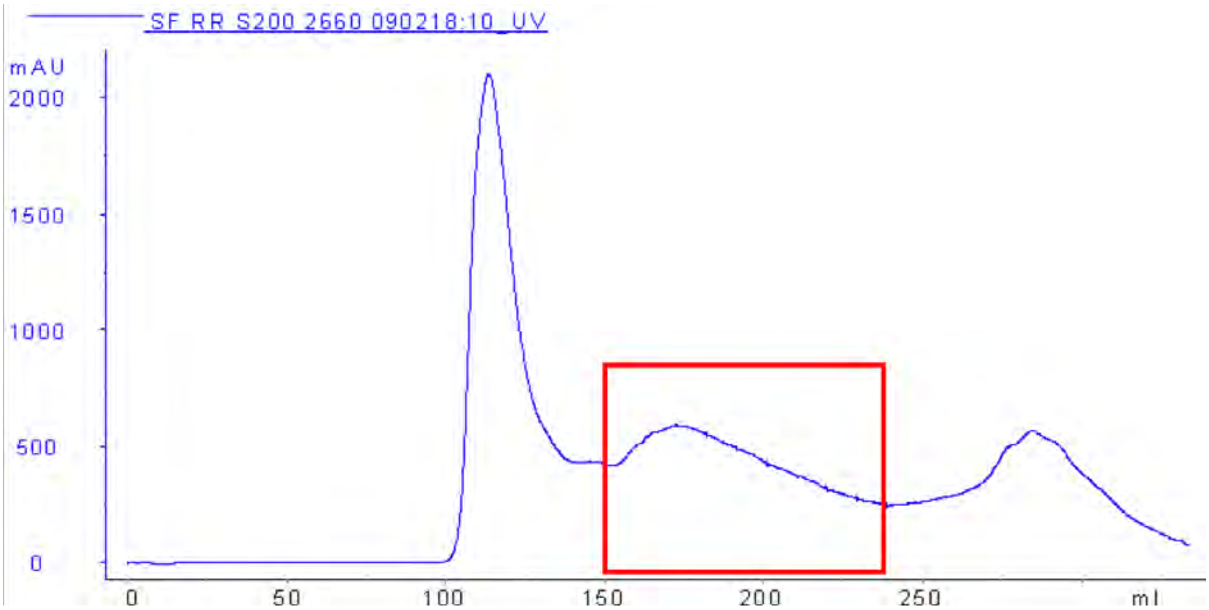


SDS-PAGE

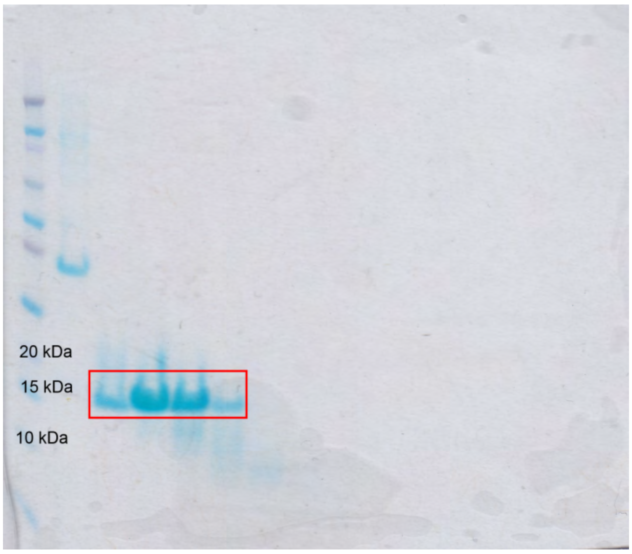


Fractions highlighted in red correspond to those containing H3 WT

**Appendix Fig. 22:** Characterization of H3 WT after SEC purification

UV trace from Hitrap

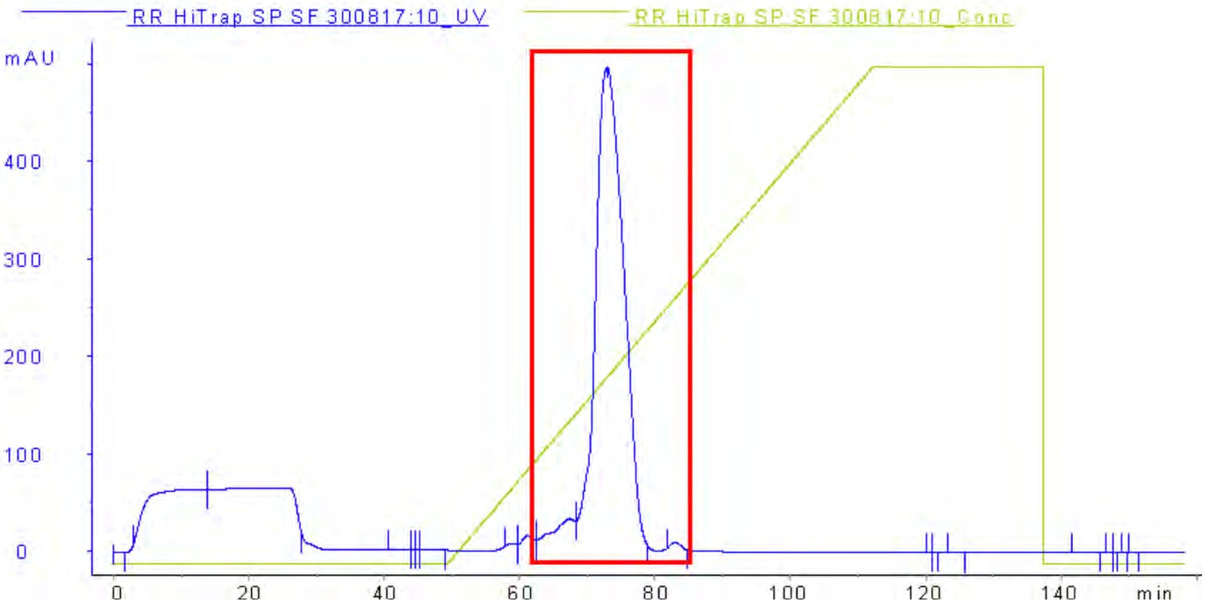


SDS-PAGE

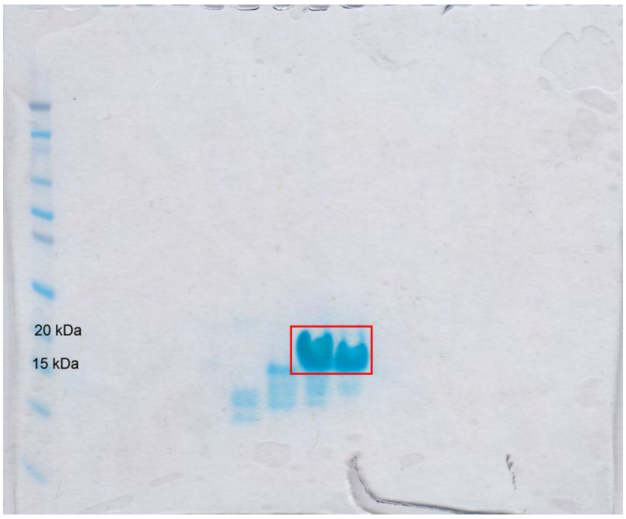


Fractions highlighted in red correspond to those containing H3 WT

**Appendix Fig. 23:** Characterization of H3 WT after Hitrap purification- UV trace and SDS-PAGE.

LCMS-ESI+ characterization: chromatogram

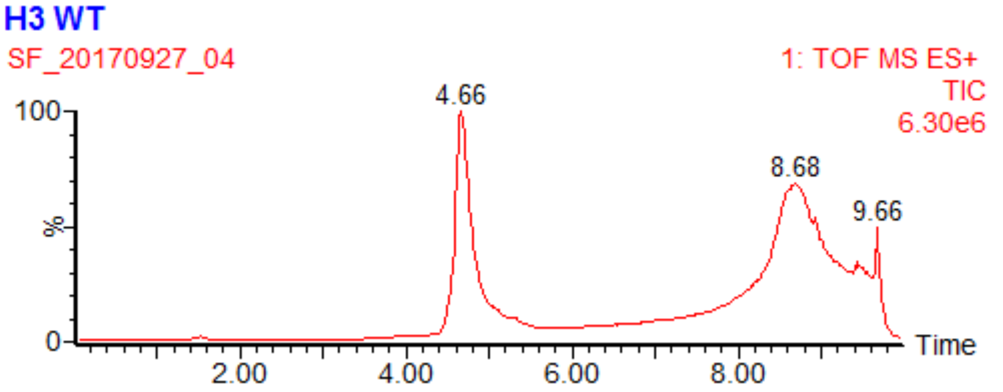


Ion series

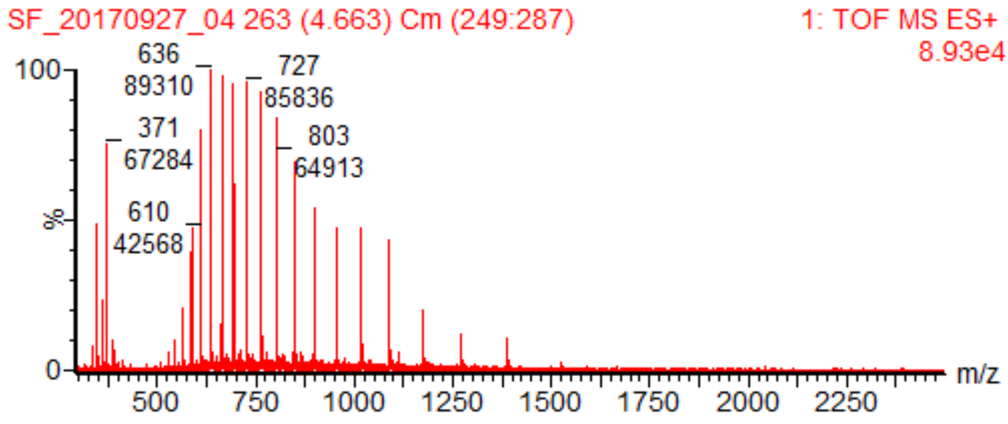


Deconvoluted mass spectrum

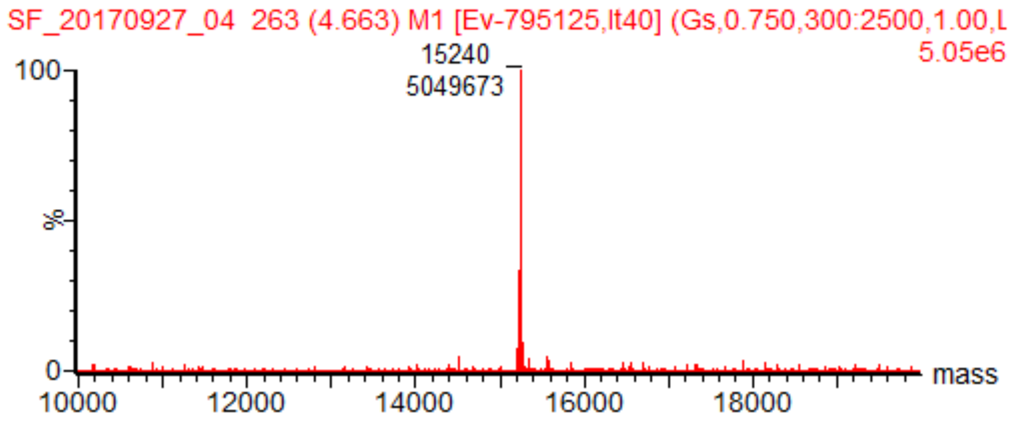


MS-ESI+: Expected 15238.80 Da ; Found 15240 Da

(Note that M from start codon is not present in final protein)

**Appendix Fig. 24:** Characterization of H3 WT after Hitrap purification- LCMS-ESI+

**Yield = 101 mg (42 mg/ L of culture)**

**H3K36C**

UV trace from SEC

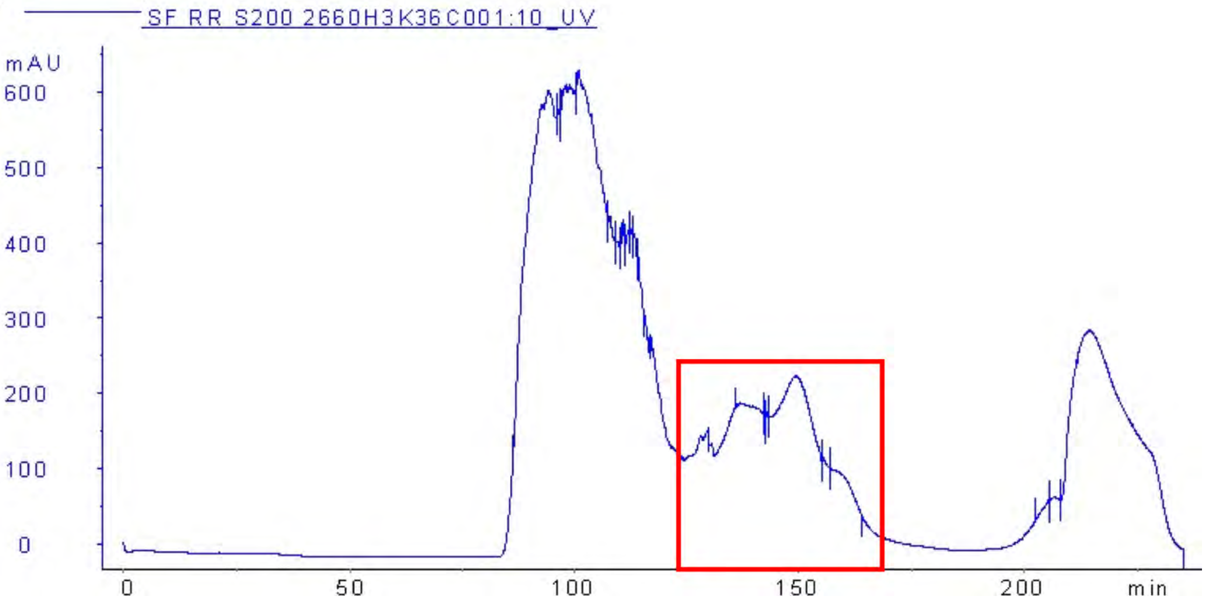


SDS-PAGE

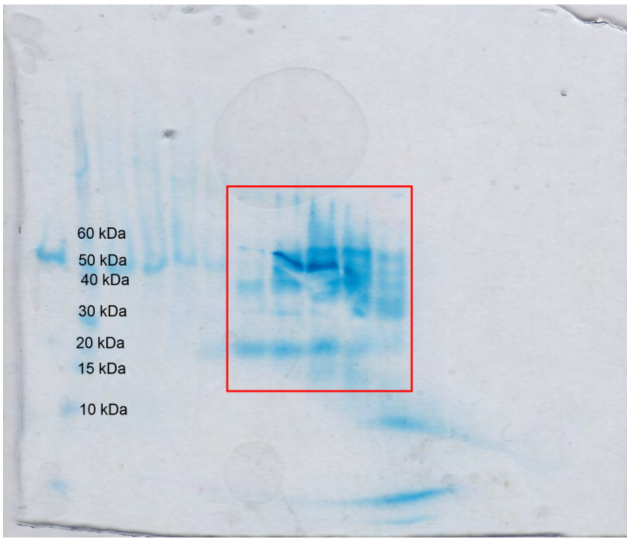


Fractions highlighted in red correspond to those containing H3K36C

**Appendix Fig. 25:** Characterization of H3K36C after SEC purification

UV trace from Hitrap

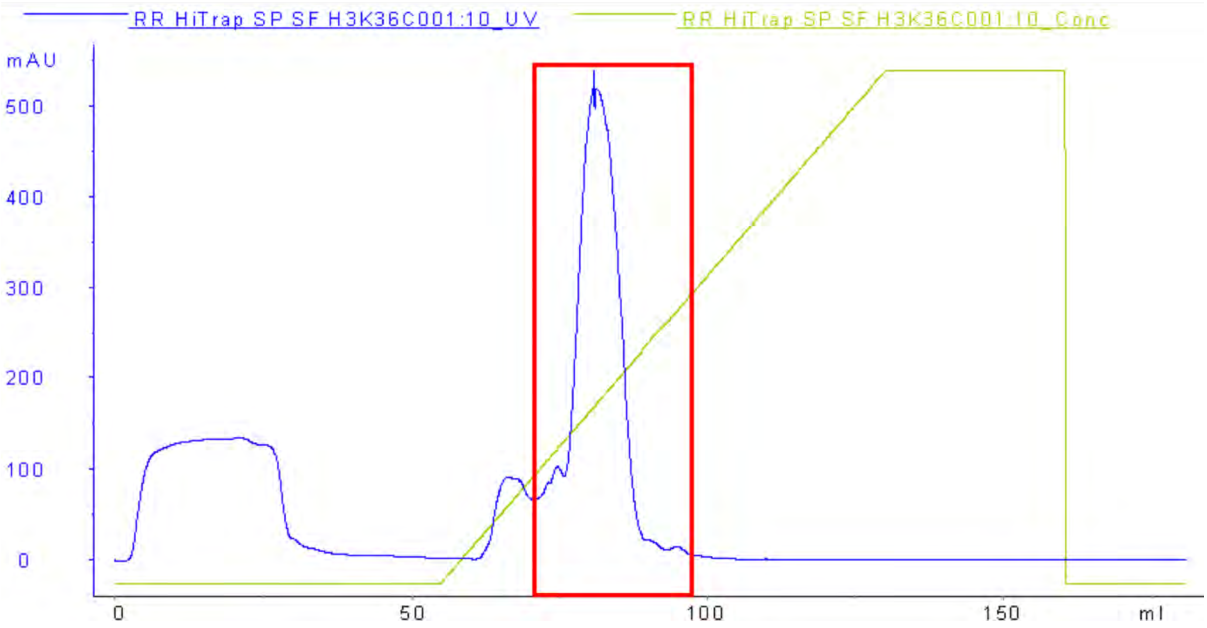


SDS-PAGE

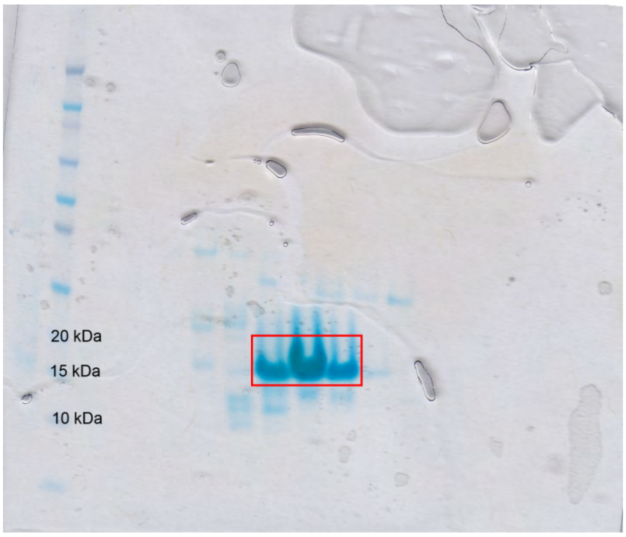


Fractions highlighted in red correspond to those containing H3K36C

**Appendix Fig. 26:** Characterization of HK36C after Hitrap purification- UV trace and SDS-PAGE.

LCMS-ESI+ characterization: Chromatogram

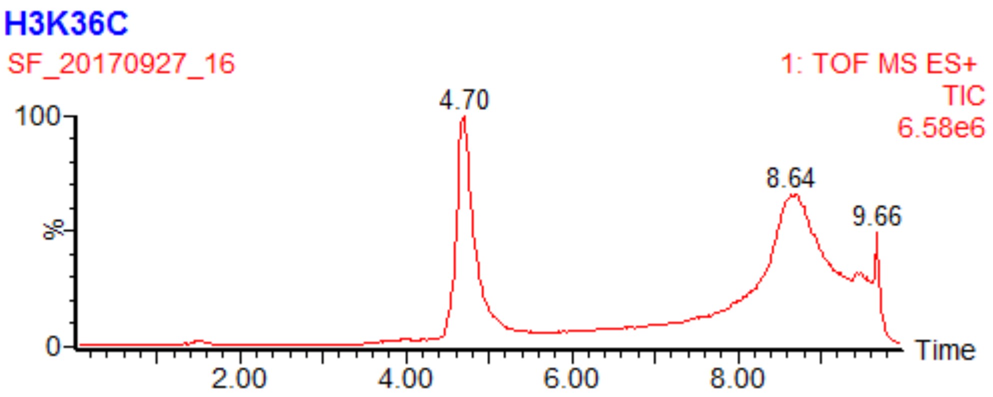


Ion series

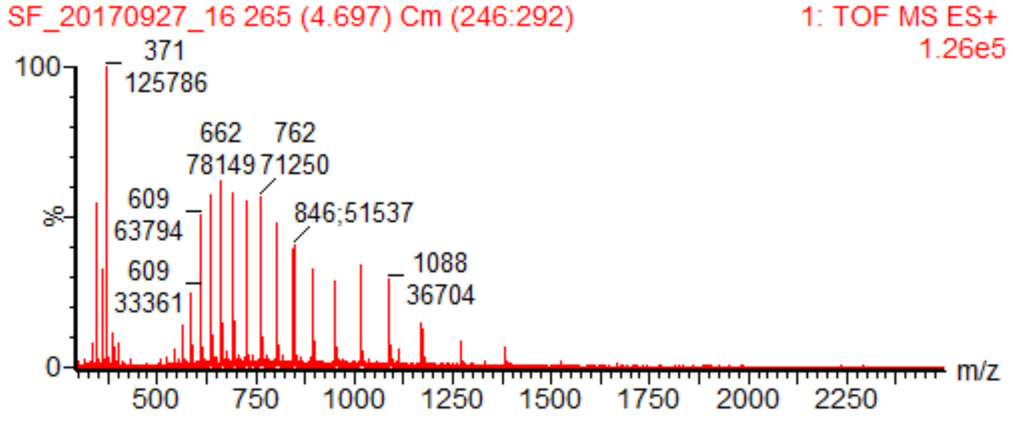


Deconvoluted mass spectrum

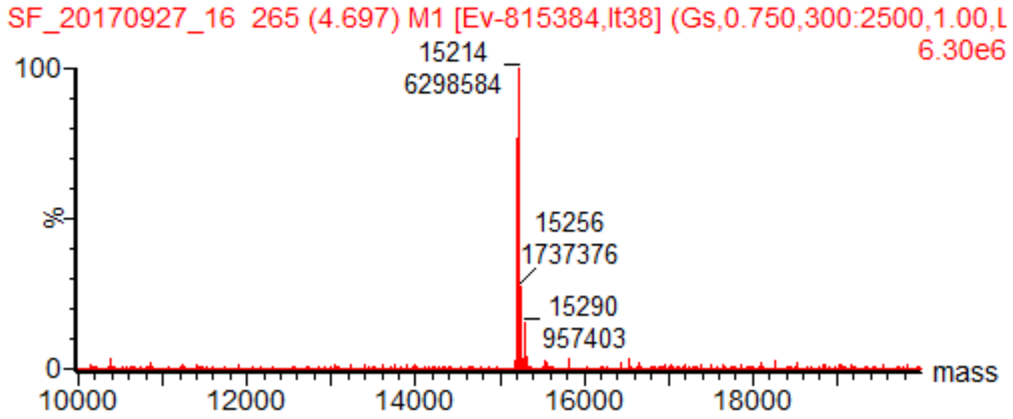


MS-ESI+: Expected 15213.76 Da ; found 15214 Da (15256 peak= +42= acetonitrile adduct)

(Note that M from start codon is not present in final protein)

**Appendix Fig. 27:** Characterization of H3K36C after Hitrap purification- LCMS-ESI+

**Yield = 66 mg (27.5 mg/ L of culture)**

**H2B**

UV trace from SEC

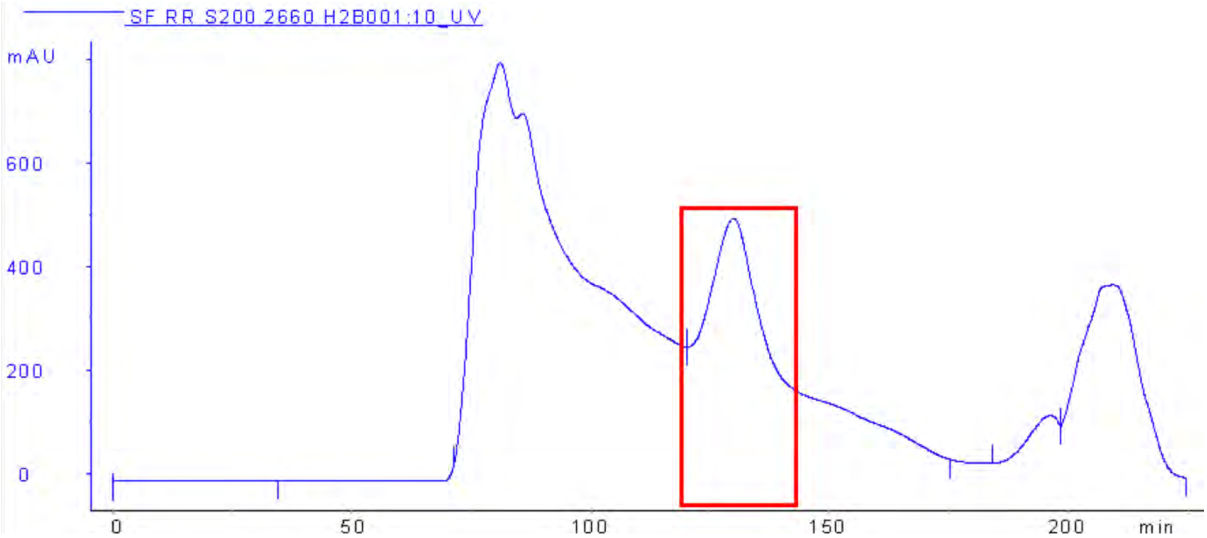


SDS-PAGE

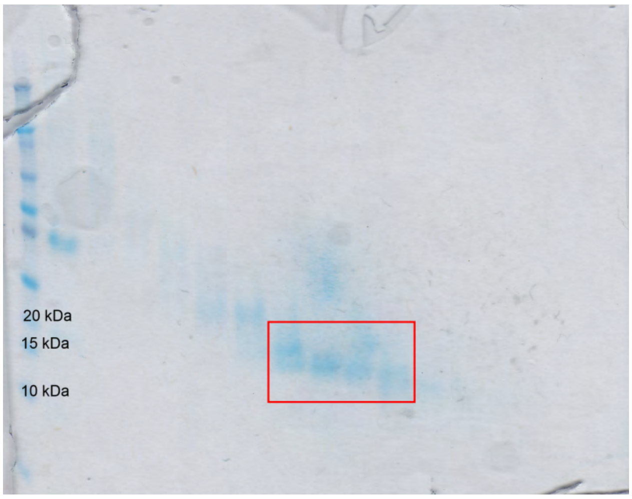


Fractions highlighted in red correspond to those containing H2B

**Appendix Fig. 28:** Characterization of H2B after SEC purification

UV trace from Hitrap

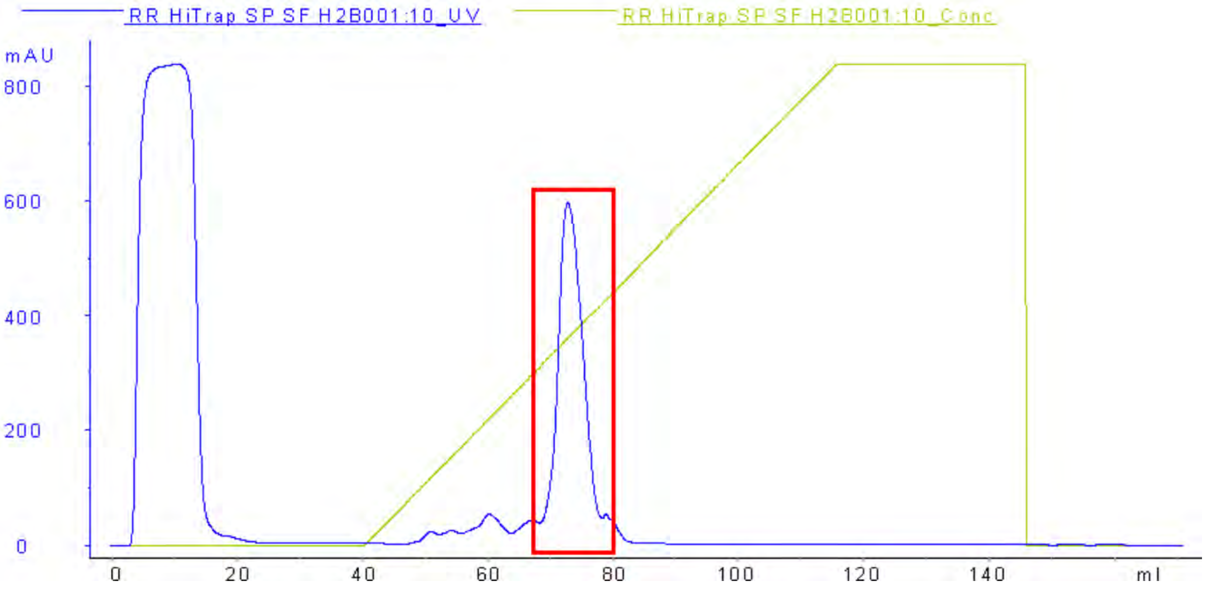


SDS-PAGE

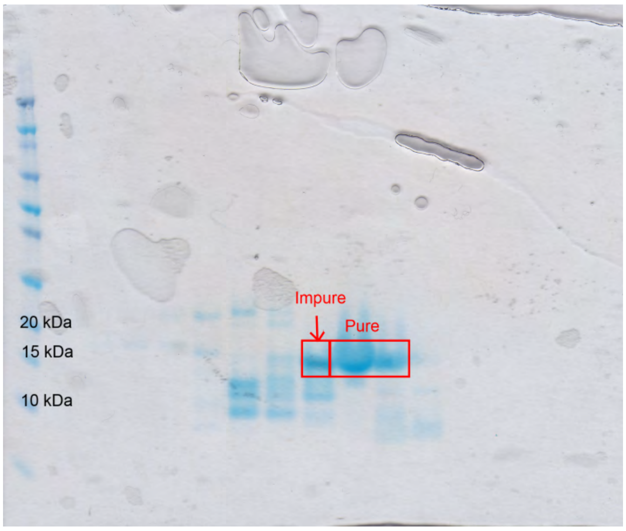


Fractions highlighted in red correspond to those containing H2B

**Appendix Fig. 29:** Characterization of H2B after Hitrap purification- UV trace and SDS-PAGE.

LCMS-ESI+ characterization- Chromatogram

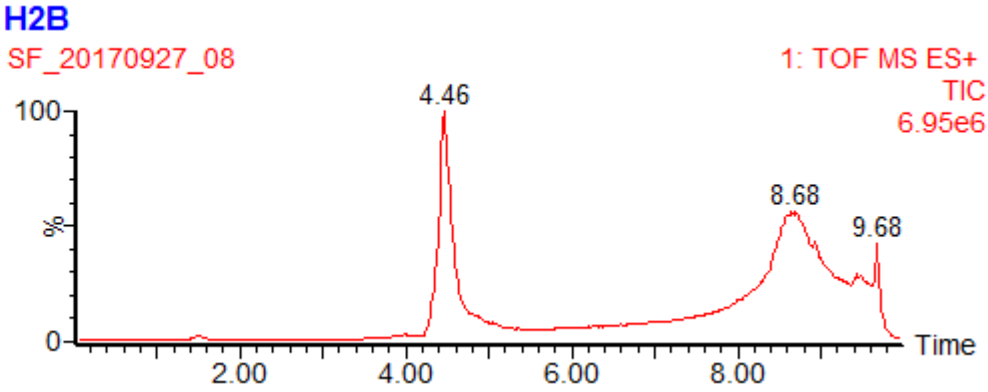


Ion series

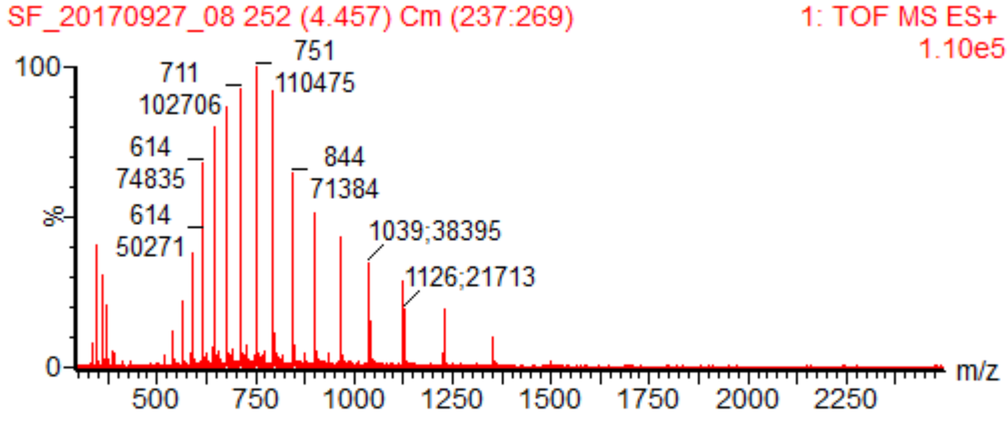


Deconvoluted mass spectrum

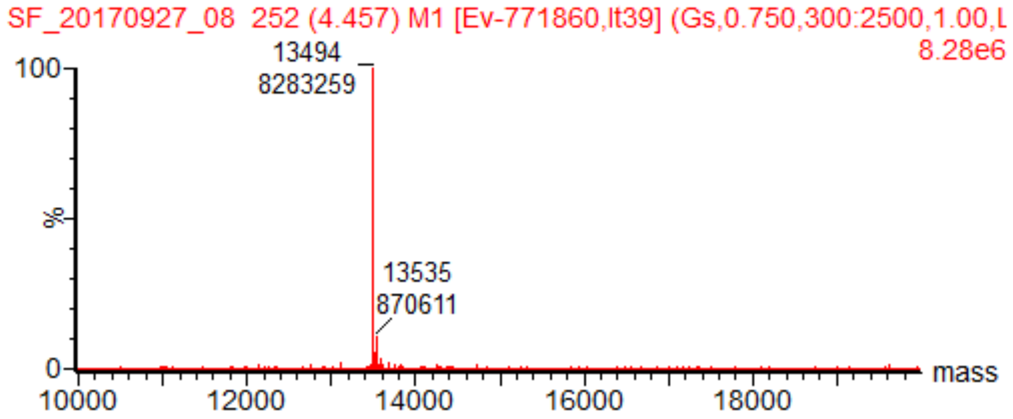


MS-ESI+: Expected 13493.68 Da ; Found 13494 Da (Note that M from start codon is not present in final protein)

**Appendix Fig. 30:** Characterization of H2B after Hitrap purification- LCMS-ESI+

**Yield = 60 mg (25 mg/ L of culture)**

H2A

UV trace from SEC

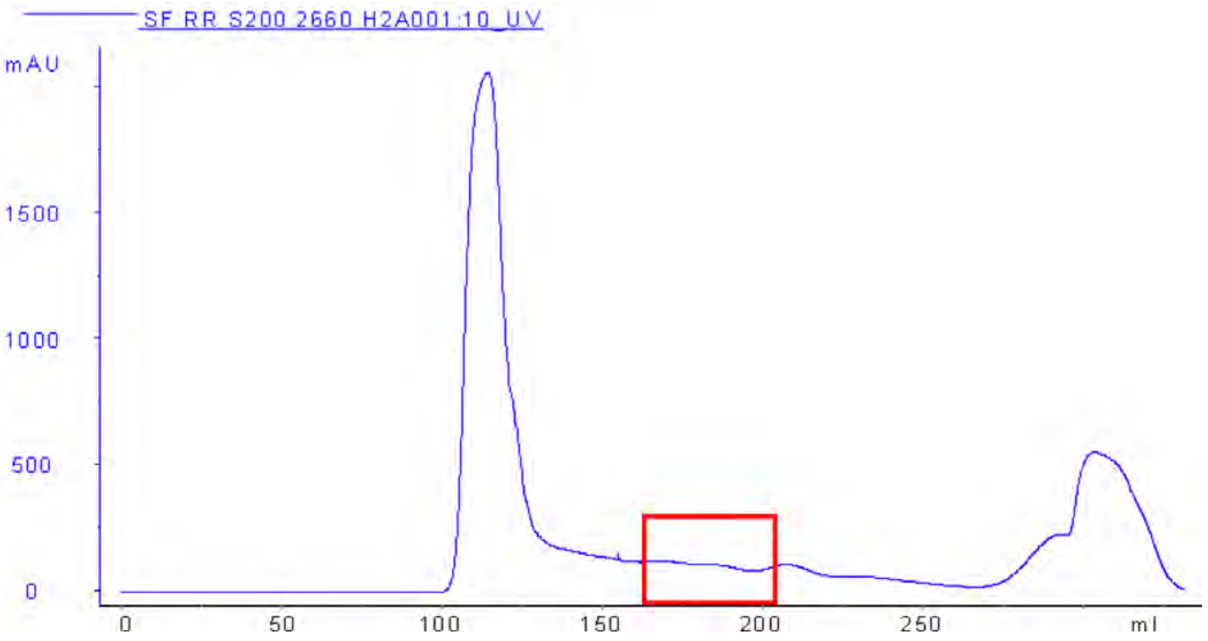


SDS-PAGE

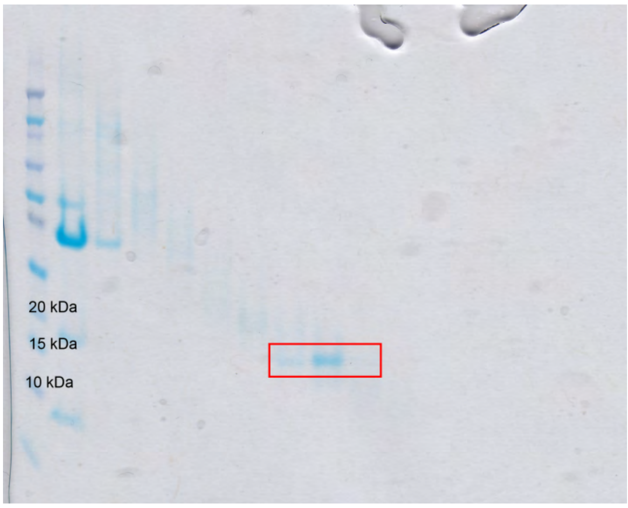


Fractions highlighted in red correspond to those containing H2A.

Note: H2A yield was unusually low. This was because FPLC was left to run overnight, but stopped before H2A had eluted; as a result, H2A had time to diffuse through the column and was lost.

**Appendix Fig. 31:** Characterization of H2A after SEC purification

UV trace from Hitrap

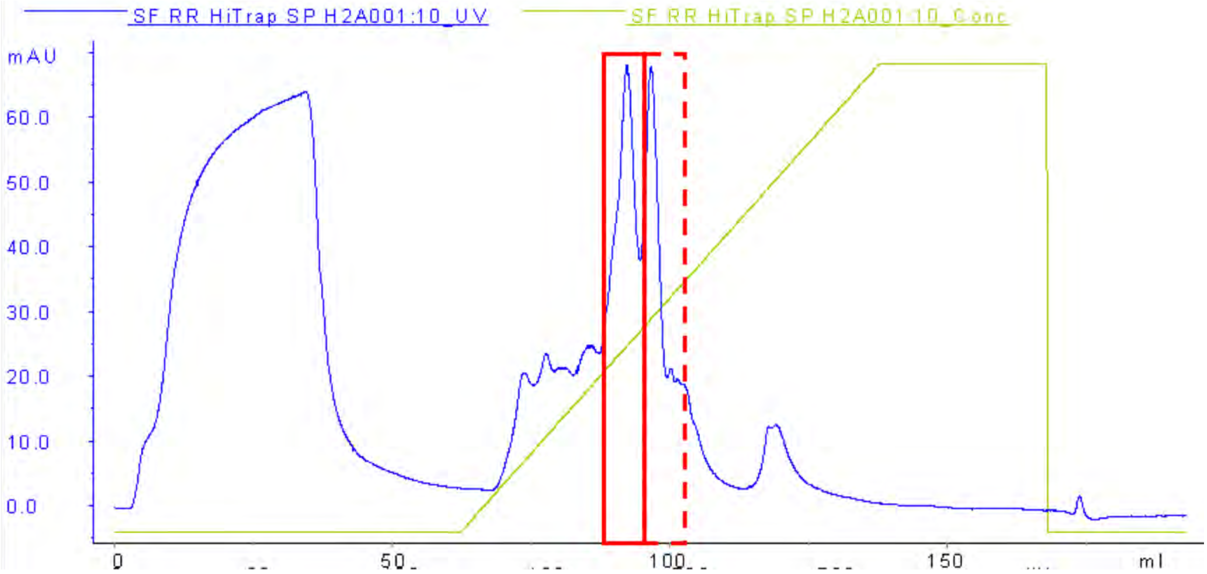


SDS-PAGE

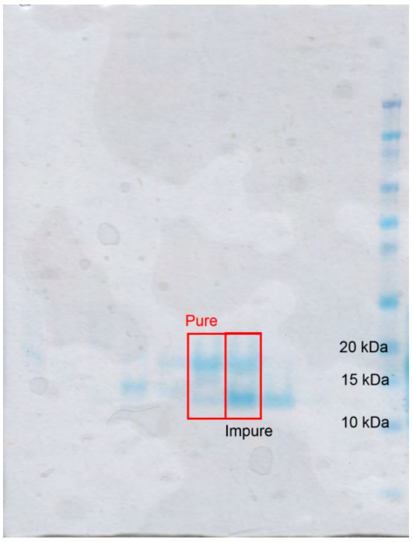


Fractions highlighted in red correspond to those containing H2A

**Appendix Fig. 32:** Characterization of H2A after Hitrap purification- UV trace and SDS-PAGE.

LCMS-ESI+ characterization- Chromatogram

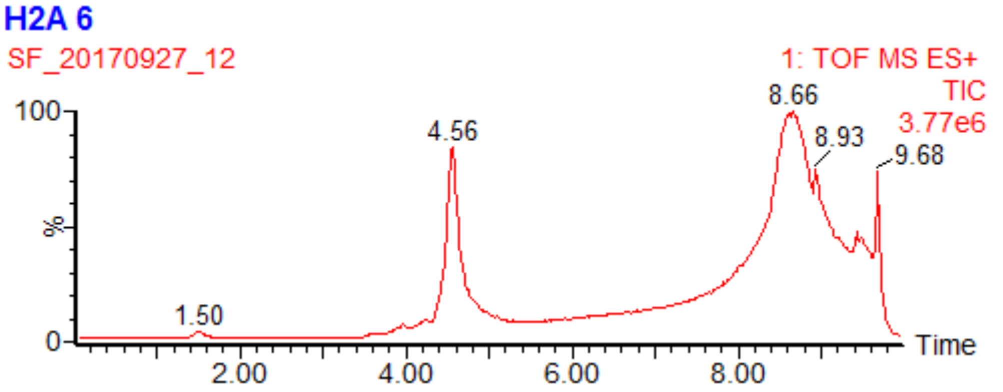


Ion series

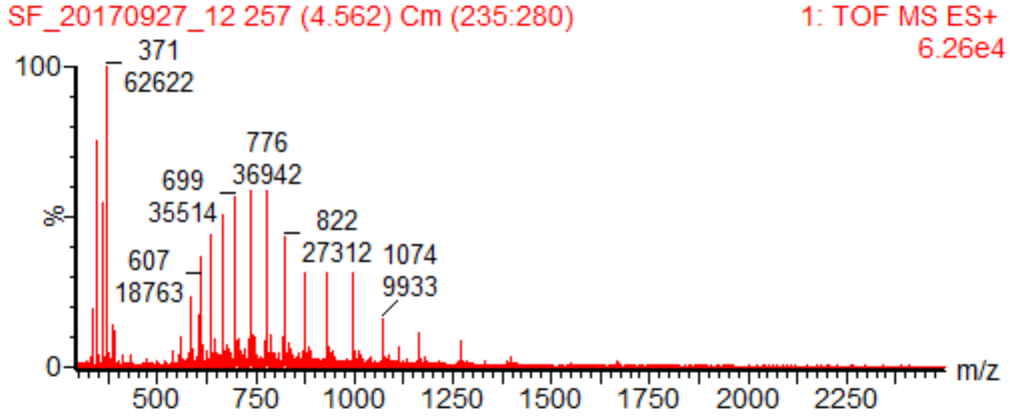


Deconvoluted mass spectrum

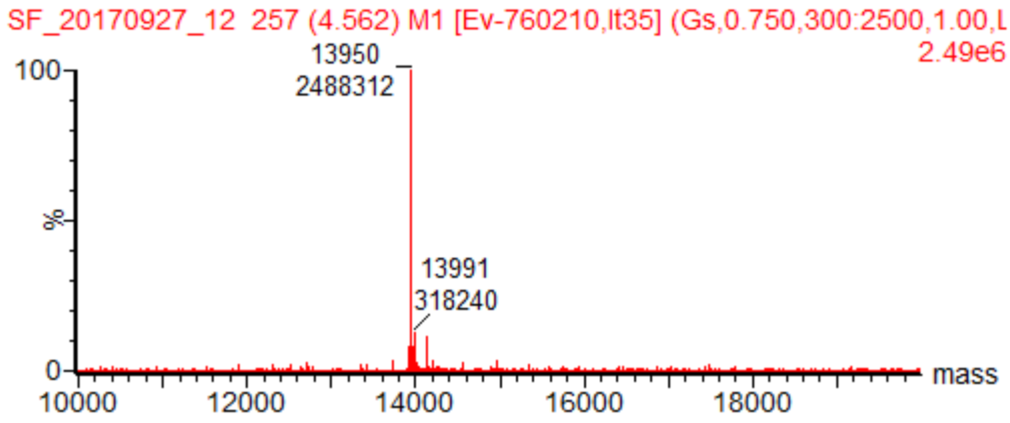


MS-ESI+: Expected 13950.20 Da ; Found 13950 Da.

(Note that M from start codon is not present in final protein)

**Appendix Fig. 33:** Characterization of H2A after Hitrap purification- LCMS-ESI+

**Yield = 13 mg (5.4 mg/ L of culture).** Low yield is due to loss on SEC column.

**H4**

SEC UV trace

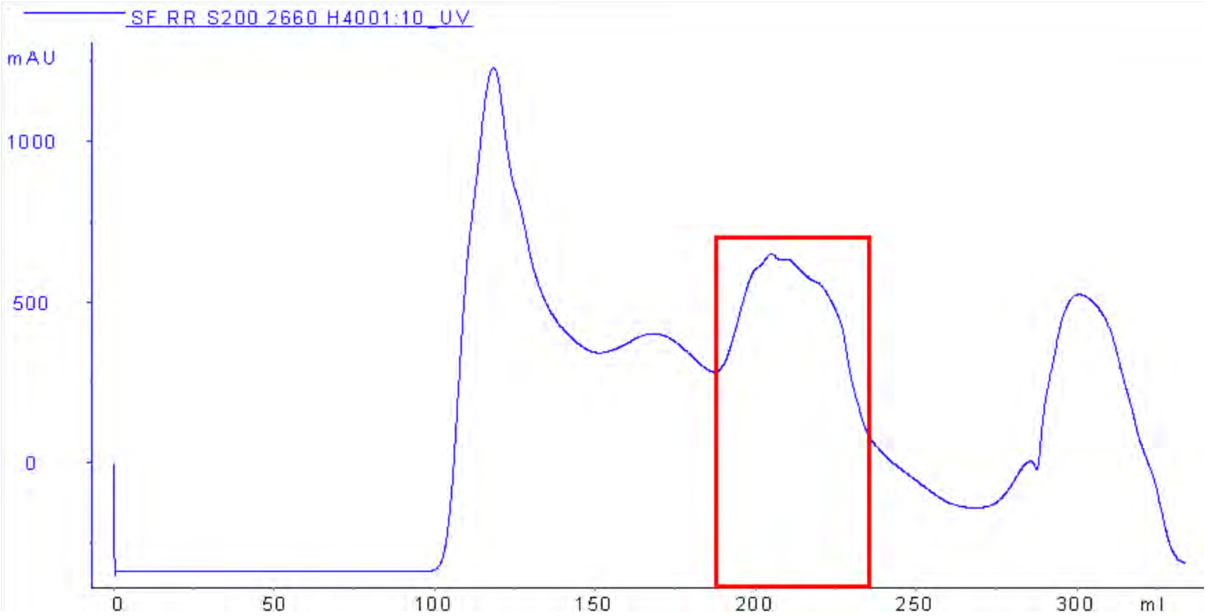


SDS-PAGE

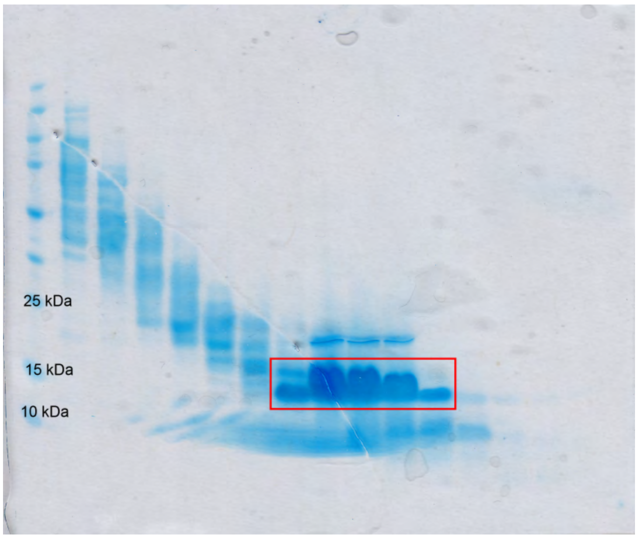


Fractions highlighted in red correspond to those containing H4

**Appendix Fig. 34:** Characterization of H4 after SEC purification

UV trace from Hitrap

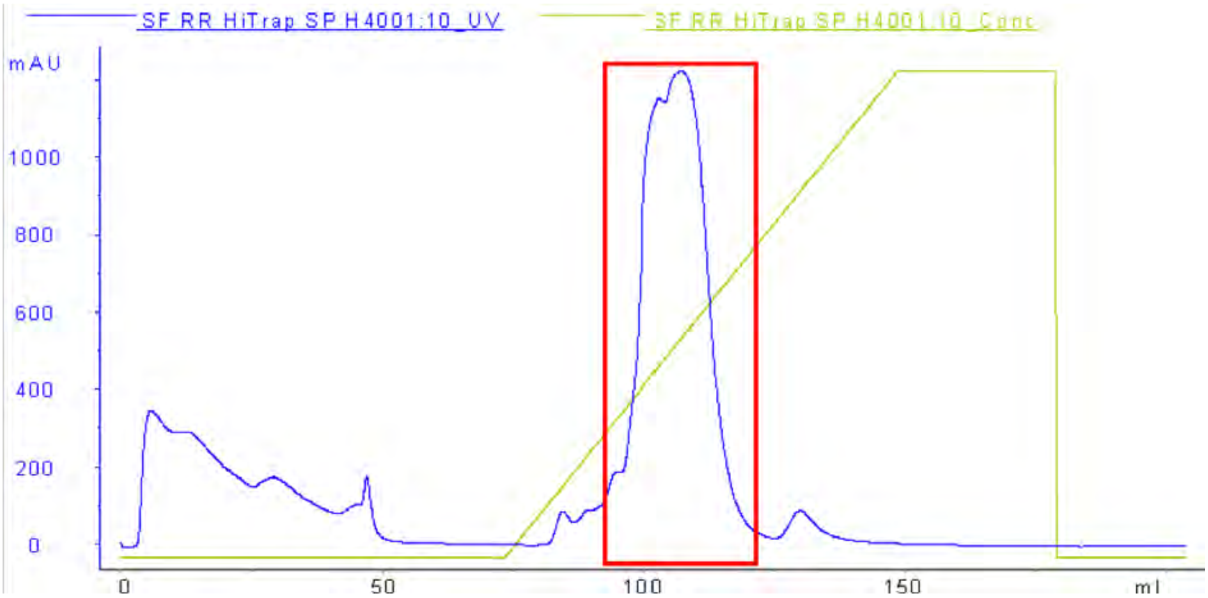


SDS-PAGE

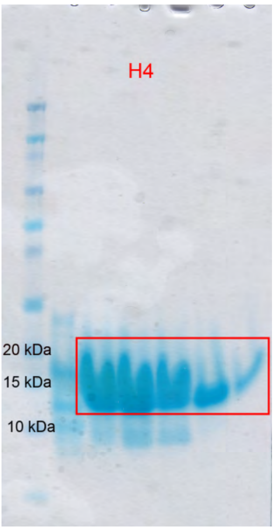


Fractions highlighted in red correspond to those containing H4

**Appendix Fig. 35:** Characterization of H4 after Hitrap purification- UV trace and SDS-PAGE.

LCMS-ESI+ characterization- Chromatogram

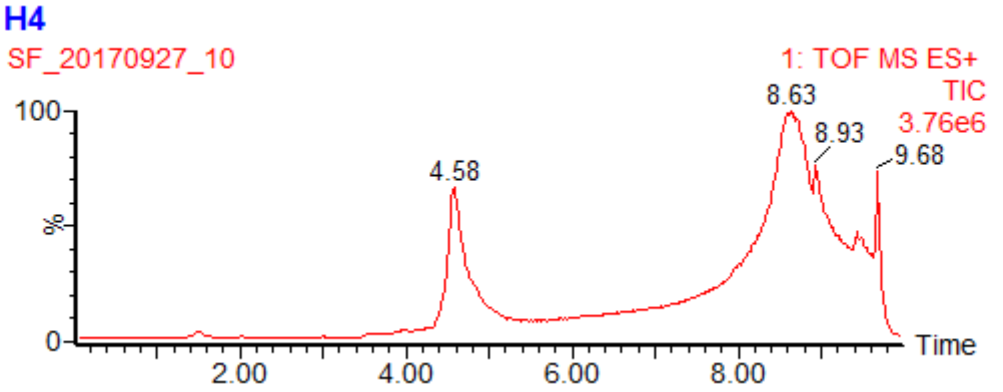


Ion series

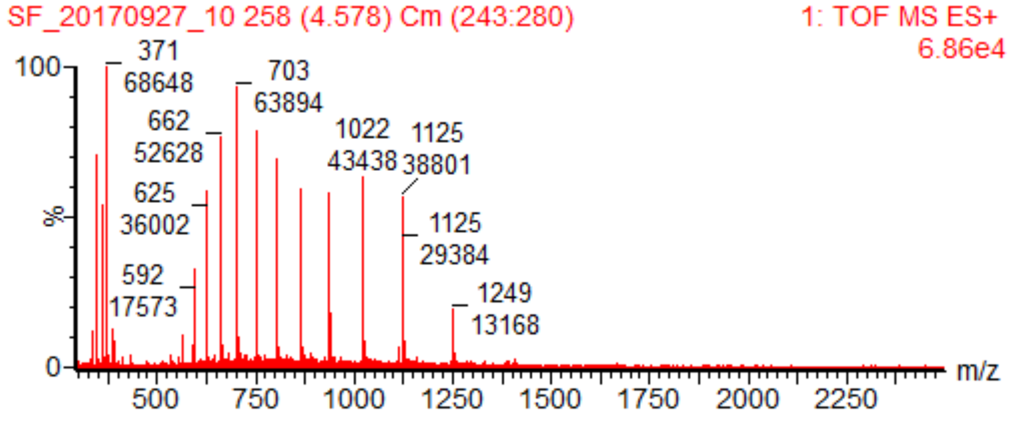


Deconvoluted mass spectrum

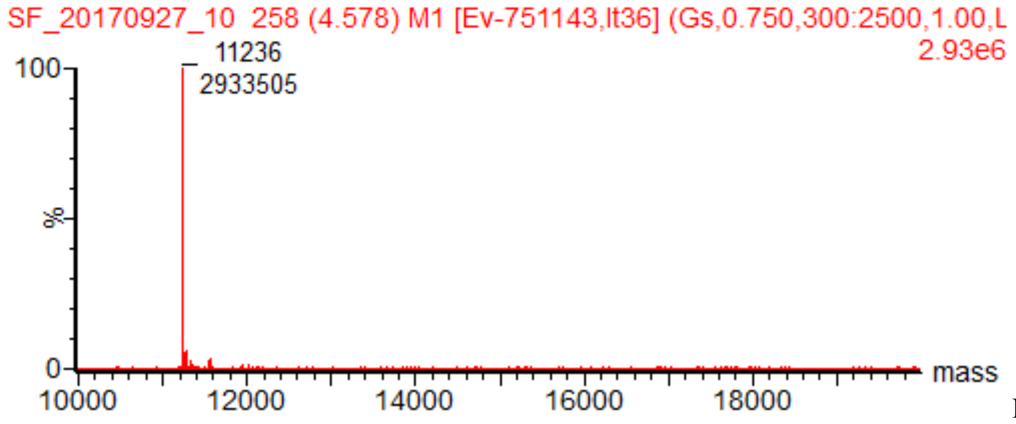


MS-ESI+:

Expected 11236.15 Da ; Found 11236 Da.

(Note that M from start codon is not present in final protein)

**Appendix Fig. 36:** Characterization of H4 after Hitrap purification- LCMS-ESI+

**Yield= 127 mg (53 mg/ L of culture)**

#### Additional data for 186 bp biotinylated DNA preparation (Appendix Figs. 36-37)

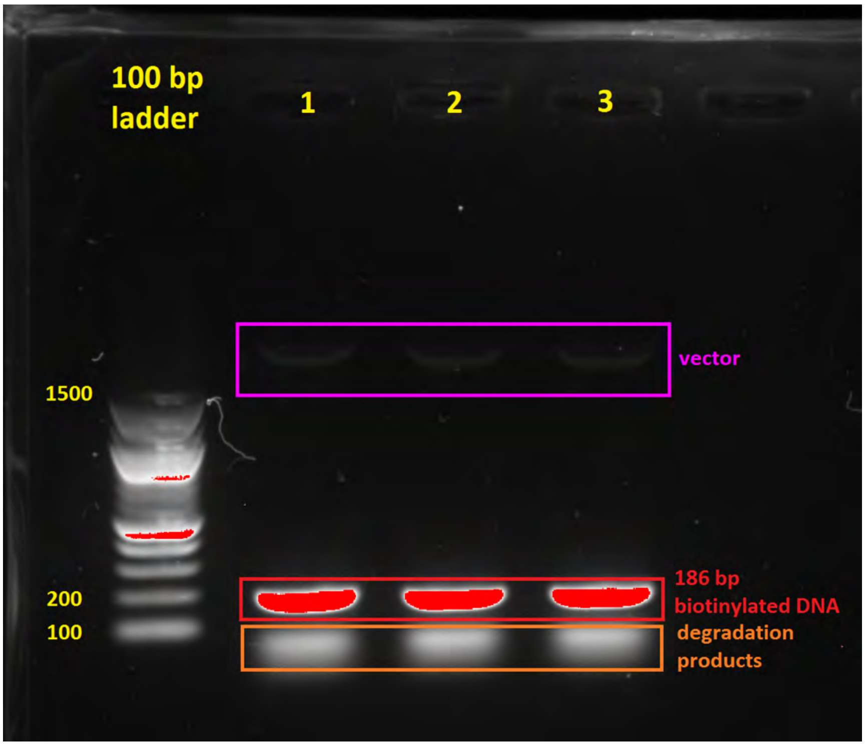


**Appendix Fig. 37:** Agarose gel electrophoresis analysis of three aliquots of 186 bp DNA after biotinylation reaction and freeze-thawing (stored at -20 °C). Some degradation was apparent, necessitating higher DNA: octamer ratios used in assembly.

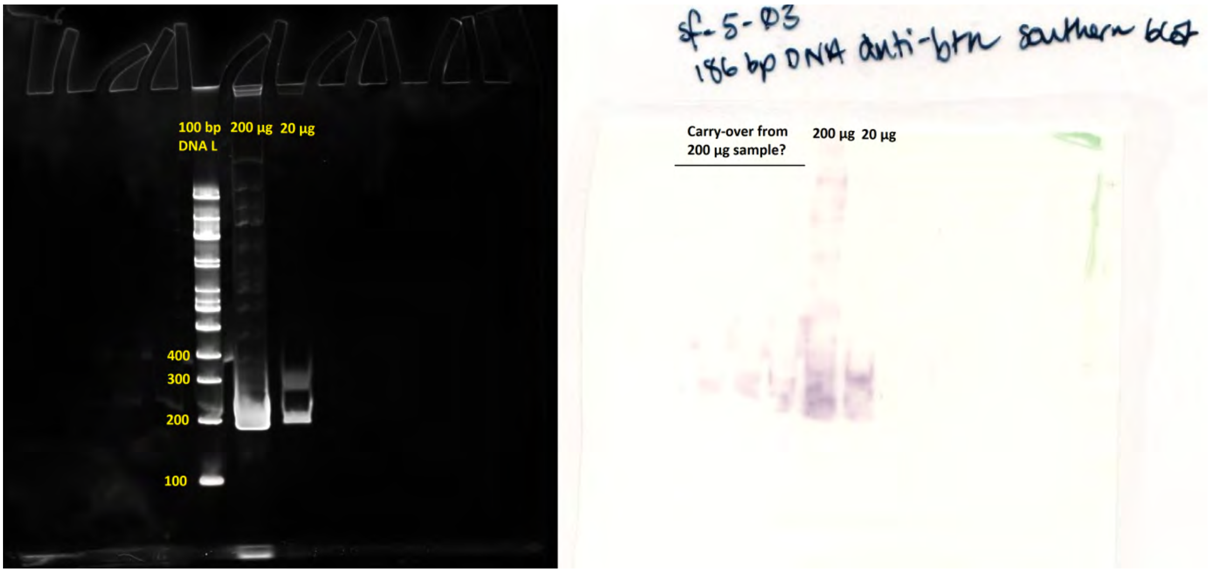


**Appendix Fig. 38:** Anti-biotin Southern Blot of 186 bp biotinylated DNA (uncropped gels).

**Additional data for MRG15 purification (Appendix Figs. 39-42)**

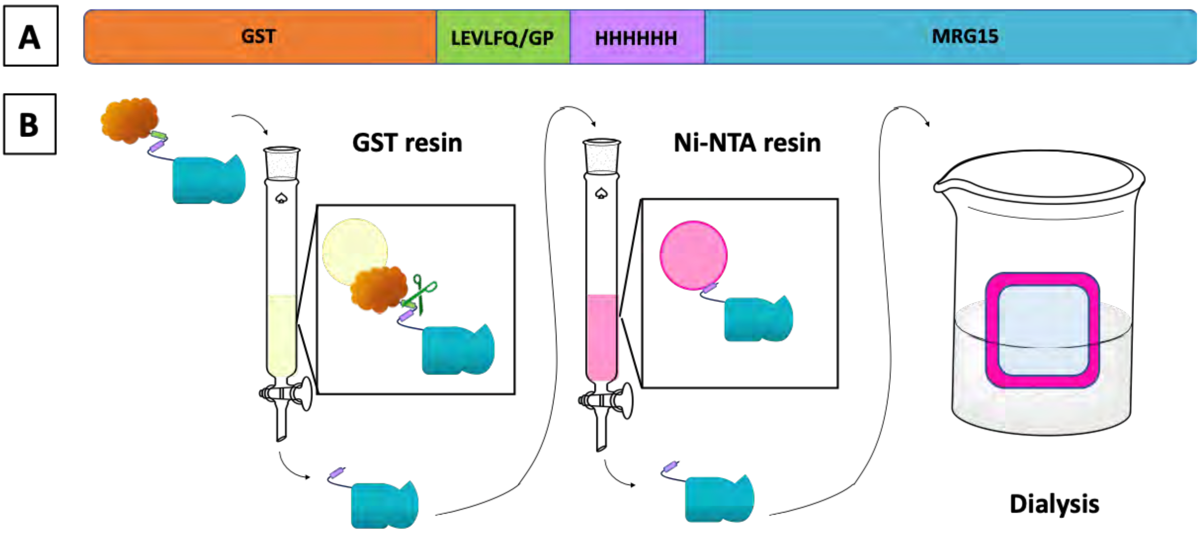


**Appendix Fig. 39:** (A) Schematic representation of the GST-MRG15 construct expressed, containing a N-terminal GST tag (orange), a Prescission Protease cleavage site (green), a $His_6$ tag (purple) and MRG15 (blue). (B). Schematic of the MRG15 purification protocol.

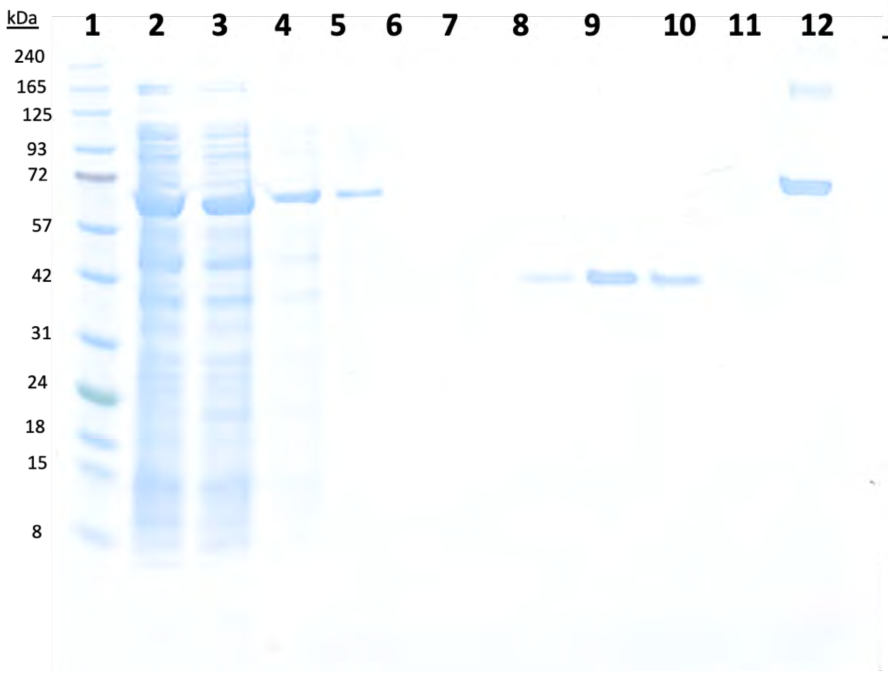


| | *Sample name* |
|---|---|
| *1* | Ladder |
| *2* | Pre-induction |
| *3* | Post-induction |
| *4* | GST beads 1 |
| *5* | GST beads 2 |
| *6* | p30 FT |
| *7* | E1 |
| *8* | E2 |
| *9* | E3 |
| *10* | E4 |
| *11* | p500 wash |
| *12* | BSA |

**Appendix Fig. 40:** SDS-PAGE analysis of MRG15 WT purification (4-12% Bis-Tris gel, MES buffer); Ladder= BluEye prestained protein standard. Pre-induction= sample taken from culture before IPTG added. Post-induction= sample taken after IPTG added. GST beads 1 and 2= samples of GST-MRG15 bound to GSH sepharose beads. P30 FT= flowthrough from wash with p30 buffer. E1,2,3 and 4= elution with p500 rounds 1, 2, 3 and 4. P500 wash= final wash with p500 buffer. BSA= positive loading control (1μg)

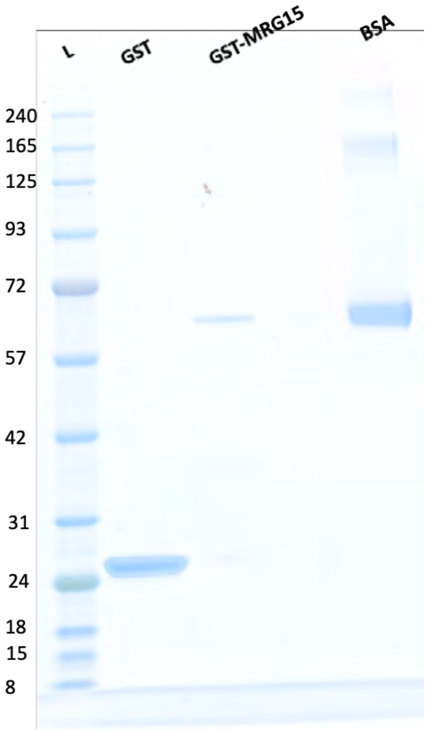


**Appendix Fig. 41:** GST and GST-MRG15 on GSH Sepharose beads. 1μg BSA used as loading control.

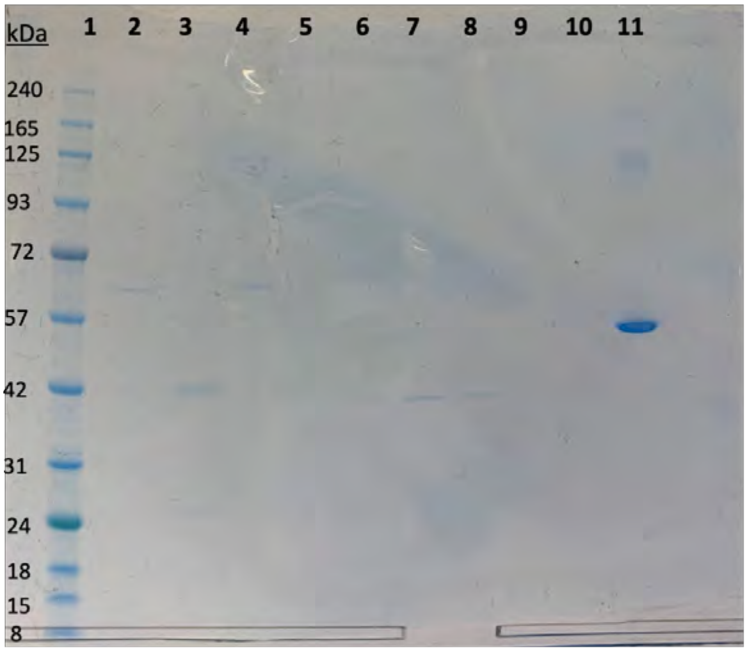


| | *Sample Name* |
|---|---|
| 1 | Ladder |
| 2 | wash |
| 3 | GST-MRG15 beads |
| 4 | GST beads |
| 5 | E1 |
| 6 | E2 |
| 7 | E3 |
| 8 | E4 |
| 9 | E5 |
| 10 | E6 |
| 11 | BSA 1 μg |

**Appendix Fig. 42:** SDS-PAGE analysis of MRG15 W53A purification (4-12% Bis-Tris gel, MES buffer); Ladder= BluEye prestained protein standard. E1- E6= elution rounds 1-6.
